# Impact on backpropagation of the spatial heterogeneity of sodium channel kinetics in the axon initial segment

**DOI:** 10.1101/2022.04.01.486760

**Authors:** Benjamin S.M. Barlow, André Longtin, Béla Joós

## Abstract

In a variety of neurons, action potentials (APs) initiate at the proximal axon, within a region called the axon initial segment (AIS), which has a high density of voltage-gated sodium channels (Na_V_s) on its membrane. In pyramidal neurons, the proximal AIS has been reported to exhibit a higher proportion of Na_V_s with gating properties that are “*right-shift*ed” to more depolarized voltages, compared to the distal AIS. Further, recent experiments have revealed that as neurons develop, the spatial distribution of Na_V_ subtypes along the AIS can change substantially, suggesting that neurons tune their excitability by modifying said distribution. When neurons are stimulated axonally, computational modelling has shown that this spatial separation of gating properties in the AIS enhances the backpropagation of APs into the dendrites. In contrast, in the more natural scenario of somatic stimulation, our simulations show that the same distribution can impede backpropagation. We implemented a range of hypothetical Na_V_ distributions in the AIS of three multicompartmental pyramidal cell models and investigated the precise kinetic mechanisms underlying such effects, as the spatial distribution of Na_V_ subtypes is varied. With axonal stimulation, proximal Na_V_ *availability* dominates, such that concentrating *right-shift*ed Na_V_s in the proximal AIS promotes backpropagation. However, with somatic stimulation, the models are insensitive to *availability*. Instead, the higher *activation* threshold of *right-shift*ed Na_V_s in the AIS impedes backpropagation. Therefore, recently observed developmental changes to the spatial separation and relative proportions of Na_V_1.2 and Na_V_1.6 in the AIS differentially impact *activation* and *availability*. The effects on backpropagation, and potentially learning, are opposite for orthodromic versus antidromic stimulation.

**Author Summary:** Neurons use sodium ion currents, controlled by a neuron’s voltage, to trigger signals called action potentials (APs). These APs typically result from synaptic input from other neurons onto the dendrites and soma. An AP is generated at the axon initial segment (AIS) just beyond the soma. From there, it travels down the axon to other cells, but can also propagate “backwards” towards the soma and dendrites. This “backpropagation” allows a comparison at synapses of the timing of outgoing and incoming signals, a feedback process that modifies synaptic connection strengths linked to learning. It is puzzling that in many neurons, sodium ion channels come in two types: high-voltage threshold channels clustered near the soma where the AIS begins, and low-voltage ones further away towards the axon. This separation changes in the early development of the animal, which raises the question of its role in backpropagation. We constructed a detailed mathematical model to explore how separation affects backpropagation. Separation either impedes or enhances learning, depending on whether the AP results from synaptic inputs or, less typically, currents moving backwards from the axon. This is explained by the different effects the separation has on two key kinetic processes that govern sodium currents.

## 1 Introduction

In fluorescence microscopy images of neurons, the axon initial segment (AIS) is visible as a patch of axonal membrane near the soma with a high density of voltage-gated ion channels. These channels enable the AIS to initiate and shape action potentials (spikes) and regulate neuronal excitability [1]. The AIS can be thought of as an organelle, that lives within the first ≈ 100μm of axonal membrane and whose function is to supply the current needed to initiate spikes when the neuron decides to fire —usually in response to synaptic input. The AIS can move, up and down the axon and also change its length on a timescale of hours to days. This phenomenon, called structural AIS plasticity, enables neurons to optimize their sensitivity to specific input frequencies during development and to homeostatically adjust their intrinsic excitability [2, 3, 4]. GABAergic input can also impinge on the AIS from axo-axonic synapses, such that the AIS can be modulated directly by interneurons. Synaptic input at the AIS can rapidly and precisely control the excitability of individual neurons for sound localization [5]. Fast AIS plasticity, including receptor-mediated changes to local ion channel properties and endocytosis of voltage-gated channels, occurs on timescales of seconds to minutes [6]. (This is distinct from pathological remodelling induced by ischemia, although in [7], it was recently demonstrated that cortical neurons are more robust to interruptions in blood flow than previously thought.) The outsized electrophysiological influence of the AIS demands robust characterization of this short piece of axon as it interacts with its environment.

Over three-quarters of all neurons in the mammalian cortex are pyramidal cells (see Figure S1), which have dendrites spanning the thickness of the cortex (several mm) and AIS lengths on the order of tens of μm [8, 9, 10]. The AIS requires a high density of voltage-gated sodium channels (Na_V_s) to prime and initiate action potentials [11, 12, 13]. In pyramidal cells, the AIS features two Na_V_ subtypes, with an interesting spatial distribution: Na_V_1.2 channels cluster near the soma (i.e. at the proximal AIS) while Na_V_1.6 cluster toward the distal AIS [14, 15, 16]. However, the purpose of this separated distribution of Na_V_ subtypes remains unclear [17, 18]. Further, recent experiments have revealed that as neurons develop, the spatial distribution of Na_V_s in the AIS can change substantially, suggesting that neurons tune their excitability by modifying said distribution [19].

It is a prevalent view that the Hodgkin-Huxley kinetics of Na_V_1.2 are *right-shift*ed relative to those of Na_V_1.6 by an amount *V*_RS_ ∼ 10 − 15mV [20, 14]. The separated Na_V_ distribution is reported to promote backpropagation —which is important for learning— following axonal stimulation [14]. There is also evidence that mutations which alter the gating properties of Na_V_1.2 are involved in epilepsy and autism [21]. Backpropagated spikes drive learning by depolarizing the postsynaptic membrane, which triggers metabolic events that give rise to synaptic plasticity, including spike-timing-dependent plasticity [22]. There is experimental evidence that postsynaptic backpropagation can release retrograde messengers into the synapse, and influence the future release of neurotransmitters from the presynaptic neuron [23].

A backpropagating action potential (BAP) can underlie bursting in cortical neurons as it can return to the cell body from the dendrites as a depolarizing after potential, which in turn can initiate another somatic AP [24, 25]. Bursting can also occur in layer 5 pyramidal cells following the generation of a dendritic BAP-activated Ca^2+^ spike (BAC spike), e.g. in the presence of synaptic input. The associated BAPs can further influence the dendritic dynamics [26, 27, 28, 29].

Not all layer 5 pyramidal cells can generate dendritic spikes as the size of the apical dendritic tree varies [30]. Dendritic spikes have also been reported to vary across species, and are not common in human layer 5 pyramidal cells [31], owing partly to their enhanced dendritic compartmentalization [32]. Thus, to validate and further understand the original reports that Na_V_ segregation promotes BAPs, we investigate how Na_V_ segregation in the AIS can promote BAPs using the model of [14] (itself based on [24]). This provides the backbone to study the basic effects on BAPs of the AIS excitability profile, under both somatic and axonal stimulation. For the sake of generality, we complement these results by considering a state-of-the-art model of layer 5 pyramidal cells with perisomatic BAPs and dendritic BAC firing [27], adapted to include a more realistic AIS and axon.

Other computational powers are attributed to the AIS. Moving the initiation site away from the soma (i.e. toward the distal AIS) beyond a critical distance enables high-frequency spiking in cortical neurons, increasing the maximum spike frequency by an order of magnitude [13]. Separating Na_V_1.6 into the distal AIS is said to push the initiation site toward that location, owing to those channels’ lower voltage threshold [14, 20]. However, in [33], simulations having only one Na_V_ type demonstrated that passive cable properties are sufficient to locate AP initiation at the distal AIS.

In our simulations, we alter the composition of the AIS and look for changes in the backpropagation threshold. The model neurons are allowed to distribute *right-shift*ed Na_V_ gating properties along the AIS, by differentially distributing two functionally distinct classes of sodium channels, referred to here as Na_V_1.2 and Na_V_1.6 following [14], [20] and [19]. Our modelling study is motivated by the following question: what effect does the separated spatial distribution of Na_V_1.2 (or *right-shift*ed gating properties) and Na_V_1.6 (*left-shift*) in the AIS have on excitability and backpropagation? In particular, how does the finding in [14], that the separated distribution of Na_V_ subtypes favours backpropagation, generalize to the more common situation of somatic stimulation?

A natural approach is to systematically alter that Na_V_ distribution, by varying the extent to which Na_V_ subtypes are spatially segregated in the AIS without affecting the total Na_V_ density. We compute the threshold for backpropagation as the amplitude of a brief current pulse that causes an AP to propagate back into the dendrites and cause a sufficient depolarization (Section S1.2, Section S1.3). This is done in three biophysically detailed and independently tuned multicompartmental pyramidal cell models ([14], [27]), two of which are based on the Hu et al. 2009 model and involve the same morphology but with differing soma-dendrite excitability balance (cell geometries are provided in the Supplementary Information: Figure S1 and Figure S2). This threshold is computed as a function of the spatial segregation of the Na_V_ subtypes in the AIS by continuously varying their density profiles from fully overlapping to strongly separated (Figure 1).

**Figure 1:**
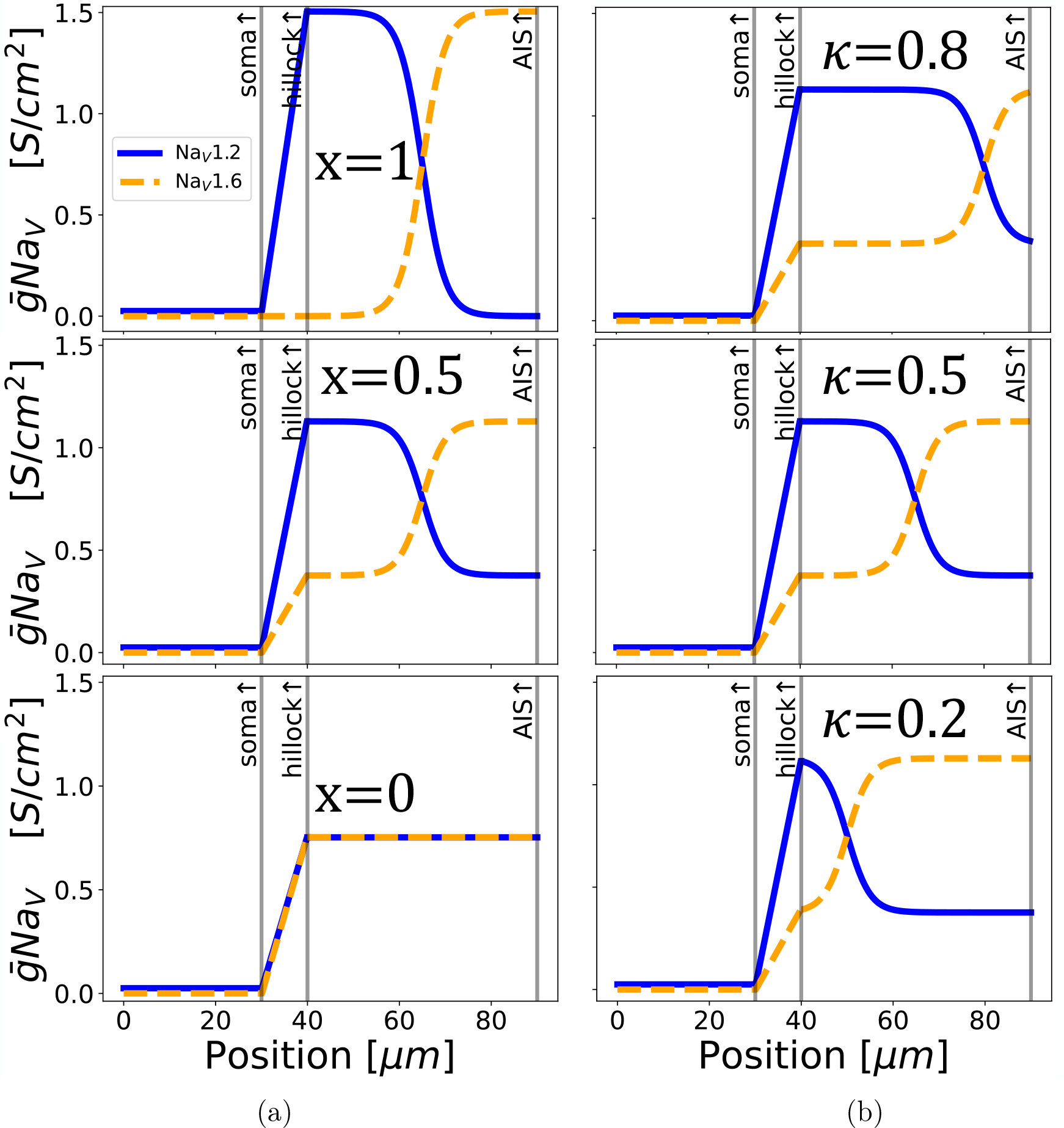
Modifying the spatial distribution of Na_V_ subtypes in the AIS while keeping the total conductance constant. **(a)** The spatial separation of Na_V_ subtypes in the AIS is varied using the parameter “*x*” with *κ* = 0.5. The top plot is a model setup with a separated distribution [14] of Na_V_s in the AIS. The high threshold Na_V_1.2 (indicated in blue) are concentrated close to the soma, and the low threshold Na_V_1.6 (indicated in orange) are kept distal to the soma. Moving from top to bottom, both Na_V_ subtypes are distributed ever more evenly along the AIS. We chose the parameter name “*x*” to vary the spatial separation of the AIS Na_V_ distributions, because the separated distribution is *x*-shaped. Setting *x* = 1 in our simulations gives the separated distribution, and *x* = 0 gives the “flat” distribution wherein both Na_V_ subtypes are uniformly mixed. **(b)** Variation of the crossover location (*κ*) of Na_V_s in the AIS with *x* = 0.5. We have lengthened the AIS to 50μm in this graphic for visual clarity.

We show that Na_V_ separation enhances backpropagation with axonal stimulation but can impede it with somatic stimulation. This asymmetrical result was not expected. To explain our results, we independently modify the *right-shift* (*V*_RS_) of selected Na_V_1.2 gating variables and their respective time constants by an amount Δ*V*_RS_ (i.e. *V*_RS_ ⟶ *V*_RS_ + Δ*V*_RS_). These modifications to Na_V_1.2 gating are applied only in the AIS.

Sweeping Δ*V*_RS_ (while clamping other gating variables to nominal *V*_RS_ values) reveals that (I) Na_V_1.2 *availability* and its time constant explain how proximal Na_V_1.2 promotes backpropagation with axonal stimulation, and (II) the threshold of steady-state *activation* explains how Na_V_1.2 suppresses backpropagation and reduces excitability with somatic stimulation.

Being a feature of pyramidal cells, the plastic distribution of AIS Na_V_ subtypes that we model applies to something like eight out of ten cortical neurons. We have demonstrated opposing effects on backpropagation with orthodromic versus antidromic stimulation by altering said Na_V_ distribution. Various experimental and computational techniques used to study the biophysical determinants of AIS excitability across the lifespan have involved different stimulation sites. It is thus important to know whether and how the spatial profile of Na_V_ channel subtypes really enhances backpropagation in vivo, and whether moving the stimulating electrode can bias or even invert experimental findings, as our work demonstrates. Apart from explaining the dynamical mechanism behind the dependence of AP generation on AIS Na_V_ distribution, we clearly show that the site of stimulation matters, a finding that is present robustly in different models and which merits experimental confirmation. Changes to AIS properties and the follow-on effects on backpropagation must affect the entire cortex.

## 2 Results

### 2.1 Hypothetical Na_V_ distributions in the AIS

We begin with our implementation of the model from Hu et al. 2009 [14] (Hu-based model), using their morphology, K_V_ and Na_V_ kinetics. The standard AIS length in our model is 25 μm, based on measurements from [10]. A key feature of the Na_V_ distribution that changes during development, is the extent to which the voltage-gated sodium channel subtypes Na_V_1.2 and Na_V_1.6 are localized in the proximal and distal AIS, respectively [19]. In these simulations, the relative proportion of Na_V_1.2 versus Na_V_1.6 at a given position along the AIS can be changed without affecting the total Na_V_ density at any point (Equation 5).

Figure 1 shows how the parameters *x* and *κ* control the way Na_V_ subtypes are spread out along the AIS. When *x* is at its highest value of 1, the subtypes Na_V_1.2 and Na_V_1.6 are spaced apart from each other, with Na_V_1.2 concentrated in the proximal AIS and Na_V_1.6 in the distal AIS, approximating the distribution observed in developing pyramidal neurons (see [19]). Decreasing *x* transforms this separated distribution into a uniform mix (*x* ⟶ 0) where Na_V_1.2 and Na_V_1.6 are distributed homogeneously. This can be seen in Figure 1a.

Every distribution except the uniform Na_V_ mix has a location along the AIS at which the density of Na_V_1.6 overtakes the Na_V_1.2 density. That location, which we call the Na_V_ crossover and denote *κ*, is also varied in our simulations (see Figure 1b; *κ* is a dimensionless length normalized by the AIS length.).

To cement our results, we will further apply identical transformations to the Na_V_ distribution in a cell having a ‘backward’ AIS, that is, with *distal* Na_V_1.2 and *proximal* Na_V_1.6. The results from the backward AIS model are nearly a mirror image of our findings.

For each hypothetical Na_V_ distribution, a short current pulse (1ms) is injected at a specific site, and the minimum (i.e. threshold) pulse amplitude *I* (in nA) required to elicit a spike is determined. Brief pulse durations separate the stimulation waveform from the intrinsic response of the cell. We define excitability in terms of two thresholds: backpropagation threshold *I*_*BP*_ (AP leading to a spike in the distal dendrites) and forward-propagation threshold *I*_*FP*_ (axonal AP threshold, recorded without regard to the amplitude of the somatodendritic depolarization).

Current is injected either in the middle of the soma (somatic stimulation) or the axon just distal to the AIS (axonal stimulation). Both are used by experimentalists [34], and certain pyramidal neurons are also known to receive axo-axonic input at the AIS as well as somatodendritic input [6]. In both cases, forward propagation refers to an AP travelling down the axon, and backpropagation always refers to an AP visible as a spike in the dendrites. Backpropagation was deemed to have occurred if all apical dendritic tips exceeded −63.0mV (i.e. a depolarization of 7.0mV above *V*_rest_) following stimulation (see Supplementary Information, Figure S3).

In the following sections, we implement the above Na_V_ distributions in the Hu model [14]. Later, in Section 2.6, we repeat this procedure in the model of Hay et al. [27]. The third model, an alternate tuning of the Hu model with significant qualitative differences in its backpropagating action potential, is included in the supplementary information (Section S1.4).

### 2.2 Somatic stimulation

In Figure 2, both negatively and positively sloped backpropagation threshold curves with respect to *x* are present, indicating that Na_V_ separation can promote or impede backpropagation (respectively). Changes in threshold can be as large as 30%. Moving the Na_V_ crossover (*κ*) toward the distal AIS shifts the backpropagation threshold curves upward. A qualitative change, namely the sign of the slope, occurs around *κ* = 0.4.

**Figure 2:**
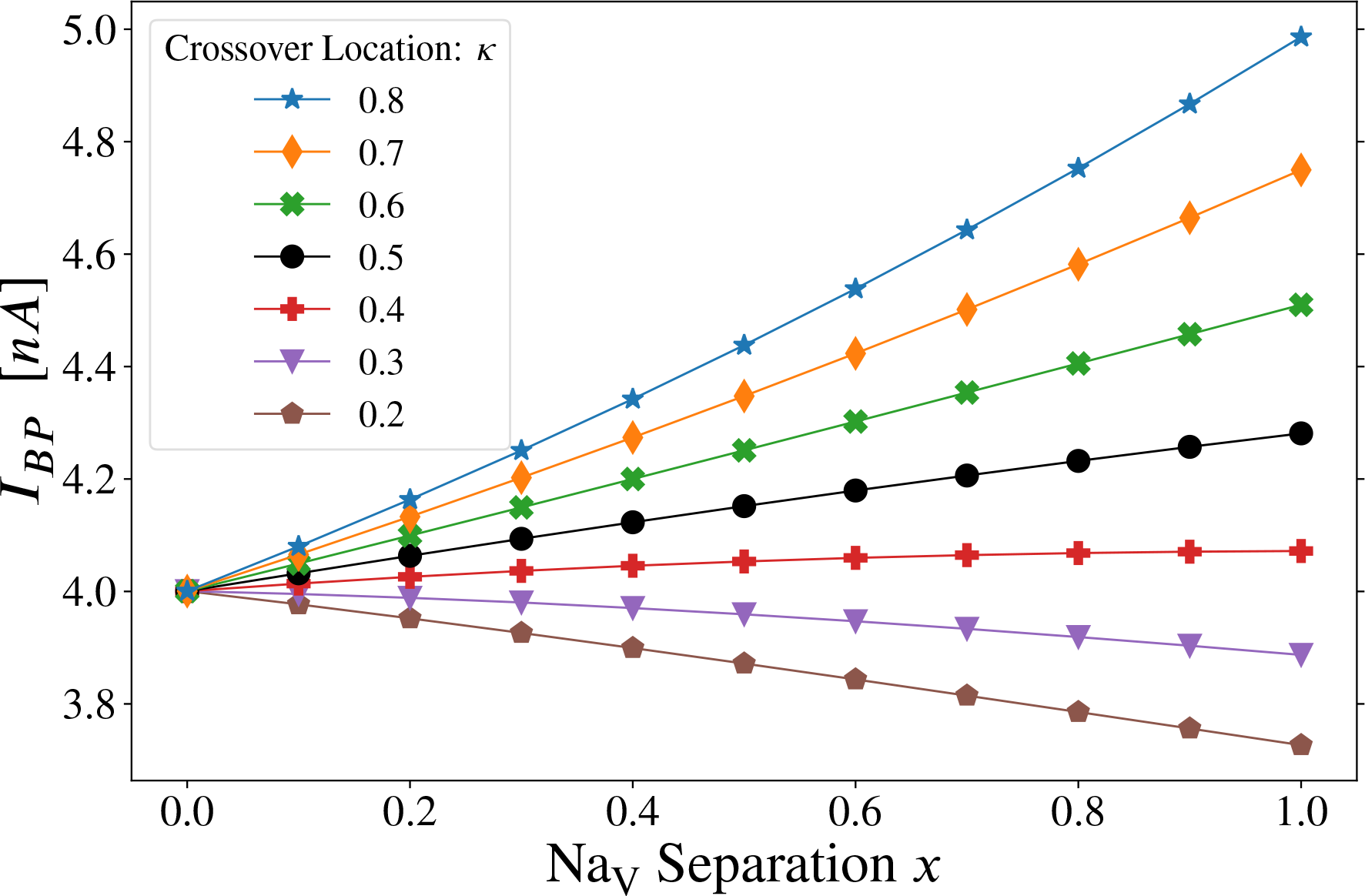
Somatic stimulation: combined effect of varying crossover location (*κ*) and Na_V_ separation (*x*) in the axon initial segment. The threshold for forward AP propagation is the same as for backpropagation. Varying the separation parameter “*x*” from *x* = 0 to *x* = 1, the distribution of Na_V_ channels goes from flat (homogeneous) to separated, the latter approximating the distribution observed in developing pyramidal neurons (see Figure 1a). Note that curves for all values of *κ* converge to a single point at *x* = 0 since *κ* can have no effect when the two Na_V_ subtypes are uniformly distributed along the AIS. The lines have been drawn to guide the eye.

An intuitive explanation for this latter effect is that moving the crossover location away from the soma causes the AIS to be dominated by Na_V_1.2 channels (see Figure 1b, *κ* = 0.8), which have a higher voltage threshold than Na_V_1.6. APs still initiate in the distal AIS, but the dominant Na_V_1.2 renders the cell less excitable. Further, for *κ* ≳ 0.4, the backpropagation threshold increases as we tend toward the separated, *x*-shaped distribution of Na_V_s. This behaviour is the opposite of what is observed for axonal stimulation below and in [14]. We repeated these simulations with AIS length up to 100μm (Figure S11) and also with stimulation at the main apical dendrite (Figure S13) instead of the soma, and obtained the same qualitative results as Figure 2 (see Supplementary Information Section S1.4.1).

The negatively sloped curves do not necessarily imply that proximal Na_V_1.2 promotes backpropagation in the case of somatic stimulation. In those curves (*κ* ≲ 0.4), the AIS is mainly populated with Na_V_1.6 when *x* > 0. Also note that decreasing *κ* places more Na_V_1.6 channels nearer to the soma (see Figure 1b,

*κ* = 0.2). In that case, the threshold-lowering effect of Na_V_ separation could come from the increased total Na_V_1.6 density that results from increasing *x* when *κ* is relatively small, rather than from the proximal accumulation of Na_V_1.2 with increasing *x*. Further, increasing *κ* (which increases the ratio of Na_V_1.2 to Na_V_1.6 in the AIS) raises the threshold for all curves in Figure 2 (see also Figure S12). It is then consistent to postulate that for somatic stimulation, the backpropagation threshold is increased by AIS Na_V_1.2 at all values of *x* and *κ*, and Figure 2 is consistent with AIS Na_V_1.6 enhancing excitability and backpropagation. In other words, for **somatic stimulation**:

- when *κ* < 0.5 and *x* > 0, the AIS is dominated by Na_V_1.6: increasing *x* increases the proportion of total AIS Na_V_ conductance due to Na_V_1.6 (negative slope: separated distribution yields the lowest backpropagation threshold).
- when *κ* > 0.5 and *x* > 0, the AIS is dominated by Na_V_1.2: increasing *x* decreases the proportion of total AIS Na_V_ conductance due to Na_V_1.6 (positive slope: separated distribution yields the highest backpropagation threshold).

Lengthening the hillock with *κ* fixed also moves the crossover away from the soma. Curves with negative slope in Figure 2 became positively sloped when the hillock was lengthened from 10μm to 30μm (Figure S14). The forward propagation threshold for somatic stimulation with a single 1ms current pulse is not included in a separate figure since it is identical to the backpropagation threshold in this model. This does not depend on the somatic injection site. The effect of Na_V_ gating properties in the AIS on backpropagation threshold is examined systematically in Section 2.5, Figure 6.

An informative variation on Figure 2 is shown in Figure 3c in which the AIS is “put on backward”, such that Na_V_1.2 is concentrated in the distal AIS and Na_V_1.6 is proximal to the soma. As one might expect, the effect of varying *x* and *κ* in Figure 3 is opposite to what is seen in Figure 2, albeit with some new curvature at low *κ*. This reinforces the observation that proximal Na_V_1.6 facilitates backpropagation with somatic stimulation.

**Figure 3:**
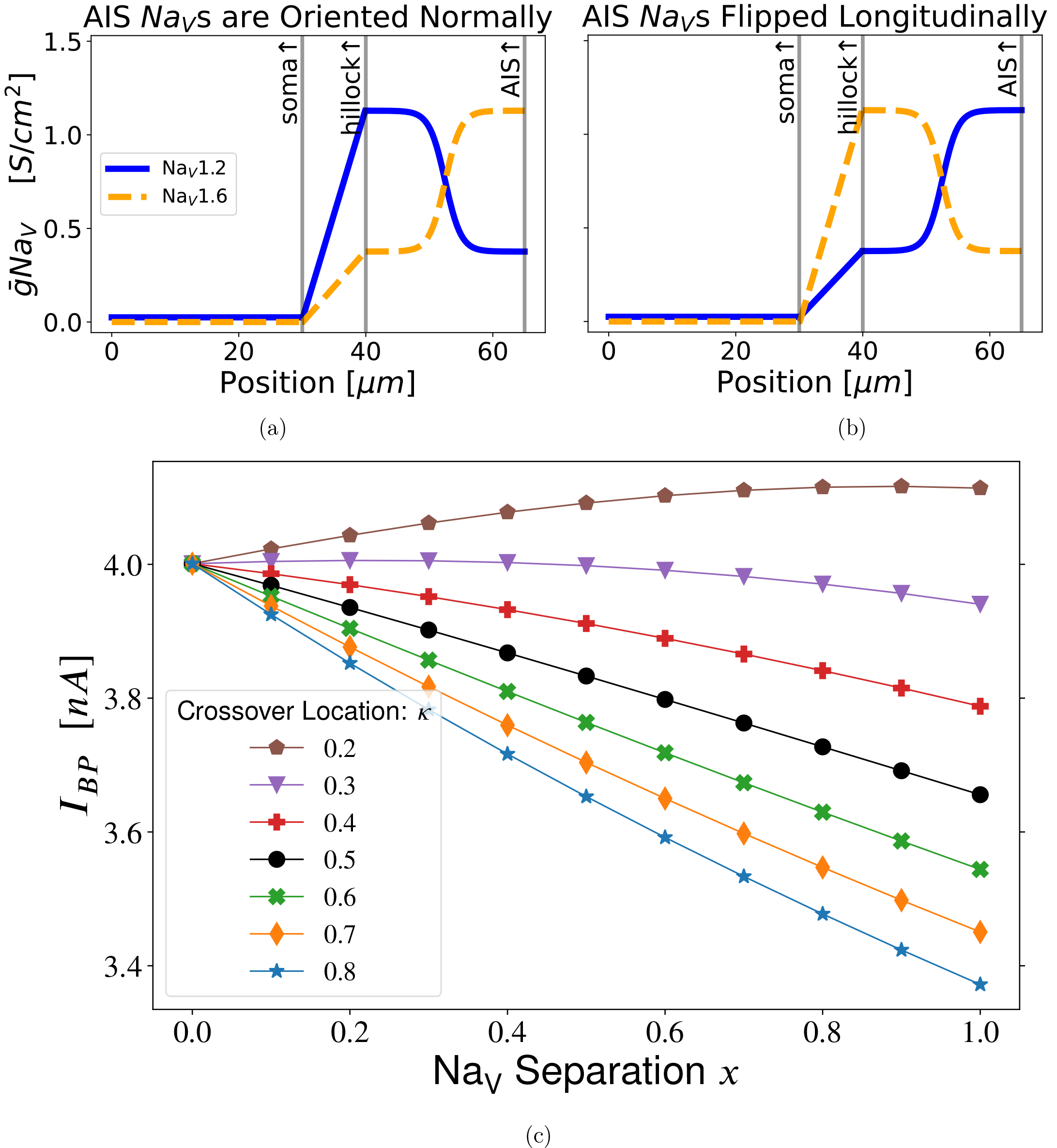
Somatic stimulation with a flipped Na_V_ distribution: backward AIS. When the AIS Na_V_ distribution is flipped proximal-to-distal, setting *x* = 1 concentrates Na_V_1.6 at the proximal AIS and Na_V_1.2 at the distal AIS —the opposite of what is observed in many pyramidal cells [14, 15, 16]. **(a)** AIS with proper longitudinal placement of Na_V_s. **(b)** AIS with a longitudinally flipped Na_V_ distribution. In both plots, *x* = 0.5 and *κ* = 0.5. **(c)** Somatic stimulation with AIS Na_V_s flipped as in **(b)**: This result is close to a mirror image of Figure 2. The lines have been drawn to guide the eye.

### 2.3 Axonal stimulation

With axonal stimulation (current injection just distal to the AIS), Na_V_ separation consistently lowers the backpropagation threshold (Figure 4). Contrary to somatic stimulation (Figure 2), moving the Na_V_ crossover (*κ*) toward the distal AIS shifts the backpropagation threshold curves downward.

**Figure 4:**
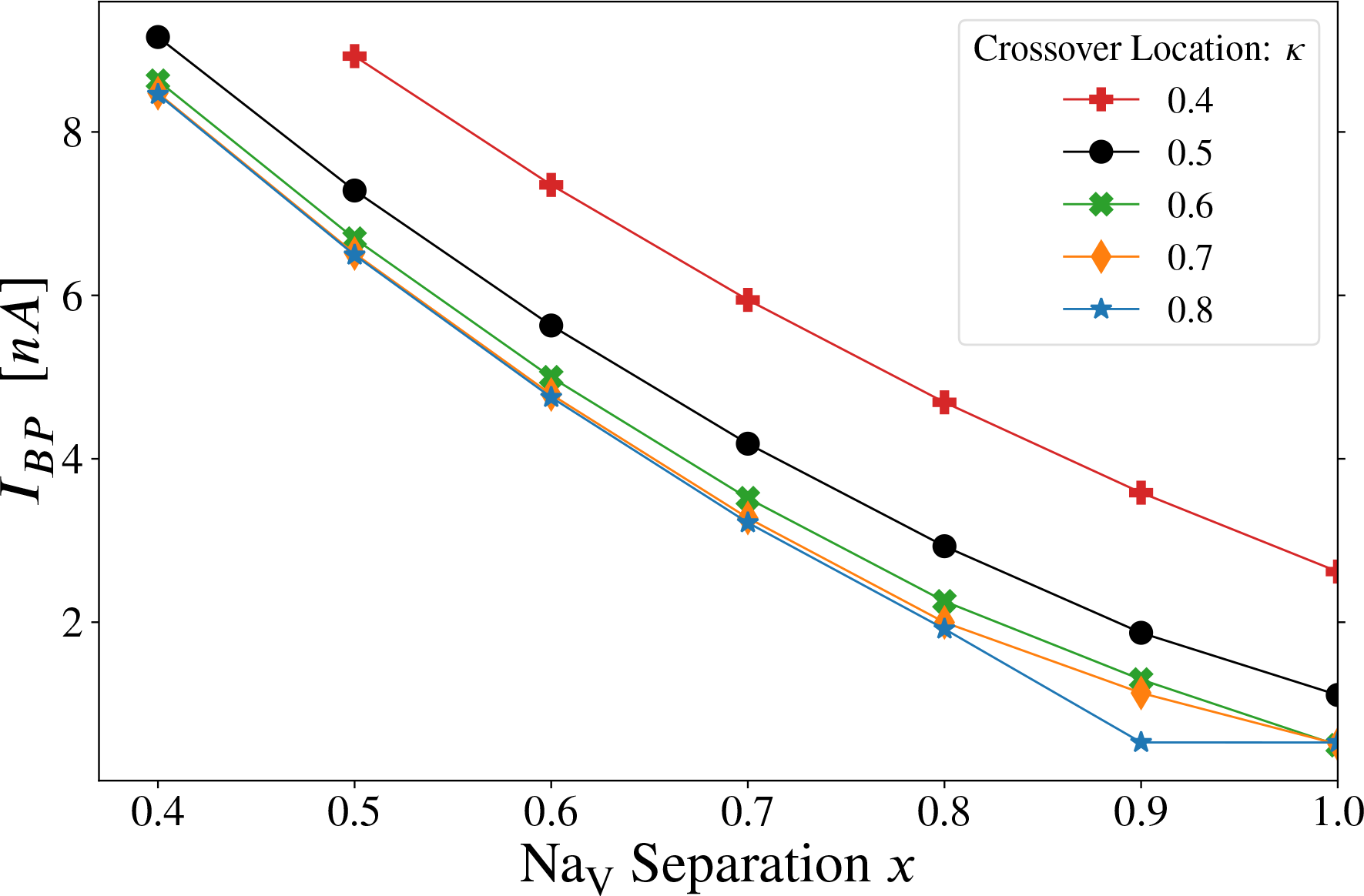
Axonal stimulation: effect of varying crossover location (*κ*) and Na_V_ separation (*x*) in the AIS on the backpropagation threshold (see Figure 1). When computing the threshold, the stimulating current was limited to a maximum of 10nA, to prevent unphysiological local depolarization at the stimulation site. Due to the smaller diameter of the axon (relative to the soma), 10nA is sufficient to depolarize the membrane potential to ≈ +80mV at the stimulation site, whereas the resting potential is *V*_rest_ = −70mV. To achieve backpropagation within that constraint (following axonal stimulation), our model required some amount of proximal Na_V_1.2, delivered through the combined effects of Na_V_ separation (*x* ≳ 0.5) and a sufficiently distal crossover position *κ* ≳ 0.4. Separating the two Na_V_ subtypes (*x* ⟶ 1) lowers the threshold, in agreement with the finding in [14] that proximal accumulation of Na_V_1.2 promotes backpropagation, albeit due to different gating properties (Figure 6b). Increasing *κ* raises the proportion of Na_V_1.2 (relative to Na_V_1.6) in the AIS and lowers the backpropagation threshold as well. Threshold changes here are larger than for somatic stimulation (Figure 2). The lines have been drawn to guide the eye.

The decreasing threshold with respect to *x* in Figure 4 is consistent with the conclusion from [14], which used axonal stimulation, that proximal Na_V_1.2 in the AIS promotes backpropagation. Our results for *κ*, with axonal stimulation, provide new support for their findings.

This agreement is interesting because our method of modifying the AIS Na_V_ distribution (described above in Figure 1) is quite different from their simulations. Our transformations deliberately preserve the total Na_V_ density at every AIS segment —if Na_V_1.2 is removed, Na_V_1.6 must take its place. Conversely, in [14], the density profile of Na_V_1.2 is scaled by a constant factor everywhere in the AIS, leaving the Na_V_1.6 profile intact. We denote the scaling factor

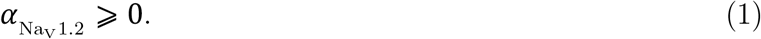

That is, if the Na_V_1.2 density profile is scaled down in [14], nothing is added to compensate for the missing channels. Under the latter transformation, we expect that 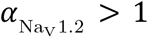 would lower *I*_*BP*_ and 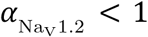 would raise *I*_*BP*_ in our models as well, since scaling the density profile of Na_V_1.2 in a separated distribution with a specified *κ* and *x* > 0 would scale the total AIS Na_V_ conductance, especially at the proximal AIS. We have reproduced this procedure in the Hay model, see Section 2.7.

It is one thing to say that reducing (increasing) the total density of voltage-gated sodium channels in the proximal AIS, which happen to be Na_V_1.2 channels, will raise (lower) the backpropagation threshold (respectively). But since we preserved the local Na_V_ density in our results (above), the changes to *I*_*BP*_ can only be a manifestation of the spatial heterogeneity of sodium channel *gating properties*. In Section 2.5, Figure 6, the *right-shift* (*V*_RS_) of Na_V_1.2 gating properties is modified in the AIS. Since *right-shift* is the most important feature distinguishing Na_V_1.2 from Na_V_1.6 in this model, the analysis in Figure 6 explains how the proximal accumulation of Na_V_1.2 is able to simultaneously lower *I*_*BP*_ with axonal stimulation (Figure 4) and raise *I*_*BP*_ with somatic stimulation (Figure 2) .

### 2.4 Forward propagation threshold

The forward-propagation threshold *I*_*FP*_, also referred to as the AP threshold, is shown in Figure 5 for the Hu-based model. With axonal stimulation only, it is possible to elicit an action potential without creating sufficient depolarization in the apical dendrites to meet our strict criterion for backpropagation (see Figure S4, Figure S3). Note, however, that the most distal dendrites depolarize to several mV above their local resting potential (see Figure S4b). Stimulation amplitude is an order of magnitude lower than in the case of *I*_*BP*_. This is expected with axonal stimulation due to the high Na_V_ density of the distal AIS, its electrical isolation from the soma, its proximity to the stimulus, and our stringent definition of *I*_*BP*_ (Section S1.2). (Further, as discussed in [35], Hu et al.[14] built their model on [24], in which somatic invasion of the axonal action potential is reduced.)

**Figure 5:**
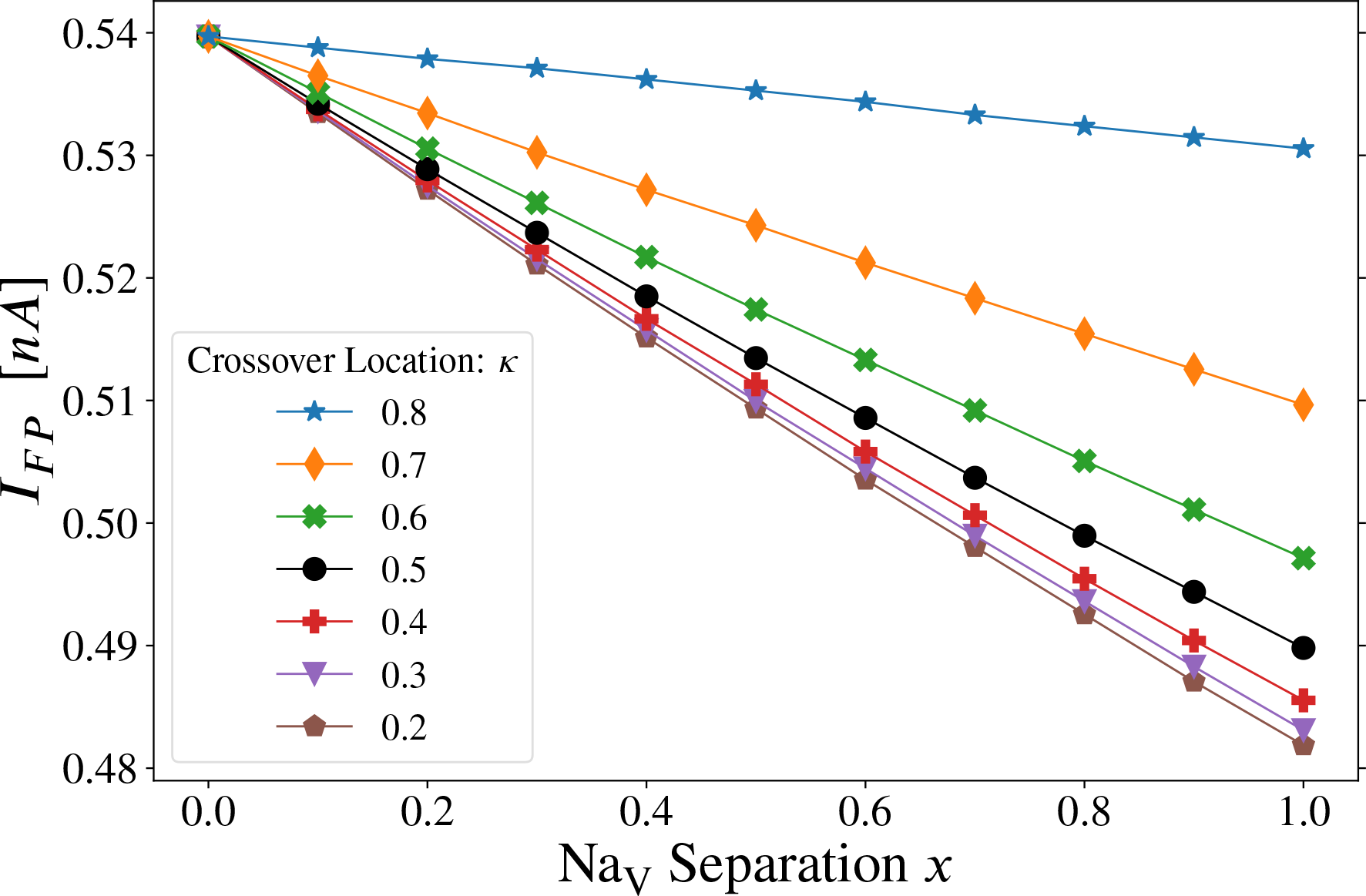
Axonal stimulation: effect of *x* and *κ* on forward propagation threshold. The trend for all constant *κ* curves is that raising the proportion of total AIS Na_V_1.6 (by reducing *κ*) or concentrating Na_V_1.6 in the distal AIS (by increasing *x*) lowers the threshold to initiate forward propagating action potentials (see Figure 1). Note that although this threshold current pulse is not sufficient to satisfy our strict backpropagation criterion (see Section S1.2), the most distal apical dendrites will be depolarized by several mV relative to their local resting potential (see Figure S4b). The lines have been drawn to guide the eye.

**Figure 6:**
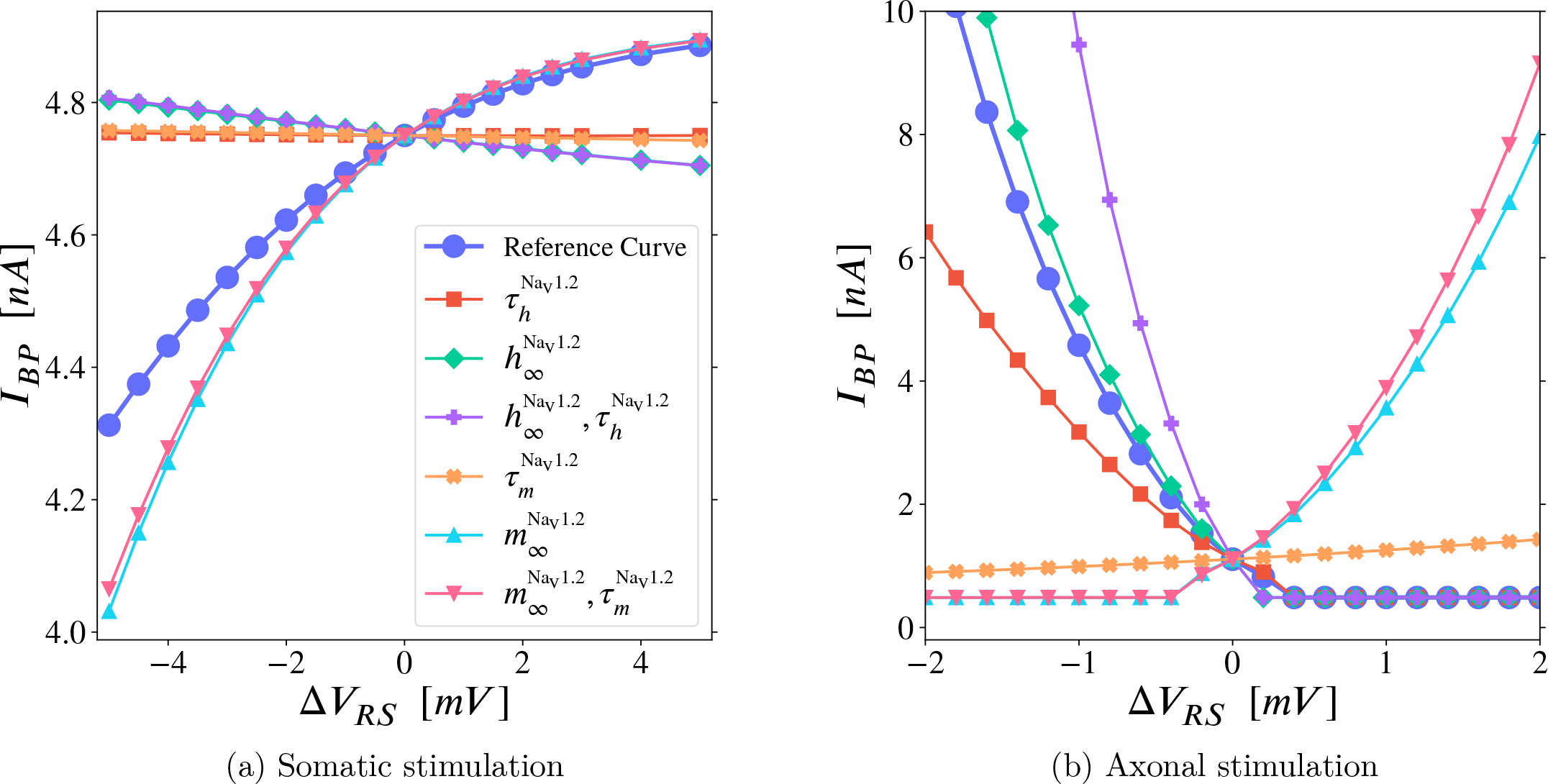
Sensitivity analysis of the backpropagation threshold to the *right-shift* of Na_V_1.2 gating properties. Along each curve, the gating properties named in the legend have their *right-shift* changed from *V*_RS_ to (*V*_RS_+Δ*V*_RS_),and the omitted gating properties are left unchanged (full definition and notation in Section S1.6.2). When Δ*V*_RS_ = 0, the *right-shift* is the reference value (or ‘nominal value’) of 13.0mV used for Na_V_1.2 in our simulations(see Section S1.6.2, *V*_RS_ indicated by small 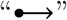 in Figure S15), around which we are performing this sensitivity analysis. The reference curve (legended 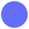) shows the net effect of *right-shift*ing all Na_V_1.2 properties on *I*_*BP*_, via its slope. (It may be useful to imagine points on the reference curve as being pulled toward all the other curves that only change one property. The reference curve would then be the result of the combined pulls of those curves.) For each mode of stimulation, we identify the key gating properties through which *right-shift* controls backpropagation, by comparing the single property curves (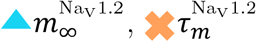, etc.) to the reference curve (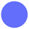). **(a) Somatic stimulation:** The reference curve has a positive slope (*right-shift* raises *I*_*BP*_), and it follows curves legended with 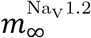 near the nominal point (i.e. near Δ*V*_RS_ = 0). Hence, *I*_*BP*_ is governed by Na_V_ steady-state *activation* and is insensitive to the *right-shift* of all Na_V_ time constants. **(b) Axonal stimulation:** The reference curve has a negative slope (*right-shift* lowers *I*_*BP*_, i.e. promotes backpropagation), and it follows curves legended with 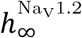 or 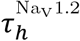. *I*_*BP*_ is then governed by proximal Na_V_ *availability*, owing to the *right-shift* of Na_V_1.2. Notably, with axonal stimulation, *I*_*BP*_ is also sensitive to the *right-shift* of 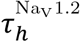 (*V*), the —voltage-sensitive— *availability* time constant. Results are summarized in **Table 1**. The lines have been drawn to guide the2eye. In both plots, *x* = 1.0. On the left *κ* = 0.7(to increase the slope, see Figure 2), and on the right, *κ* = 0.5.

As with *I*_*BP*_, increasing *x* lowers *I*_*FP*_. Na_V_ separation concentrates Na_V_1.6 in the distal AIS, making it more excitable in the portion nearer to the stimulation site. This finding is consistent with [14], who found that distal Na_V_1.6 density places the lowest initiation threshold (and therefore the AP trigger zone) at the distal AIS. However, [33] has shown that cable properties are sufficient to explain why the trigger zone is located at the distal AIS (see Discussion). Moving the crossover distally (*κ* ⟶ 1) increases the total proportion of Na_V_1.2 in the AIS and thereby raises the *I*_*FP*_ threshold due to *activation right-shift*.

### 2.5 Modifying the *right-shift* of selected gating properties in the AIS

Our results from varying the Na_V_ distribution may be counterintuitive. With axonal stimulation, con-centrating low threshold Na_V_1.6 channels at the distal AIS ought to promote forward propagation (and it does, see Figure 5), but why would concentrating the *high threshold* Na_V_1.2 channels at the proximal AIS facilitate backpropagation? And how does the asymmetry come about, such that separating Na_V_ sub-types can raise the backpropagation threshold with somatic stimulation, but always lowers it with axonal stimulation?

In this section, we perform a type of sensitivity analysis with respect to the effects of the *right-shift*ed Na_V_1.2 subtype. Figure 6 allows us to isolate the effects of *activation right-shift* versus *availability right-shift* on the backpropagation threshold.

Na_V_ subtypes are defined by their gating properties. Each Na_V_ distribution (Figure 1) produces a corresponding spatial profile of gating properties, including *right-shift*. (Gating properties are detailed in Supplementary Information, see Figure S15.) In Figure 6, the AIS Na_V_ distribution is fixed at *x* = 1. This spatial separation of Na_V_s concentrates *right-shift* in the proximal AIS, by concentrating Na_V_1.2 in that region (see Figure 1).

Our model sets the nominal *right-shift* of Na_V_1.2 at *V*_RS_ = 13.0mV for compatibility with [14]. We use the parameter Δ*V*_RS_ to alter, in the AIS only, the *right-shift* of specific Na_V_1.2 gating properties. (For additional details, see Section S1.6.2 and Section S1.6.3.) If a gating property (e.g. 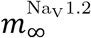) is listed in the legend of Figure 6, then its voltage-dependence is displaced by Δ*V*_RS_ (see, e.g., Equation 2). Likewise, if a gating property is *not* displayed in the legend, its voltage-dependence will *not* be modified during the sensitivity analysis.

The *activation* 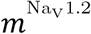 and *availability* 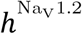 of real Na_V_1.2 channels in nature are *right-shift*ed by similar amounts relative to their Na_V_1.6 counterparts [20, 14, 12, 36]. (Although certain receptors can temporarily *right-shift activation* without shifting steady-state *availability* [37].) In our simulations, we can simultaneously decrease (or increase) the *right-shift* of 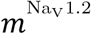 and 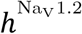 together, making the Na_V_1.2 channels more (or less) similar to the Na_V_1.6 channels in the AIS (respectively). This produces the ‘reference curves’ legended 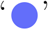 in Figure 6a and Figure 6b. In those curves, Δ*V*_RS_ shifts the voltage-dependence of every Na_V_1.2 gating property.

That is, as functions of membrane potential, the steady-states and time constants in the curves marked 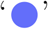 are 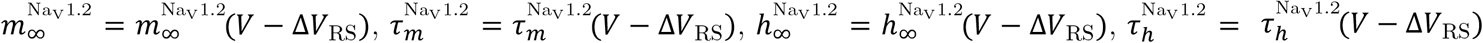 in the AIS.

Figure 6 connects local gating properties in the AIS, and their influence on the backpropagation threshold under somatic and axonal stimulation, to the effects of altering the Na_V_ distribution (seen above in Figure 4 and Figure 2, respectively). For example, making Δ*V*_RS_ negative will *left-shift* the voltage-gated sodium current in the proximal AIS, which is analogous to adding more proximal Na_V_1.6. However, this is merely an analogy: With *x* = 1 and *κ* ≳ 0.8, the proximal AIS has only Na_V_1.2 channels, and positive values of Δ*V*_RS_ will *right-shift* the sodium current in that area beyond what is attainable by changing the local mix of Na_V_ subtypes.

The reference curve (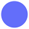) in Figure 6a shows that *right-shift*ing Na_V_s in the AIS —akin to replacing Na_V_1.6 channels with more Na_V_1.2— *increases I*_*BP*_ for somatic stimulation. And the reference curve in Figure 6b confirms that proximal *right-shift* from Na_V_1.2 *lowers I*_*BP*_ for axonal stimulation.

Since Δ*V*_RS_ only affects Na_V_1.2 channels *within the AIS* (example provided in Figure S16b), we can determine which *right-shift*ed gating properties drive the changes to *I*_*BP*_ that occur when the Na_V_ distribution is altered. To make said observation, in Figure 6 we “shift-clamp” selected gating properties: We ignore the experimental fact that Na_V_ *activation* and *availability* tend to *right-shift* in unison [20, 36, 38, 39], and that gating variables’ steady states and their voltage-sensitive time constants *right-shift* together as well [40].

Instead, we isolate the effects of individual gating properties by

1. applying Δ*V*_RS_ to 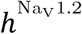 independently of 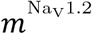 (curves legended 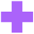 and 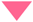 respectively),

2. applying Δ*V*_RS_ to 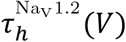 (*V*) independently of 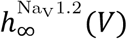 (*V*), and likewise for 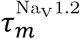,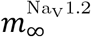; (curves legended 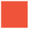,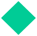 and 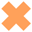, 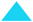 respectively),

And compute the new backpropagation threshold *I*_*BP*_(Δ*V*_RS_). (For mathematical details, see Section 4.2.)

An example of the latter is Equation 2 below, in which only 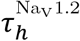 has its *right-shift* modified by Δ*V*_RS_ (see Figure S16b). The other three Na_V_1.2 variables have the nominal *right-shift* of 13mV. The *I*_*BP*_(Δ*V*_RS_) curves in Figure 6 that correspond to Equation 2 are legended 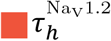:

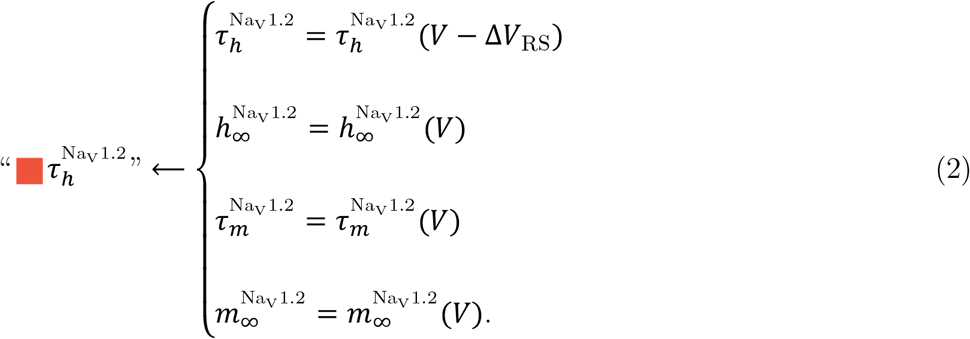

(Figure S16 visualizes the impact of Equation 2 on gating properties as a function of position at *V*_rest_.)

To unpack these additional curves, we begin at the coordinate we will call ‘the nominal point’ in each plot of Figure 6, which is the backpropagation threshold at Δ*V*_RS_ = 0, where all curves must intersect by definition. Starting at the nominal point, as one moves leftward along a given curve (Δ*V*_RS_ < 0), the gating properties indicated in the legend are *left-shift*ed (e.g. Equation 2), and the other gating properties are left alone. Likewise, travelling away from the nominal point to the right (Δ*V*_RS_ > 0) will *right-shift* the indicated properties, relative to their nominal kinetics (Figure S15).

Figure 6b reveals that, with axonal stimulation, the *right-shift*ed *availability* (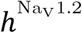) drives *I*_*BP*_. Specifically, the curves legended 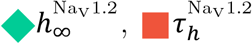 and 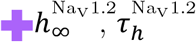 show how 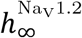 (the *availability* at steady-state, as a function of membrane potential) and 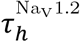 (the voltage-dependent time constant of *availability*) work together to promote backpropagation: They drive the reference curve (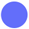) downward as Δ*V*_RS_ increases, in spite of the higher *activation* threshold. In other words: The threshold-lowering effects that result from *right-shift*ing the *availability* overpower the opposing influence of *right-shift*ed *activation* —on its own the latter would raise the threshold (see the curves: 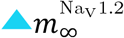 and 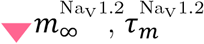 in Figure 6b).

Further, removing the *right-shift* from 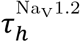 stops backpropagation: In the 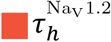 curve of Figure 6b, all gating properties other than 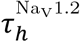, including 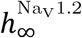, retain their nominal *right-shift*, yet backpropagation ceases (according to our strict BAP criterion, see Section S1.2) for axonal stimulation when Δ*V*_RS_ ≲ −2mV.

For somatic stimulation, the *right-shift*ed *activation* (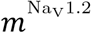) drives *I*_*BP*_. Travelling from right to left in Figure 6a, the most significant decrease in threshold occurs in the curves legended 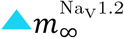 and 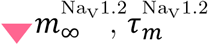 as the nominal *right-shift* is removed (Δ*V*_RS_ ⟶ −13.0mV ⟹ *V*_RS_ + Δ*V*_RS_ ⟶ 0, see Figure S15). The 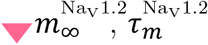curve differs negligibly from the 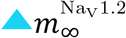 curve, showing that 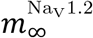 *right-shift* dominates in raising the threshold near the nominal point, and the *right-shift* of 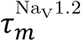 matters little.

Our Δ*V*_RS_ results from Figure 6 are summarized in Table 1.

**Table 1:** Summary of sensitivity analysis: Impact of *right-shift*ed Na_V_1.2 gating properties in the AIS on backpropagation threshold (see Figure 6).

| Type of Stimulation | reference curve | Increasing $\Delta V_{\text{RS}}$ of *availability* | | |
| --- | --- | --- | --- | --- |
| | | $\blacksquare \tau_h^{\text{Na}_V1.2}$ | $\blacklozenge h_{\infty}^{\text{Na}_V1.2}$ | $\blackplus h_{\infty}^{\text{Na}_V1.2}, \tau_h^{\text{Na}_V1.2}$ |
| Axonal | ↓ | ↓ | ↓ | ↓ |
| Somatic | ↑ | × | $\downarrow(\text{slight})$ | $\downarrow(\text{slight})$ |
| Increasing $\Delta V_{\text{RS}}$ of *activation* | | | | |
| | | $\blacktimes \tau_m^{\text{Na}_V1.2}$ | $\blacktriangle m_{\infty}^{\text{Na}_V1.2}$ | $\blacktriangledown m_{\infty}^{\text{Na}_V1.2}, \tau_m^{\text{Na}_V1.2}$ |
| Axonal | ↓ | $\uparrow(\text{slight})$ | ↑ | ↑ |
| Somatic | ↑ | × | ↑ | ↑ |

Arrows (↓,↑) indicate the sign of the **slopes** of backpropagation threshold curves in Figure 6a (somatic stimulation) and Figure 6b (axonal stimulation). A downward arrow (↓) indicates that the backpropagation threshold *I*_*BP*_ decreases with increasing *right-shift*, applied to the gating properties specified above it. Likewise, an upward arrow (↑) indicates that the threshold increases when the specified combination of Na_V_1.2 variables is *right-shift*ed. In cells marked “×” the effect of *right-shift* was negligible.

### 2.6 Generalization to other models

To demonstrate that our primary result —the separation of Na_V_1.2 and Na_V_1.6 into the proximal and distal AIS, respectively, promotes backpropagation with axonal stimulation but can *impede or promote* backpropagation with somatic stimulation— is not an artifact of our implementation of the model from Hu et al. (2009) [14] used thus far, we have inserted Na_V_ distributions analogous to Figure 1 into the model of Hay et al. (2011) [27] below. (Also, in Section S1.4, we include an alternate tuning of the Hu model, with a less excitable soma and robust backpropagation in the entire dendritic tree without attenuation. Despite its significantly higher dendritic excitability and qualitatively different backpropagation pattern, the results reported above are reproduced there as well.)

We replaced the single population of Na_V_s in the Hay model’s 60μm AIS with two Na_V_ subtypes, based on their original Na_V_ kinetics: One population *left-shift*ed by 6mV and the other *right-shift*ed by 6mV, relative to the original *V*_1/2_, to represent Na_V_1.6 and Na_V_1.2 respectively. Our manipulations of the Na_V_ channels’ distribution (varying *κ* and *x*) did not change the total Na_V_ density in the AIS, which was kept identical to their model (https://modeldb.science/139653). Further, we attached an additional 400μm-long section of passive cable to the end of the AIS, where their axon originally stopped, to allow the AP to exit the AIS orthodromically as well as antidromically, as is the case in real neurons, in order to make AP generation in the Hay model more realistic. This was necessary to recover our qualitative results.

Our intention was to modify the Hay model as little as was necessary, since its parameters are tailored to a specific neuron and morphology —they will not necessarily transfer well even between specimens of the same cell type (see Hay et al., 2011 [27]). Presumably, the tuning may also be sensitive to the excitability of newly attached compartments.

We note that Hay et al. optimized their models to fit experimentally observed somatic and dendritic spiking patterns, including BAC firing, but their focus was not on action potential initiation. The models that best fit their data had AP initiation in the soma rather than the AIS, but they provide an additional model where APs were constrained to initiate in a 60μm section named “axon”, which had been set aside due to excessive BAP attenuation (see [27]). Since we required a parameter tuning with AP initiation in the AIS, the latter model was the necessary choice, despite its unrealistically strong attenuation of the backpropagated AP.

In Figures 7 and 8 we register backpropagation in the Hay model [27] as either a spike in the soma, or a depolarization of several mV in the apical dendrites, following current injection. The threshold was set at 10mV in these figures for the somatic criterion; and at −70mV when measuring the depolarization near the bifurcation of the main apical dendrite, where *V*_rest_ ≅ −74.1mV. See Figure S5.

**Figure 7:**
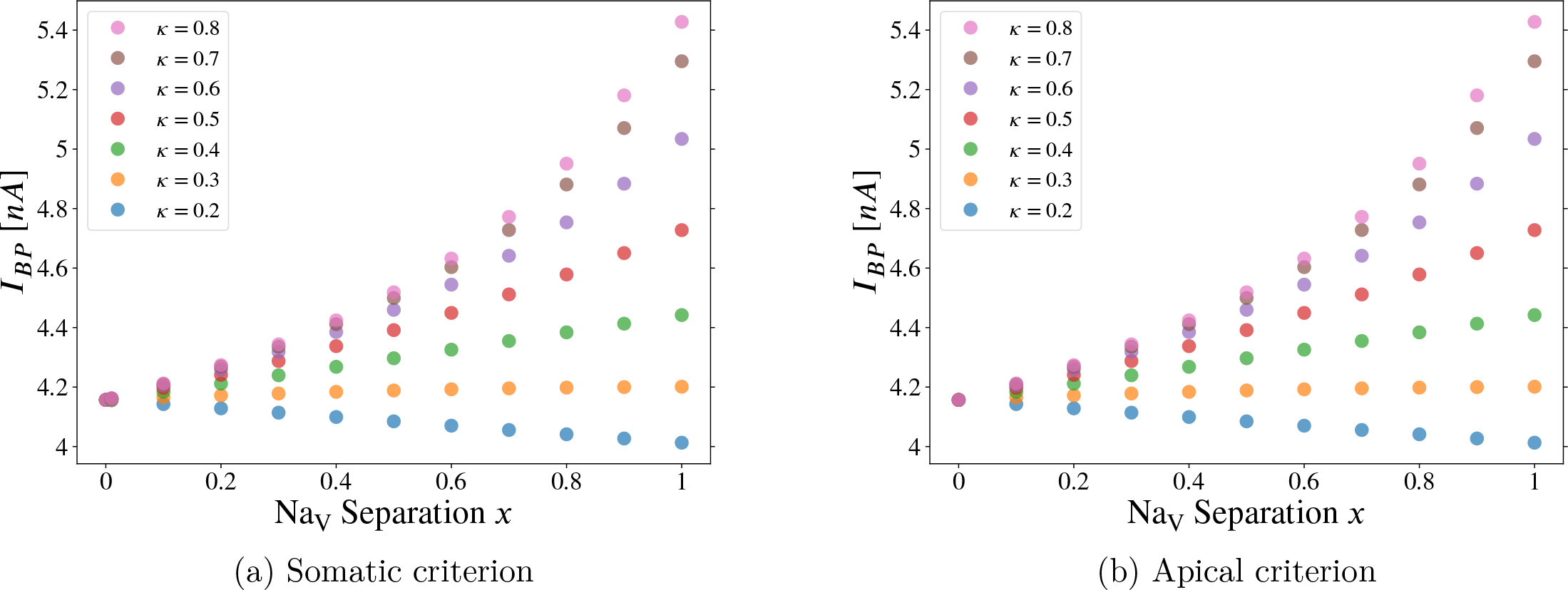
Somatic stimulation in the Hay model: Equivalency of apical vs. somatic back-propagation criteria. Combined effect on the backpropagation threshold (*I*_*BP*_, defined below) of varying crossover location (*κ*) and Na_V_ separation (*x*) in the axon initial segment. Note the qualitative and quantitative similarity to Figure 2. Varying the separation parameter “*x*” from *x* = 0 to *x* = 1, the distribution of Na_V_ channels goes from flat (homogeneous) to separated, the latter approximating the distribution observed in developing pyramidal neurons (see Figure 1a). Note that curves for all values of *κ* converge to a single point at *x* = 0 since *κ* can have no effect when the two Na_V_ subtypes are uniformly distributed along the AIS. Somatic backpropagation criterion = 10.0mV, apical backpropagation criterion = −70.0mV —see caption of Figure S5.

**Figure 8:**
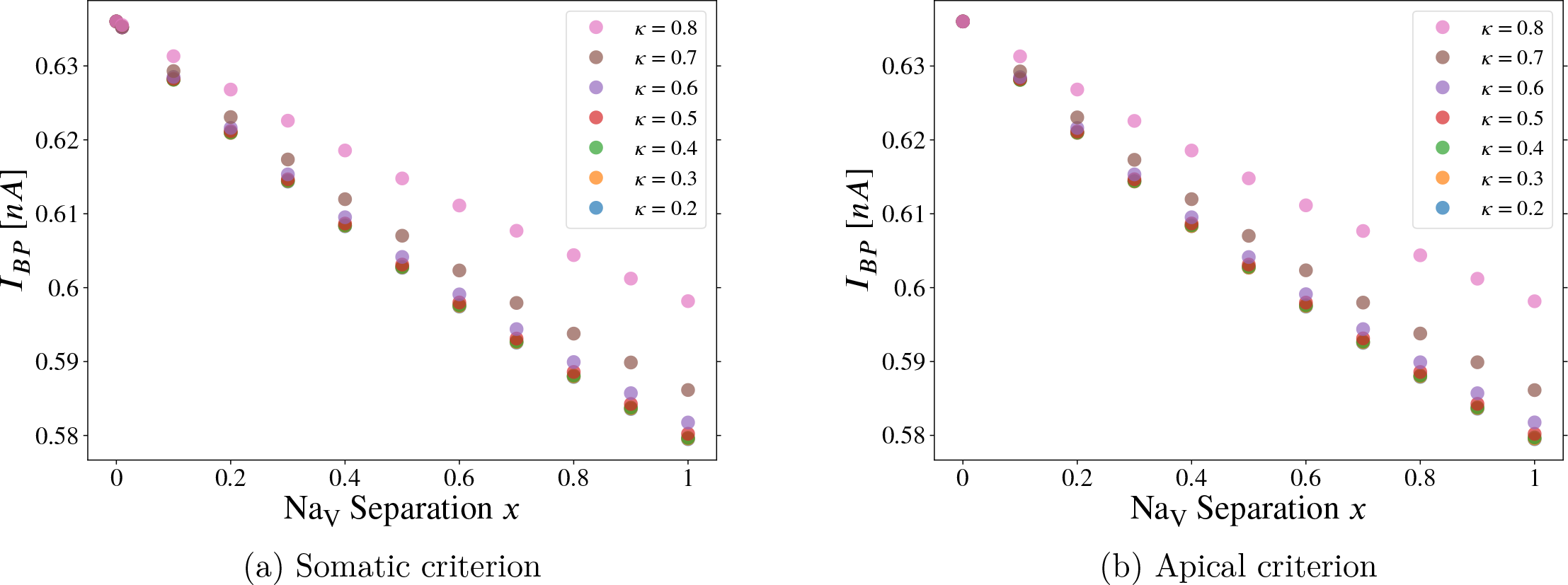
Axonal stimulation in the Hay model: Equivalency of apical vs. somatic backpropa-gation criteria. In the Hay model, the backpropagation threshold (defined above) is identical to the AP threshold. There is nowhere for saltatory conduction to occur, as there is no excitable axon beyond the AIS. Varying the separation parameter “*x*” from *x* = 0 to *x* = 1, the distribution of Na_V_ channels goes from flat (homogeneous) to separated, the latter approximating the distribution observed in developing pyramidal neurons (see Figure 1a). Note that curves for all values of *κ* converge to a single point at *x* = 0 since *κ* can have no effect when the two Na_V_ subtypes are uniformly distributed along the AIS. Somatic backpropagation criterion = 10.0mV, apical backpropagation criterion = −70.0mV —see caption of Figure S5.

Note the qualitative agreement between the Hay model implemented below and the Hu model above. In Figure 7 (below), we simulate backpropagation following somatic stimulation. As above in Figure 2, concentrating Na_V_1.2 in the proximal AIS tends to raise the backpropagation threshold, and increasing the proportion of total sodium conductance in the AIS allocated to Na_V_1.6 lowers *I*_*BP*_.

In Figure 8, we simulate backpropagation following axonal stimulation. As above in Figure 4 and Figure 5, the separated Na_V_ distribution (*x* ⟶ 1) lowers the backpropagation threshold in the Hay model. Quantitatively, Figure 8 is closer to Figure 5, suggesting that the concentration of low-threshold Na_V_1.6 at the distal AIS, rather than the concentration of Na_V_1.2 at the proximal AIS, promotes backpropagation. What is important to keep in mind is that, in both models, concentrating Na_V_1.2 in the proximal AIS only lowered *I*_*BP*_ in the unphysiological case of depolarizing axonal current injection.

In Figure 7a and Figure 8a, backpropagation is registered if the somatic membrane potential exceeds 10.0mV. The threshold need not be this high, but it does not affect the results since the somatic depolarization is large. In Figure 7b and Figure 8b, backpropagation is recorded in the main apical dendrite just distal to its main bifurcation (where *V*_rest_ ≅ −74.1mV) if the voltage exceeds −70mV. (See Figure S5a, Figure S5c.)

Since the Na_V_ distribution changes throughout development, a further investigation —beyond the scope of this paper, as we will explain— would be to understand how accompanying developmental changes in morphological complexity and voltage-gated channel density elsewhere in the neuron [41] interact with developmental plasticity in the AIS. This would require new parameter sets at each iteration of the morphological complexity. Since Hay et al. [27] had to fit each morphology’s parameter set to match firing patterns observed in real neurons, that procedure would need to be repeated. If sufficient experimental data are not available to perform the fitting at each iteration, new electrophysiological experiments would be necessary at the corresponding developmental stages. That endeavour is beyond the scope of the present study. Also, there is experimental evidence that AIS plasticity is not limited to development [6]. Our strategy was to restrict our investigation to the effects of varying the heterogeneous distribution of Na_V_ subtypes in the AIS on backpropagation threshold, with different modes of stimulation. Note, that the changing Na_V_ distributions we simulate are not strictly intended to replicate observed plasticity. Even if the Na_V_ distribution in the AIS of real neurons were static, modelling the hypothetical distributions would nonetheless assist in understanding its function; via the resulting changes to cellular excitability.

### 2.7 Rescaling the Na_V_1.2 density profile by a uniform factor in the AIS

In this section, we rescale the Na_V_1.2 density profile via the maximal conductance 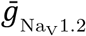. At each segment of the AIS, 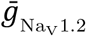 is multiplied by a positive number which we call the Na_V_1.2 scaling factor, denoted 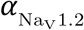. That is, at each point *s* in the AIS,

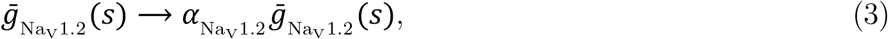

With 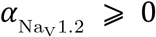. In Figure 9, we observe that reducing the density of Na_V_1.2, without adding any compensatory Na_V_1.6 density, increases the threshold as expected. Even when Na_V_1.2 is removed from the AIS and the proximal AIS contains no Na_V_ channels, backpropagation is possible with somatic stimulation.

**Figure 9:**
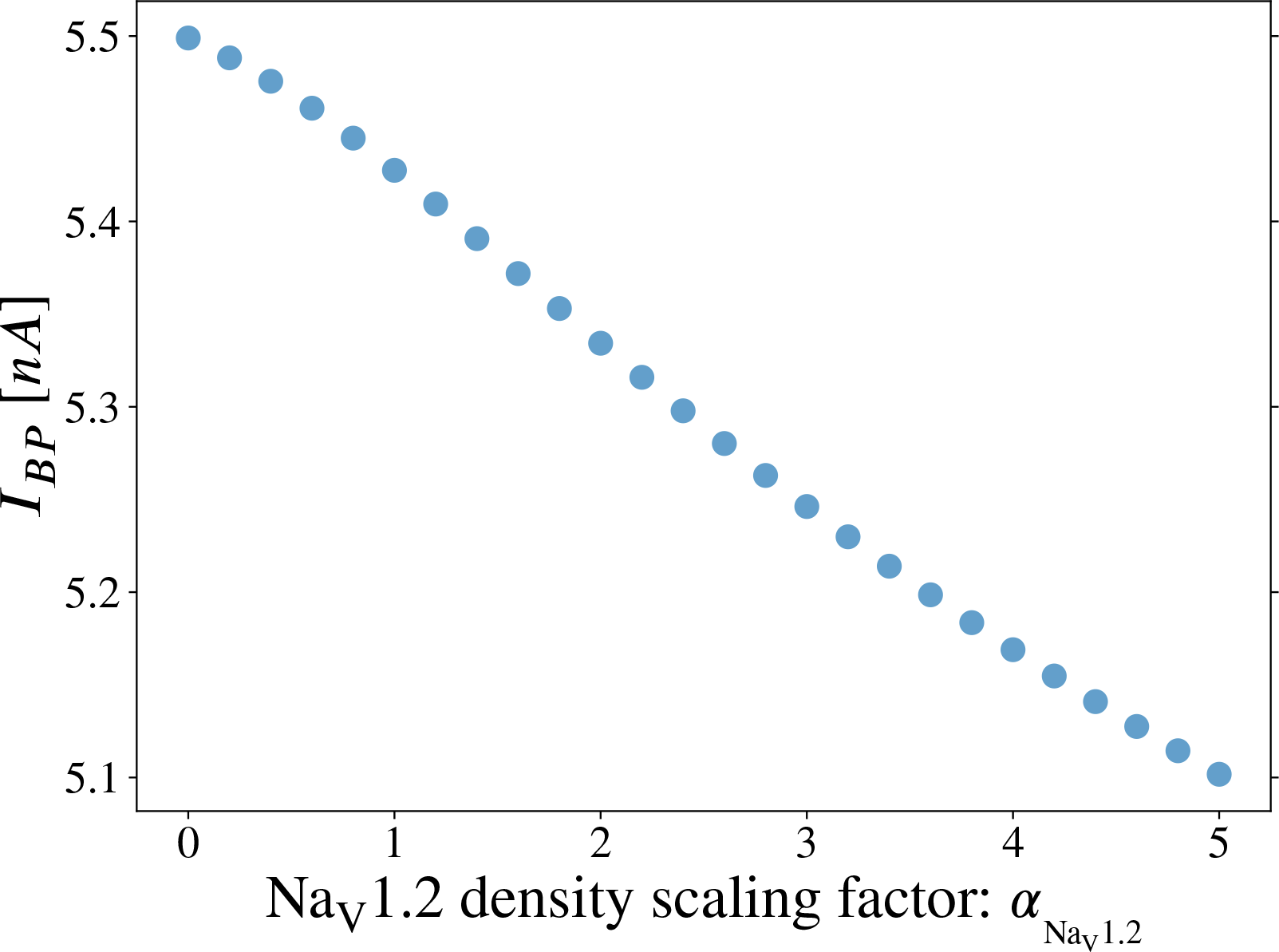
Scaling the Na_V_1.2 density profile in the AIS of the Hay model [27]. The backpropagation threshold is computed with somatic current injection (see Figure S5a, Figure S5b), while the Na_V_1.2 density profile is scaled globally by 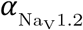. Here we have set *x* = 1 so that the Na_V_1.2 and Na_V_1.6 density profiles are separated (guaranteeing that the proximal AIS is exclusively populated with Na_V_1.2), and *κ* = 0.8 so that the majority of the AIS (except the most distal region, see Figure 1b) contains Na_V_1.2 and is therefore affected by 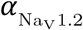.

In Figure 10 it is interesting to see, yet again, the sharp qualitative difference in the role of the Na_V_1.2 subtype with axonal versus somatic stimulation. Figure 10a uses somatic stimulation, and backpropagation remains possible when the Na_V_1.2 channels are disabled entirely (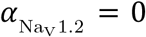). With axonal stimulation however (Figure 10b), the effect of 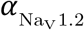 was abrupt and binary, akin to a Heaviside function. *I*_*BP*_ was nearly flat, except the Hay neuron refused to produce a BAP when 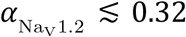 —some nonzero Na_V_ density was required in the proximal AIS for backpropagation.

**Figure 10:**
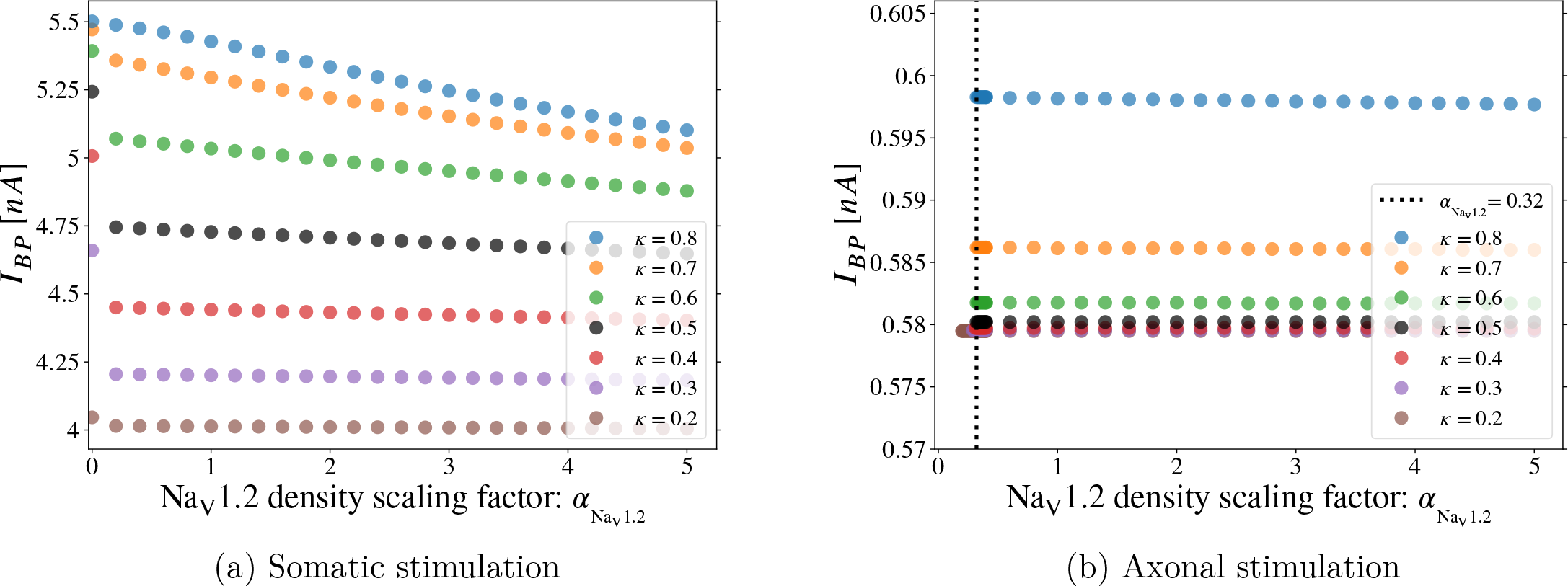
Scaling the Na_V_1.2 density profile in the AIS with somatic vs. axonal stimulation. The backpropagation threshold is computed while the AIS Na_V_1.2 distribution is scaled globally by 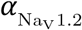. With somatic stimulation **(a)**, backpropagation persists even when Na_V_1.2 is removed (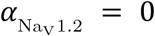). **However**, with axonal stimulation **(b)**, backpropagation ends abruptly near 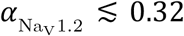. Yet again, the importance of the proximal Na_V_1.2 subtype and its qualitative effects on excitability depend heavily on the mode of stimulation. Density of data points increased in the vicinity of the vertical dashed line to detect backpropagation cutoff.

## 3 Discussion

In early development, pyramidal neurons concentrate Na_V_1.2 in the proximal AIS, and Na_V_1.6 in the distal AIS. As these cells mature, Na_V_1.6 invades the proximal AIS, and the two Na_V_ subtypes lose their separated distribution [19]. We have investigated the effects of Na_V_ separation in the axon initial segment on the initiation and backpropagation of action potentials in three different pyramidal neuron models. In spite of their different parameters, axonal and dendritic morphology, and biophysics, all three models (Section 2.2, 2.3, 2.5, S1.4) indicated that the effects of the separated Na_V_ distribution depend on whether stimulation is orthodromic (e.g. somatodendritic input) or antidromic (e.g. axonal stimulation).

With somatic stimulation, the greater the proportion of Na_V_1.2 in the AIS, relative to Na_V_1.6, the less excitable the cell becomes. This is contrary to past modelling which used axonal current injection [14], though it is consistent with the experimental results in [17] where somatic stimulation was used. The threshold-raising effect of proximal Na_V_1.2 is confirmed by repeating the simulations with a model cell in which the AIS has been flipped longitudinally (Figure 3), placing Na_V_1.6 proximally and Na_V_1.2 distally in the AIS.

Our results for axonal stimulation agree qualitatively with and expand upon past modelling efforts [14]. In all three models, excitability is greatest when Na_V_ subtypes are separated in the AIS (‘*x*-shaped distribution’). Further, in the Hu-based models (Figure 4, Figure S9), increasing the total proportion of Na_V_1.2 in the AIS —by moving the Na_V_ crossover *κ* distally— promotes backpropagation as well. In the Hay-based model, removing Na_V_s from the proximal AIS halted backpropagation. We also find that increased distal Na_V_1.6 concentration (which results from the separated distribution) lowers the AP threshold (Figure 5).

Testing both modes of stimulation can contribute to resolving inconsistencies between experiments such as [17] and [14], where stimulation was orthodromic in the former and antidromic in the latter. In [17], AP initiation was observed in pyramidal neurons which were engineered to be Na_V_1.6-deficient. In those neurons, the AIS was populated entirely with Na_V_1.2, however they still found that the AIS Na_V_ current was *left-shift*ed relative to the somatic current. From this and other observations, the authors in [17] suggest that the distribution of Na_V_ subtypes is not so important in shifting the local voltage-gated Na^+^ current.

We note that, compared to control neurons, the Na_V_1.6-deficient neurons’ AIS Na_V_ current was *right-shift*ed, and the orthodromic AP threshold (amplitude of a 2ms current pulse [17]) was nearly doubled. This is consistent with our results and the modelling assumption that *right-shift* is associated with Na_V_1.2 in the AIS —the model is agnostic about the molecular details. The decrease in excitability reported in [17] may have been even larger had they used more mature neurons. Their neurons were obtained from 4-5 week old mice, at which point the AIS will still be largely populated with Na_V_1.2, whereas in wild type mice Na_V_1.6 replaces much of the Na_V_1.2 by 90 days [19, 28]. Our results indicate that with axonal stimulation, Na_V_1.6-deficient cells may have a lower backpropagation threshold than the wild type.

The loss of the separated Na_V_ distribution in the AIS at later developmental stages, accompanied by the proximal localization of Na_V_1.6, may enhance excitability to healthy orthodromic stimulation while protecting against the backpropagation of ectopic activity from damaged axons into the soma and dendrites. Further, research into the genetic causes of autism spectrum disorder has revealed that Na_V_1.2 knockout can enhance pyramidal cells’ tendency to send action potentials and simultaneously reduce backpropagation (somatodendritic hypoexcitability) [18]. Whereas [18] reported an interplay between Na_V_1.2 and K_V_, in contrast, our results are explained by the spatial distribution of Na_V_ *right-shift* within the AIS (Table 1, Figure 6a). Indeed, the reduced excitability resulting from AIS Na_V_1.2 owes to the asymmetric impact of *availability* on backpropagation in axonal versus somatic stimulation (Figure 6).

Although *right-shift*ing Na_V_1.2 steady-state *availability* 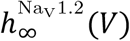 (*V*) in the AIS is necessary to promote backpropagation when stimulation is axonal, it is not sufficient on its own. Our modelling shows that the voltage-sensitive time constant 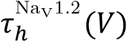 (*V*) must be *right-shift*ed as well (Figure 6b, 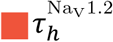 curve).

It is clear why increasing Δ*V*_RS_ lowers *I*_*BP*_ in the 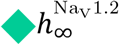 curves of Figure 6b and Figure 6a: *Right-shift*ing steady4-state Na_V_1.2 *availability* increases Na^+^ conductance at all voltages, because 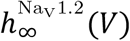 (*V*) is monotonically decreasing (Figure S15a). However, it was *not* obvious that removing the nominal *right-shift* from the voltage-sensitive time constant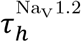—without modifying steady-state *activation* (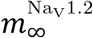) or *availability* (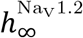)— should on its own be sufficient to eliminate the *I*_BP_ -lowering effects of Na_V_ 1.2 for axonal stimulation. But that is what our model has demonstrated in the curve legended 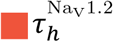 of Figure 6b. It follows that the *x*-distribution’s tendency to promote backpropagation is not merely a result of increased steady-state4*availability* of proximal AIS Na_V_s, but is a dynamic effect —dependent on the *right-shift* of 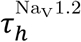 (*V*) as well.

There is an interplay between cable properties and the distribution of Na_V_s in determining the site of AP initiation [42]. Electrical isolation of the initiation site may amplify the effect of concentrating Na_V_1.6 in the distal AIS. Via fluorescence imaging of intracellular Na^+^ concentration following single action potentials, [33] located the greatest Na^+^ influx at the middle of the AIS, whereas the distal AIS (initiation site) had only 1/4 of this maximum. They inferred that the density of Na_V_ channels decreases toward the initiation site, and thus Na_V_ density does not determine the precise location where APs begin. Although [33] did not require Na_V_1.6 accumulation at the distal AIS to explain the distal location of the initiation site, the authors suggest that local Na_V_ density can have a large effect on neuronal excitability. Temperature may also play a role in local AIS Na^+^ influx measurements due to the spatial separation of Na_V_ subtypes. For high-speed spatiotemporal imaging of sodium influx, the pyramidal neurons in [33] and [17] were cooled to 21°. Na_V_1.2 and Na_V_1.6 differ in their responses to temperature changes [16]; thus, a deeper exploration of the effects of temperature on AP initiation is warranted. The temporal resolution of Na^+^ influx measurements continues to improve: [43] achieved a resolution of 0.1ms imaging pyramidal cells in mouse brain slices. Another order of magnitude improvement may be sufficient to discern the local contributions of Na_V_ subtypes to AP initiation. The qualitative dependence of the backpropagation threshold on the somatic-versus-axonal mode of stimulation is compatible with distal AP generation as found in [33, 14] and in our work, but does not seem to rely crucially on the precise determinants of AP onset position; it relies rather on the *activation* and *availability* properties, and the kinetics of the latter.

Our model neurons were kept identical in all results presented above; only the AIS was altered. Our results therefore can only be explained by the distribution of Na_V_ subtypes (or, the distribution of *right-shift*ed Na_V_ gating properties) within the AIS. Given the *τ*_*h*_-dependence of the antidromic backpropagation threshold in Section 2.5 and the differential temperature sensitivity of Na_V_1.2 versus Na_V_1.6 [16], there is good reason to expect that the effects of Na_V_ separation predicted here will be temperature-dependent.

In summary, we have simulated a range of hypothetical Na_V_ distributions in the axon initial segment of three 3D-reconstructed biophysical pyramidal cell models, including two distinct morphologies and three different parameter tunings. Our modelling shows that the spatial profile of Na_V_1.2 and Na_V_1.6 in the AIS and the kinetics of their *availability* and *activation* are important determinants of excitability and the back-propagation threshold. We predict that the separation of Na_V_ subtypes observed in early development has an asymmetrical effect on excitability which depends on whether the neuron is stimulated orthodromically or antidromically. With orthodromic stimulation, Na_V_ separation impedes backpropagation and reduces excitability unless the crossover is brought close to the soma. Backpropagation and excitability are both enhanced by Na_V_ separation when stimulation is antidromic. Maintaining a static Na_V_ distribution, we altered the *right-shift* of selected Na_V_1.2 gating properties. This revealed that steady-state *activation right-shift* controls the orthodromic backpropagation threshold, and dynamic *availability right-shift* is necessary to explain the antidromic threshold. Furthermore, given that learning is linked to backpropagation, the evolving separation of the Na_V_ subtypes may impact synaptic weight modification across developmental stages.

## 4 Methods

The pyramidal cell models were implemented in NEURON 8.0 [44] via Python. For cell geometry, local membrane properties, additional simulations, and a variety of calculations, clarifications, and definitions, see Supplementary Information (Section 8), which has its own table of contents.

Our implementation of the Hay model (https://modeldb.science/230326) is detailed in Section 2.6. Aside from modifying the *right-shift* in axonal Na_V_ channels to create Na_V_1.2 and Na_V_1.6 variants in the AIS, and attaching an additional passive section to the end of the axon, our Hay-based model is biophysically and morphologically identical to the original. Section 2.7 is an exception to this, since the total Na_V_ conductance in the AIS was modified via 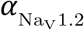.

Our Hu-based models [14] use the reconstructed morphology of a Layer 5 pyramidal neuron from cat visual cortex, modified from [24] (see SI, Section S1.1). Every compartment has explicit intracellular and extracellular concentrations of sodium, potassium, and chloride ions. The Nernst potentials 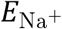, 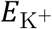, 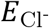 are calculated locally from each compartment’s specific ionic concentrations, which respond to transmembrane currents. Transmembrane concentration gradients of Na^+^ and K^+^ are maintained via active transport (Na^+^/K^+^-pump), and all ions are subject to longitudinal diffusion, both intra- and extracellular, implemented using NEURON’s RxD facility [45]. The cell maintains a resting potential *V*_rest_ ≅ −70mV at steady-state, and restores this state following stimulation. The biophysics that governs local ion concentrations (and Nernst potentials) in the Hu-based model is summarized in Section 4.3.

### 4.1 AIS - Na_V_ density profiles

In our model, the density profiles of Na_V_1.2 and Na_V_1.6 are left- and right-handed sigmoidal functions (respectively) of normalized length *s* along the AIS. The proximal end of the AIS is located at *s* = 0, and the distal end is located at *s* = 1. The channel densities are expressed as maximal conductances 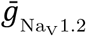 (s) and 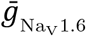 (s), where the total maximal Na_V_ conductance 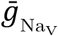 is constant along the AIS:

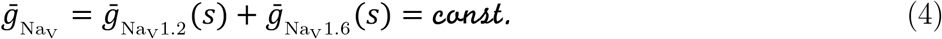

The density profiles are given by

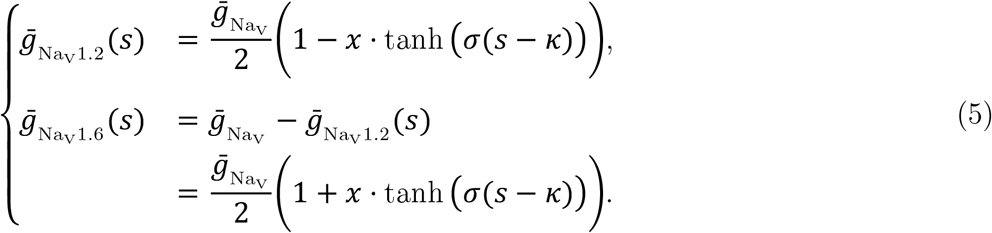

We chose the hyperbolic tangent function tanh(*s*), but other sigmoidal functions would do just as well. The parameter *x* controls the separation of the Na_V_ distribution, that is, how separated the two Na_V_ subtypes are. When *x* = 0, the distribution becomes flat —Na_V_1.2 and Na_V_1.6 are mixed uniformly along the AIS. When *x* = 1, the proximal end of the AIS contains only Na_V_1.2, and the distal end of the AIS contains only Na_V_1.6. The parameter *σ* is the reciprocal of the ‘transition width’ of the AIS Na_V_ distributions normalized by the AIS length. In all simulations shown here, *σ* = 10.0. Additional details are provided in Section S1.5.

### 4.2 Shift-Clamping and the Hodgkin-Huxley model

In the Hodgkin-Huxley model [40] a gating variable *u* evolves according to its voltage-dependent forward and backward transition rates *α*_*u*_(*V*) and *β*_*u*_(*V*) as

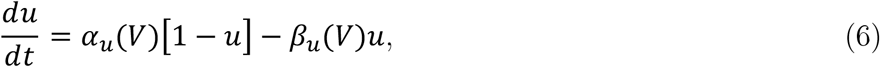

where *u* could be Na_V_ *activation m* or *availability h*, or K_V_ *activation n*, etc. This can be rewritten using the steady-state *u*_∞_(*V*) and voltage-dependent time constant *τ*_*u*_(*V*) of the gating variable

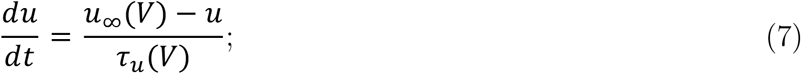

where *u*_∞_ and *τ*_*u*_ are computed from *α*_*u*_ and *β*_*u*_ via

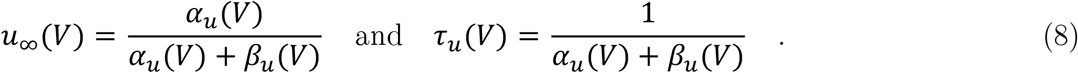

When shifting the voltage-dependence of *u*_∞_ by Δ*V*_RS_, it is natural to assume that one should apply the same shift to *τ*_*u*_ given Equation 8, since *u*_∞_ and *τ*_*u*_ are both functions of *α*_*u*_(*V*) and *β*_*u*_(*V*) in such models. However, our simulations can shift *u*_∞_(*V*) or *τ*_*u*_(*V*) independently of one another: 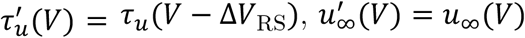. The forward and backward rates become

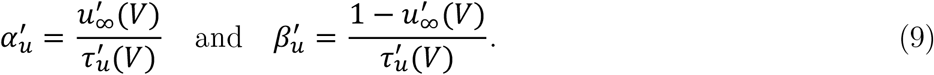

Putting this to use, one can modify the *right-shift* of combinations of

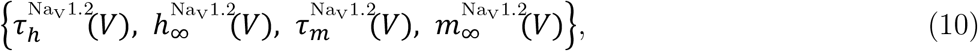

by adding “−Δ*V*_RS_” to the argument of the selected variables’ *u*_∞_(*V*)s or *τ*_*u*_(*V*)s.

### 4.3 Biophysics, Hu-based models

Action potentials propagate via the cable equation

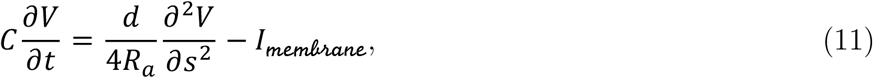

where *V* is the membrane potential, *C* is the specific membrane capacitance, *d* is the neurite diameter, *R*_*a*_ is the axial resistance, *s* is the position along the axis of the cable, and *I*_*membrane*_ is the total transmembrane current density of all ion species in the model.

In the Hu-based model, each compartment has explicit intracellular and extracellular concentrations of sodium, potassium, and chloride ions. We denote the intracellular/extracellular concentration of a given ionic species “Z” as [Z]_in_, [Z]_out_ respectively. These concentrations depend on the spatial coordinate —i.e. [Z]_in_ = [Z]_in_(*s*)— but that is not written explicitly, to simplify the notation. The Nernst potentials (reversal potentials) 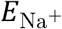, 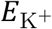, 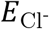 of Na^+^, K^+^, and Cl^-^ are not fixed parameters but are instead determined by the intracellular and extracellular concentrations of those ions:

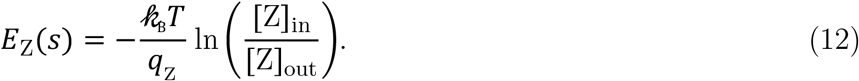

Transmembrane concentration gradients of Na^+^ and K^+^ are governed by active transport (Na^+^/K^+^-pump) and longitudinal diffusion. At each time step, ionic concentrations all over the cell are updated using transmembrane currents (Equation 15) and Fick’s law. At the *j*^*th*^ compartment this gives:

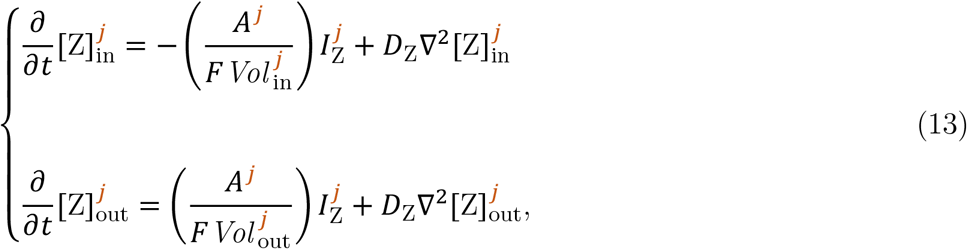

where 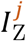 is the transmembrane current density of ion species Z at compartment *j*, with Z = Cl^-^, K^+^, Na^+^. *D*_Z_ denotes the diffusion coefficient of ion Z. *A*^*j*^ and 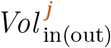 are (respectively) the membrane area and intracellular/extracellular volume at the *j*^*th*^ compartment. *F* is the Faraday constant. The total transmembrane current density at the *j*^*th*^ compartment is

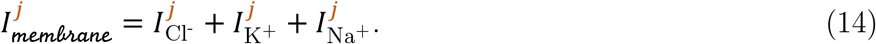

Omitting the compartment index *j*, the specific transmembrane currents are

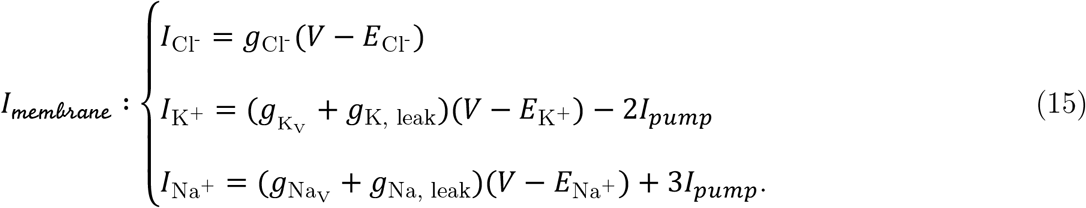

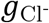, *g*_K, leak_, and *g*_Na, leak_ are passive leak conductances whereas *g*_K_ and 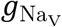 have voltage-gated Hodgkin-Huxley (HH)-style kinetics (Equation S6). Since channels are nonuniformly distributed along the cell membrane, conductances vary with location. *I*_*pump*_ is the net current produced by the Na^+^/K^+^-pump as a function of [K^+^]_out_ and [Na^+^]_in_,

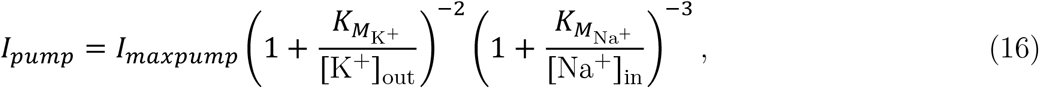

where *I*_*maxpump*_ controls the maximal pump current, 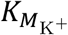 and 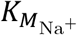 are Michaelis-Menten kinetic constants, and the Na^+^ and K^+^ currents flowing through the pump are 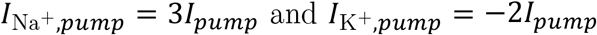. (Calcium dynamics are omitted in this section since Hu et al. [14] did not include the dendritic calcium spike initiation zone —see [35]. In Section 2.6 we include the Hay model, which features state-of-the-art calcium dynamics.)

### 4.4 Data availability

The simulation software that produces the results in this paper and the plotted data will be made available on ModelDB at https://modeldb.science/267088. Also, the Supplementary Information includes tables of parameters that detail the model setup.

### 4.5 Code availability

Model code will be posted on ModelDB upon publication at https://modeldb.science/267088.

## 5 Acknowledgments

We acknowledge the support of the Natural Sciences and Engineering Research Council of Canada (NSERC), to BJ and AL. We wish to thank: Catherine E. Morris, for directing our attention to the AIS. Nicholas T. Carnevale and Robert A. McDougal, for generously helping us to solve a number of model implementation challenges using NEURON. We also thank Louis Jacques for helpful discussion.

## 6 Author contributions

B. Barlow: conceptualization, model development, implementing the models and curating the outputs, coding, optimizing, model validation, figure preparation, and manuscript writing and revising; A. Longtin: conceptualization, model development, model validation, manuscript editing; and B. Joós: conceptualization, model development, model validation, manuscript editing. B. Joós and A. Longtin jointly supervised this work.

## 8 Supplementary information

### S1.1 Model: pyramidal cell geometry

Figure S1 displays the first of the two neurons used in this paper. The dendritic morphology is a digital reconstruction of a Layer 5 pyramidal neuron from cat visual cortex, modified from [24]. These neurons have dendrites roughly as tall as the thickness of the cortex (several mm) and axon initial segment length similar to the width of a human hair (tens of μm [9]).

**Figure S1:**
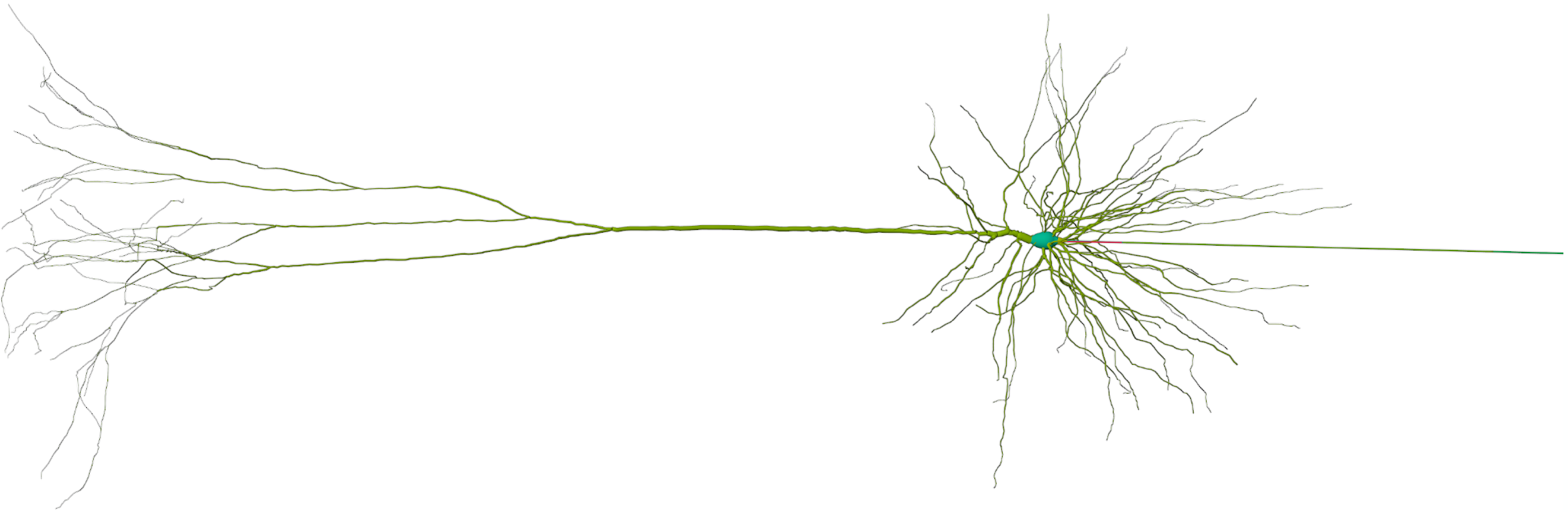
Geometry from the Hu model [14]: Layer 5 pyramidal neuron from cat visual cortex, modified from [24]. Somatodendritic morphology is digitally reconstructed from a real cell; hillock, AIS, and axon are added. Image created using blenderNEURON: blenderNEURON.org, blender.org.

The same reconstructed cell geometry was used in [14]. A cylindrical axon similar to that used in [14] was attached to the reconstructed soma by a 10μm long tapered hillock. The axon proper includes fifteen nodes of Ranvier (gray dots on the axon in Figure S3) separated by myelinated 100μm internodes. The axon initial segment length was set to *ℓ* = 25.0μm, consistent with [10]. Voltage-gated channels are present all over the cell. Channel densities and passive leaks are given as conductances. The local conductances vary with position —representative values have been tabulated in Section S1.8.

In past modelling, there is sometimes a long section of bare axon following the AIS, or myelination may begin immediately at the end of the AIS. To ensure our results did not depend on this morphological feature, we ran simulations (axonal and somatic stimulation) with and without a 400μm section of bare axon separating the AIS from the first internode. The effects of Na_V_ distribution, summarized in Figures S7 and S9, were qualitatively similar and similar in magnitude with and without the bare axon.

**Figure S2:**
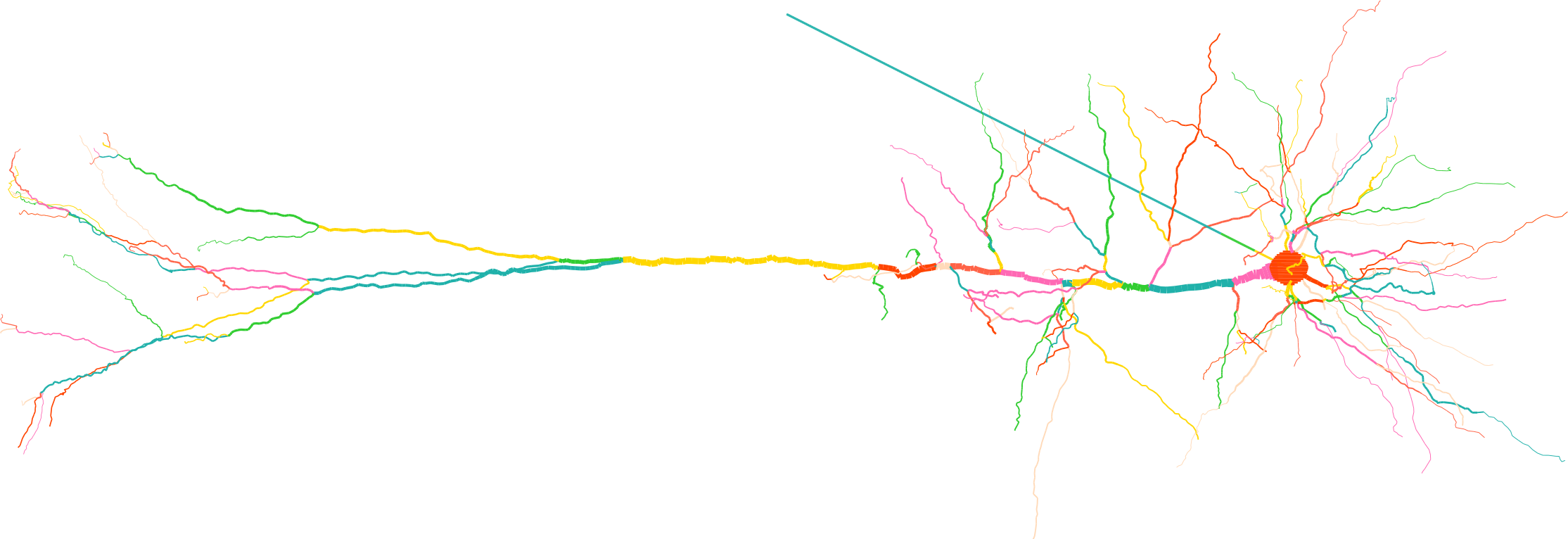
Morphology from Hay et al. [27]. In the Hay model (https://modeldb.science/230326), the AIS consists of two straight NEURON Sections called “axon[0]” and “axon[1]”, each 30μm in length, making the AIS 60μm long. The diagonal segment leading directly to the soma on the right is a 400μm long passive cable attached to the end of the AIS.

### S1.2 Spikes and backpropagation in the Hu-based model

When pyramidal cells spike, the action potential can travel backward into the dendrites. This phenomenon is called backpropagation. For example, Figure S3 shows the response of the neuron following a somatic current pulse just above (left) and just below (right) the backpropagation threshold *I*_*BP*_. Peak voltage is recorded across the entire cell model. The peak voltage at the tips of the dendrites is indicated in orange stars 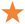 and has a higher amplitude, consistent with the sealed-end effect. Backpropagation was deemed to have occurred if all apical dendritic tips exceeded −63.0mV (i.e. a depolarization of 7.0mV above *V*_rest_) following stimulation.

**Figure S3:**
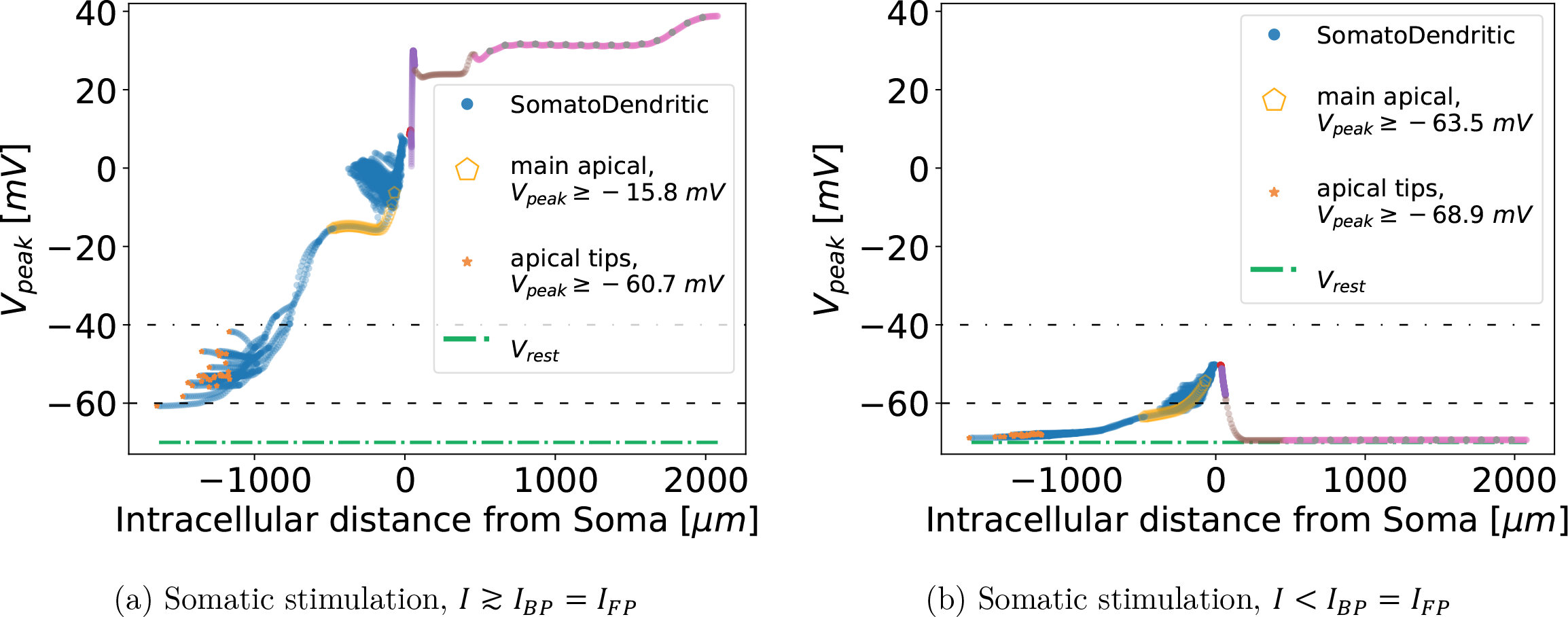
Backpropagation with somatic stimulation. Each data point maps to a location in the reconstructed pyramidal neuron. On the abscissa, negative values indicate that a datapoint is located in the soma or dendrites, and positive values correspond to the hillock, AIS, and axon. (The correspondence between the abscissa and cell morphology is illustrated in Figure S6.) (*x* = 1, *κ* = 0.7) This and similar figures are inspired by “Figure 4” of [11].

Note the qualitative change that occurs in the peak dendritic voltages of Figure S3a and Figure S4a, when the cell is above threshold. A minuscule increase in the injected current amplitude has caused the entire main apical dendrite to depolarize well above −40mV, despite being mostly below −60mV when the current was slightly below this threshold. Note the attenuation in the peak voltage when comparing the basal dendrites to the apical tips. Also note the variety of different peak voltages in the dendrites, compared to the subthreshold condition. Our backpropagation criteria for the Hu models required this qualitative change since it was a robust feature of those models. Recall also that the purpose of the dendrites in our model is to define a backpropagation threshold, not necessarily to mimic the detailed features of backpropagation. For our purposes, it is enough that the dendrites exhibit a threshold, and that threshold changes as a result of altering the distributions of Na_V_s in the AIS.

**Figure S4:**
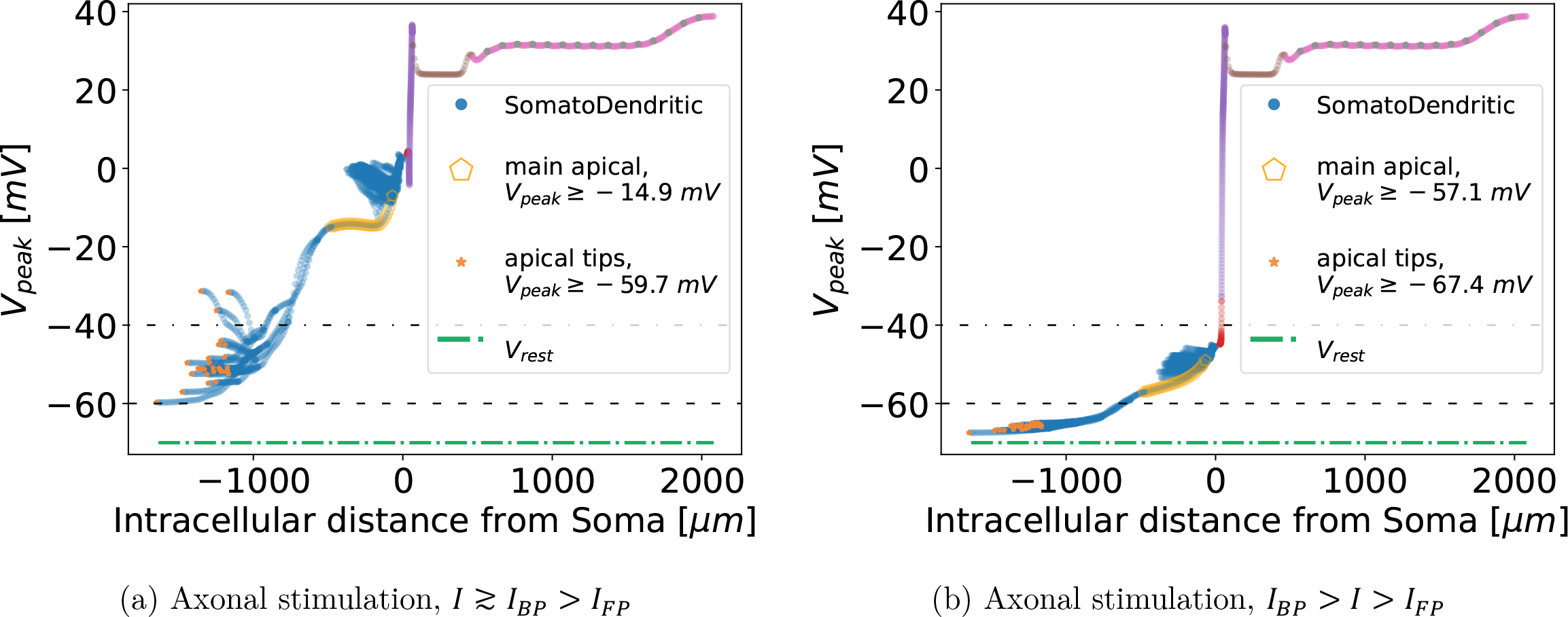
Axonal stimulation in the Hu-based model: spikes with and without backpropagation —by our stringent criterion. Current is injected just distal to the AIS. Each data point maps to a location in the reconstructed pyramidal neuron. (The correspondence between the abscissa and cell morphology is illustrated in Figure S6.) In **(a)**, an action potential (AP) has backpropagated. Note the variability of *V*_peak_ in the somatodendritic region of the neuron, and the significant attenuation as the wave travels into the distal dendrites. In **(b)**, an AP has occurred but the pattern of depolarization in the dendrites does not satisfy the backpropagation criterion we have used for this model. Note that the peak voltage in the soma and dendrites remains nearer to the resting potential ≈ −70mV, never exceeding −40mV. Also note the lack of variability in the somatodendritic *V*_peak_. The qualitative change in these two plots occurs sharply, just above *I*_*BP*_ ≈ 2.7nA (*x* = 1, *κ* = 0.4). By some definitions, both of these scenarios would be considered backpropagation, since there is always a few mV of depolarization in the most distal dendrites. As this qualitative change is a robust feature of the Hu-based model, we have defined backpropagation in those simulations to require it. In defining a forward propagation threshold (*I*_*FP*_), we call the scenario on the right, where an AP has been sent down the axon orthodromically from the stimulation site (i.e. to the right in this panel) “forward propagation” or “forwardprop” regardless of the amplitude of the somatodendritic depolarization. This and similar figures are inspired by “Figure 4” of [11].

### S1.3 Spikes and backpropagation in the Hay-based model

The backpropagation criteria we used with the Hay model[27] are given in the caption of Figure S5.

**Figure S5:**
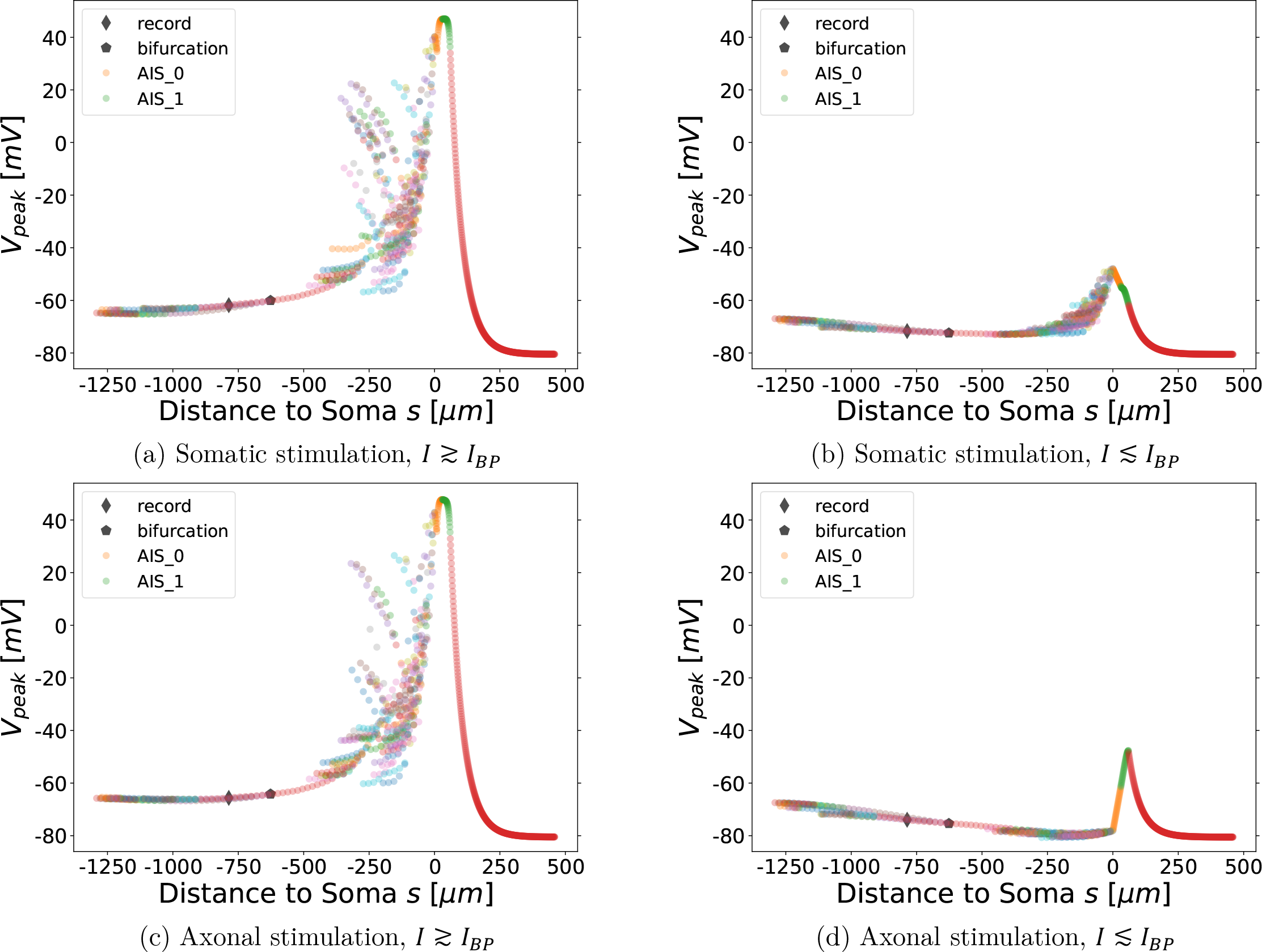
Backpropagation criteria in the Hay-based model. Each data point maps to a location in the reconstructed pyramidal neuron. On the abscissa, negative values indicate that a data point is located in the soma or dendrites, and positive values correspond to the AIS and passive cable. (The correspondence between the abscissa and cell morphology is illustrated in Figure S6.) In **(a)** and **(c)** backpropagation has occurred, whereas in **(b)** and **(d)**, the stimulation is (just) below the BAP threshold. In the model, backpropagation was recorded either at the soma (neighbouring the left end of the section labelled “AIS_0”, membrane potential exceeding 10mV), or in the apical dendrites (just distal to the main bifurcation of the main apical dendrite, membrane potential exceeding −70mV). The resting potential was ≅ −80.5mV at the soma, and ≅ −74.1mV at the apical recording site (see legend).

In Figure S5, compare the depolarization of the distal AIS (‘AIS_1’ in the legend) in the Hay model when current is injected somatically versus axonally, with the neuron just below *I*_*BP*_: In the somatic case (Figure S5b) the distal AIS never reaches −60mV, while in the axonal case (Figure S5d) it exceeds −50mV. Hay et al. did not include an axon[27], and here the AIS is followed by a passive cable, rather than an excitable axon composed of myelinated internodes segmented by nodes of Ranvier. Hence a forward-propagation threshold *I*_*FP*_ is ill-defined in this model. Were the Hay model to include such an axon, the increased depolarization of the distal AIS in (Figure S5d) compared to (Figure S5b) would be sufficient to cause an axonal AP in the former without doing so in the latter. Thus we argue, that the ability of the Hu-based model —with its excitable axon— to generate APs without meeting our criterion for backpropagation (Figure S4b), is an artifact of axonal stimulation.

### S1.4 Alternate tuning of Hu-based model

In this tuning, backpropagation is robust. A high amplitude, regenerative BAP infiltrates the entire dendritic tree. We redefined the threshold criterion such that backpropagation was deemed to have occurred if all apical dendritic tips exceeded −10.0mV following stimulation. The higher threshold value was appropriate to record backpropagation due to the robust BAP, which did not show attenuation (see Figure S6). Although this tuning diverges from the qualitative features of BAPs in real pyramidal cells, at least for single action potentials [26, 27], the relationship between the backpropagation threshold and the Na_V_ distribution presented in Section 2 was preserved again (see the figures below).

That is, despite the significant qualitative differences between the tuning of the Hu et al. [14] based model below (Figure S6) which lacks BAP attenuation, the other Hu et al. based model above in the main text (Section 2.2, Section 2.3) which we modified for BAP attenuation, and the Hay et al. [27] based model above in Section 2.6, qualitative effects of modifying the Na_V_ distribution in the AIS are identical, and for somatic stimulation they are quantitatively similar as well. This can be clearly concluded by comparing Figure S7, Figure 2, and Figure 7.

**Figure S6:**
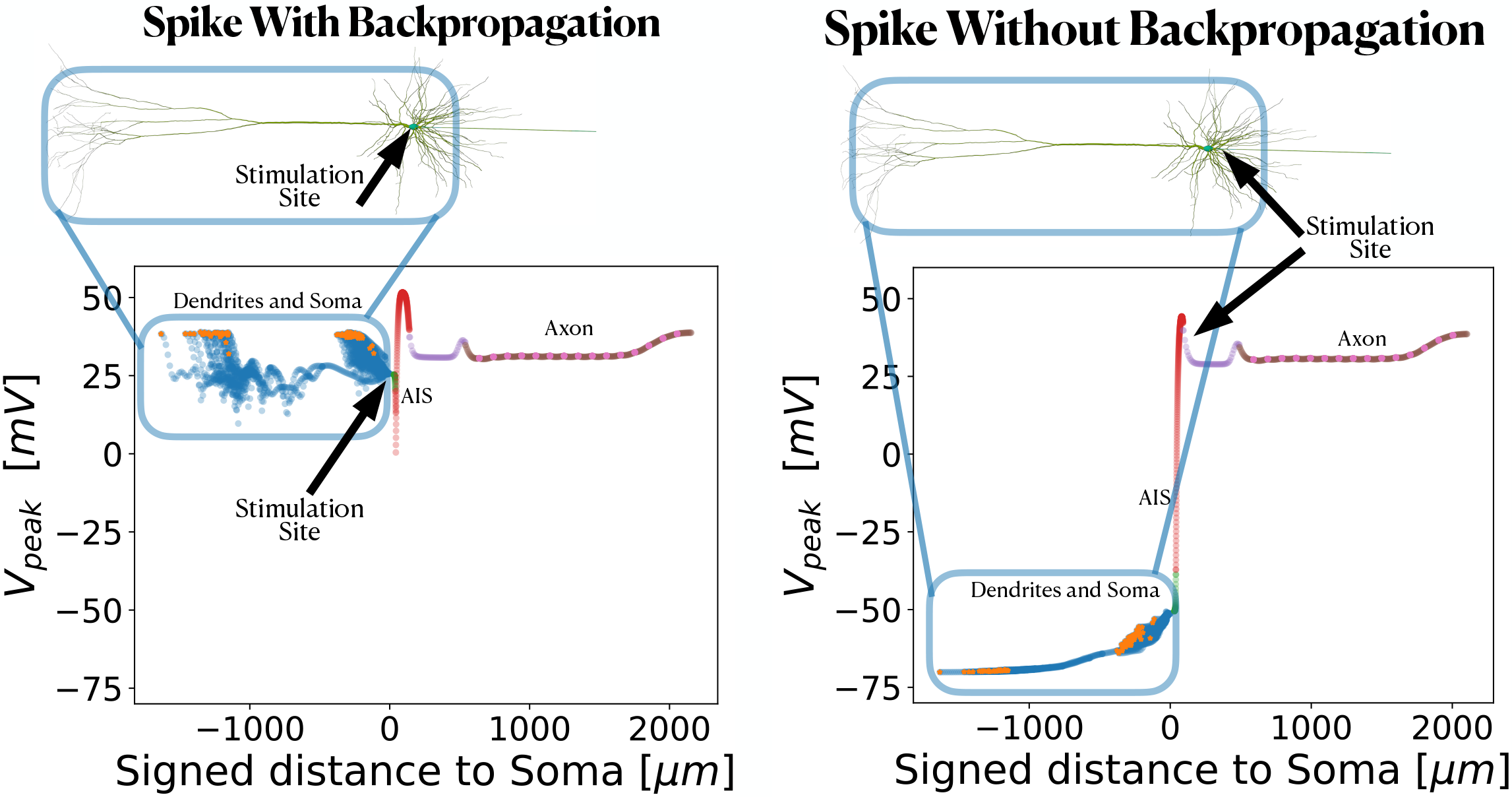
Spikes with and without backpropagation: Each data point maps to a location in the reconstructed pyramidal neuron (compare with the cell morphology above the plot). On the abscissa, negative values indicate that a datapoint is located in the soma or dendrites, and positive values correspond to the hillock, AIS, and axon. Beginning on the right-hand side: an action potential (AP) has occurred following axonal stimulation (current injected just distal to the AIS). Note that the peak voltage in the soma and dendrites remains near the resting potential ≈ −70mV, indicating that backpropagation did not occur. We call this scenario where an AP has been sent down the axon orthodromically from the stimulation site (i.e. to the right in this panel) “forward propagation” or “forwardprop” regardless of the amplitude of the somatodendritic depolarization. To the left is an AP that backpropagated: the entire cell spiked, in this case following somatic stimulation; backpropagation can also occur following axonal stimulation. The somatodendritic peak voltages are indicated by a blue box in each case. This and similar figures are inspired by “Figure 4” of [11].

**Figure S7:**
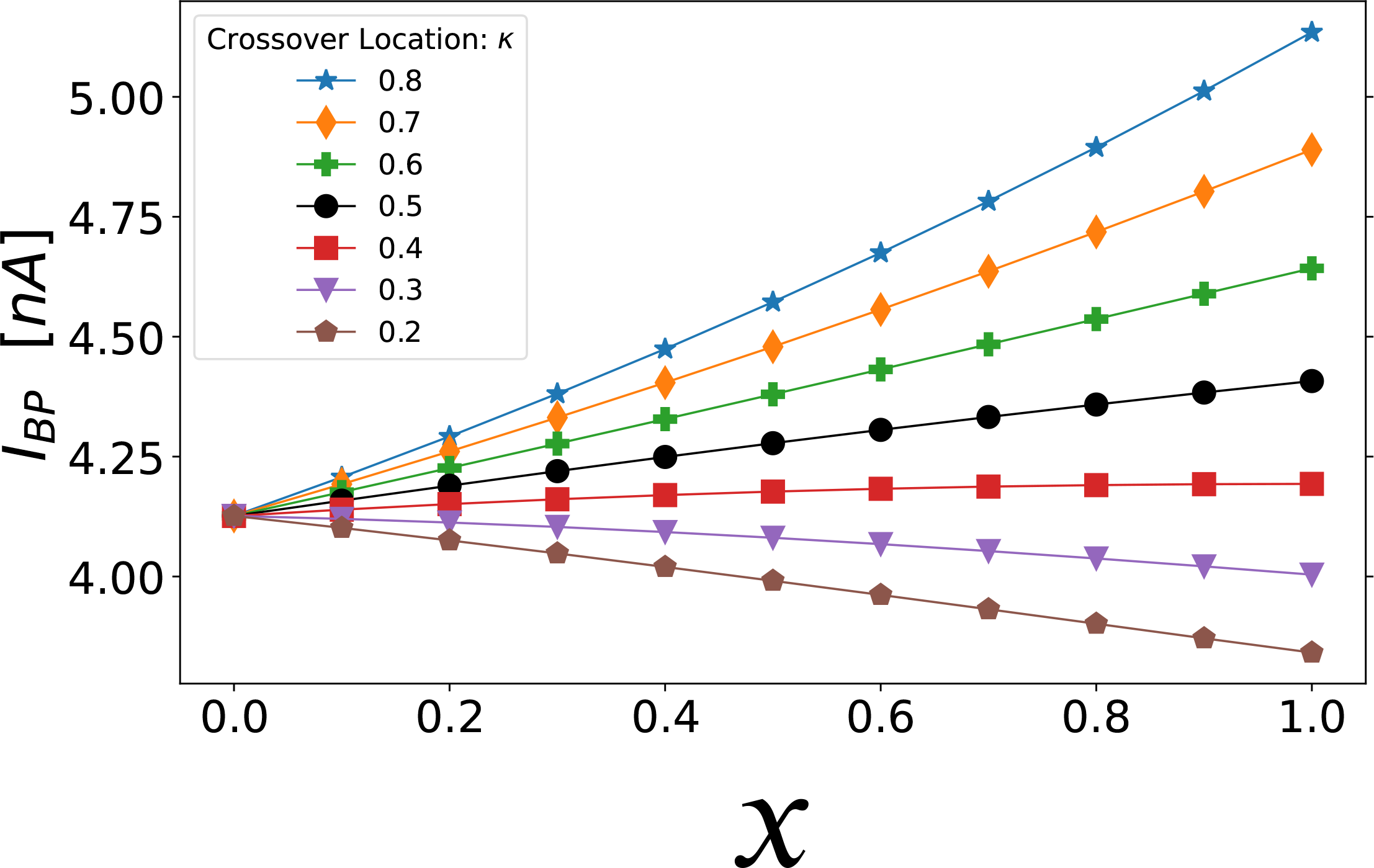
Somatic Stimulation: Combined effect of varying crossover location (*κ*) and Na_V_ separation (*x*) in the axon initial segment. The threshold for forward AP propagation is the same as for backpropagation. Varying the separation parameter “*x*” from *x* = 0 to *x* = 1, the distribution of Na_V_ channels goes from flat (homogeneous) to separated, the latter approximating the distribution observed in developing pyramidal neurons (see Figure 1a). Note that curves for all values of *κ* converge to a single point at *x* = 0 since *κ* can have no effect when the two Na_V_ subtypes are uniformly distributed along the AIS. The lines have been drawn to guide the eye.

**Figure S8:**
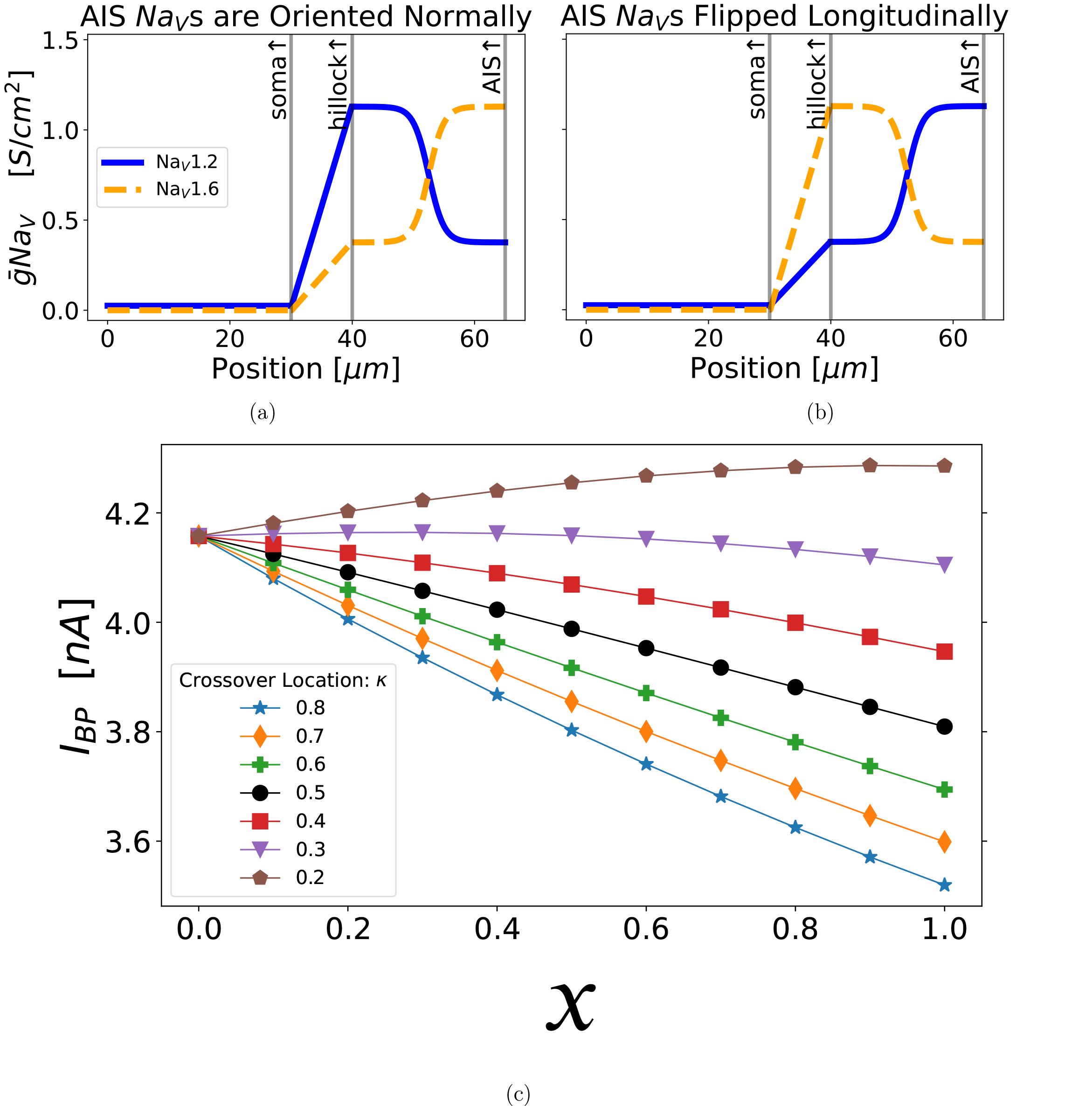
When the AIS Na_V_ distribution is flipped, setting *x* = 1 concentrates Na_V_1.6 at the proximal AIS and Na_V_1.2 at the distal AIS —the opposite of what is observed in many pyramidal cells [14, 15, 16]. (a) AIS with proper longitudinal placement of Na_V_s. (b) AIS with a longitudinally flipped Na_V_ distribution. In both plots, *x* = 0.5 and *κ* = 0.5. (c) Somatic stimulation with AIS Na_V_s flipped as in (b): This result is close to a mirror image of Figure S7. The lines have been drawn to guide the eye.

**Figure S9:**
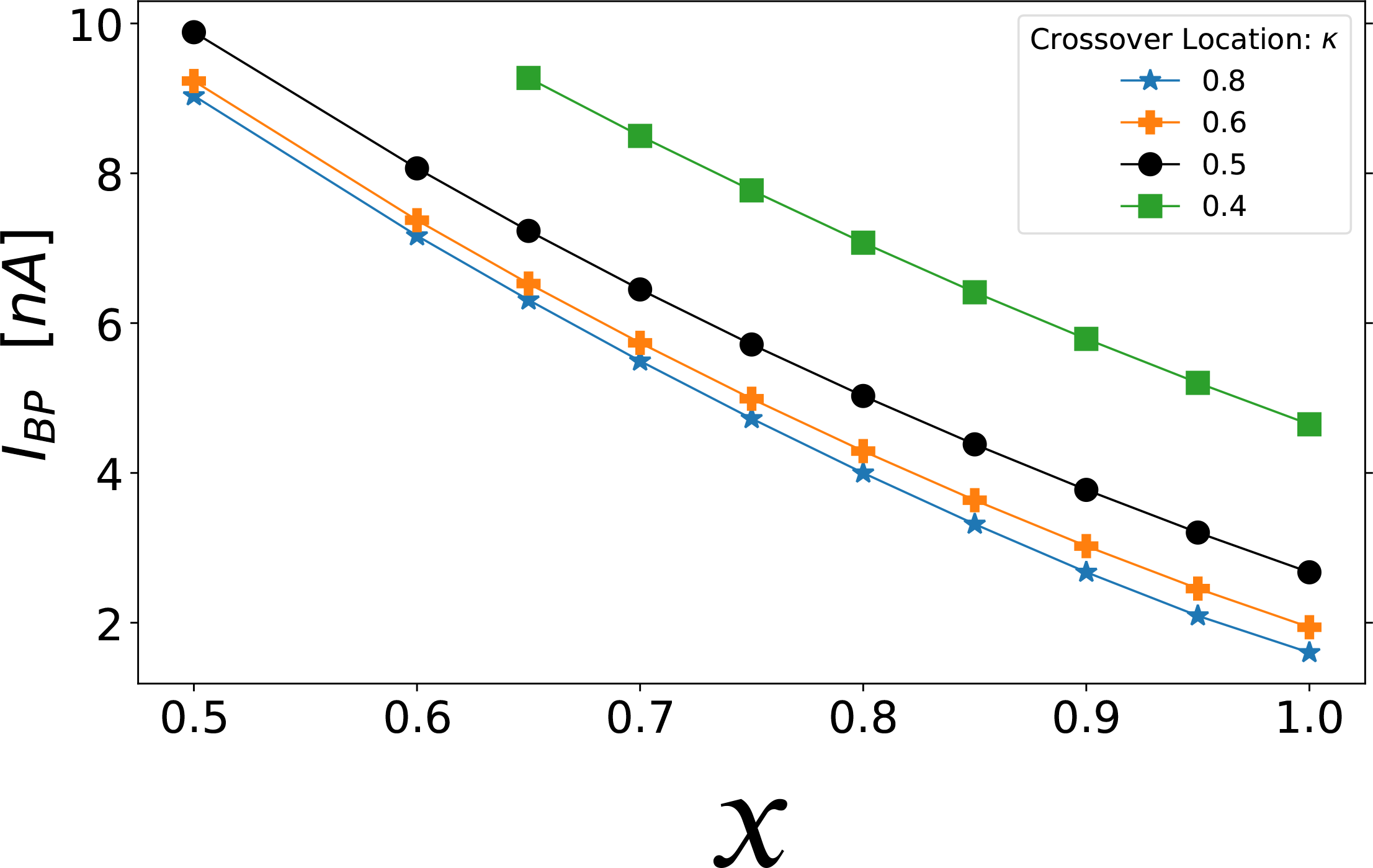
Axonal Stimulation: Effect of varying crossover location (*κ*) and Na_V_ separation (*x*) in the AIS on the backpropagation threshold (see Figure 1). When computing the threshold, the stimulating current was limited to a maximum of 10nA, to prevent unphysiological local depolarization at the stimulation site. Due to the smaller diameter of the axon (relative to the soma), 10nA is sufficient to depolarize the membrane potential to ≈ +80mV at the stimulation site, whereas the resting potential is *V*_rest_ = −70mV. To achieve backpropagation within that constraint (following axonal stimulation), our model required some amount of proximal Na_V_1.2, delivered through the combined effects of Na_V_ separation (*x* ≳ 0.5) and a sufficiently distal crossover position *κ* ≳ 0.4. Separating the two Na_V_ subtypes (*x* ⟶ 1) lowers the threshold, in agreement with the finding in [14] that proximal accumulation of Na_V_1.2 promotes backpropagation, albeit due to different gating properties (Figure 6b). Increasing *κ* raises the proportion of Na_V_1.2 (relative to Na_V_1.6) in the AIS and lowers the backpropagation threshold as well. Threshold changes here are larger than for somatic stimulation (Figure S7). The lines have been drawn to guide the eye.

**Figure S10:**
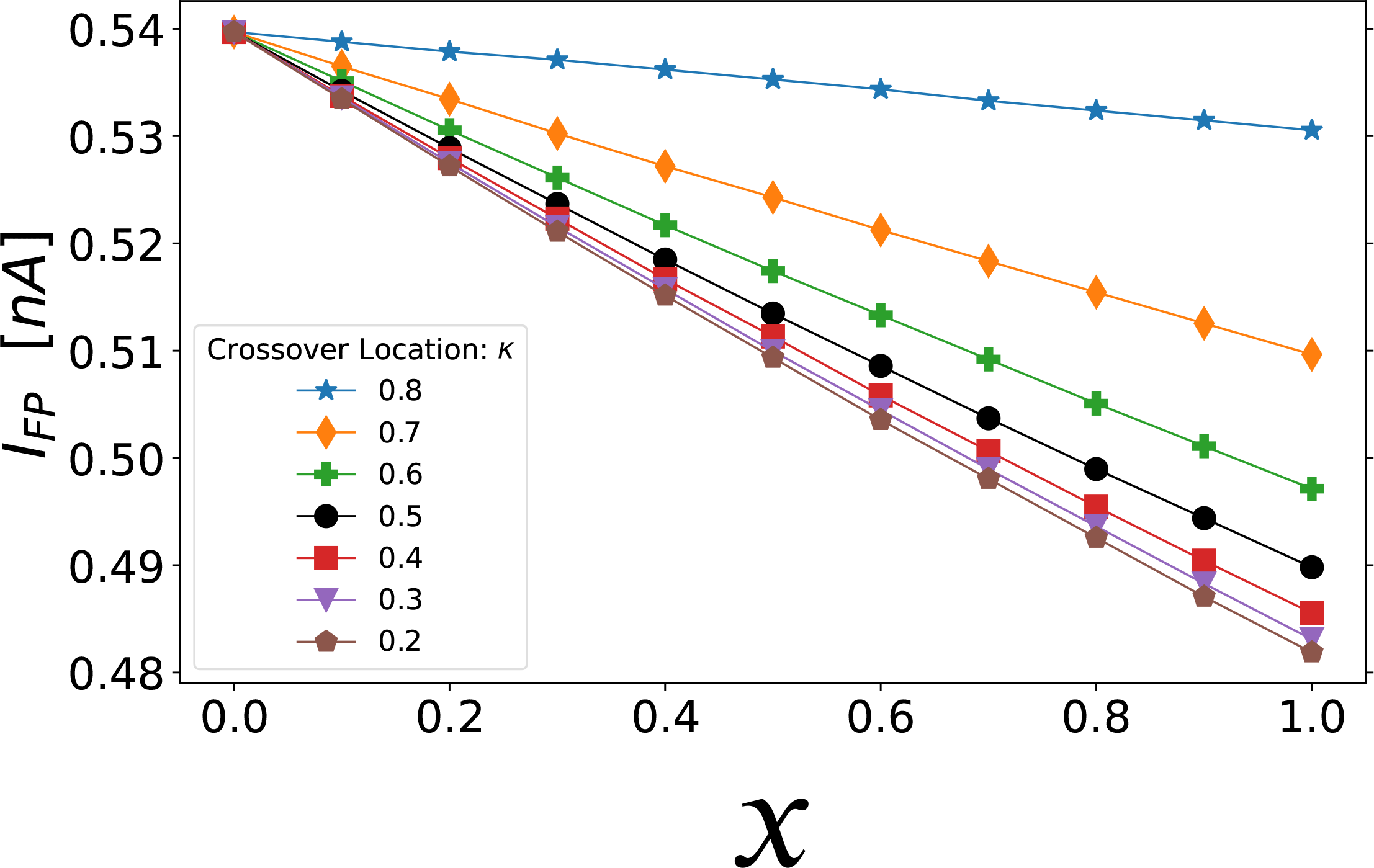
Axonal Stimulation: Effect of *x* and *κ* on forward propagation threshold. The trend for all constant *κ* curves is that raising the proportion of total AIS Na_V_1.6 (by reducing *κ*) or concentrating Na_V_1.6 in the distal AIS (by increasing *x*) lowers the threshold to initiate forward propagating action potentials (see Figure 1). Note that this threshold current pulse is not sufficient to achieve backpropagation. The effect of Na_V_ separation is much smaller here than for the backpropagation threshold. The lines have been drawn to guide the eye.

#### S1.4.1 Additional simulations

In the main text, we found that with a 25μm AIS, the slope of *I*_*BP*_ versus *x* for somatic stimulation became flat around *κ* ≈ 0.4, which is 20μm away from the soma since *L*_hillock_ = 10μm (Figure 2). In Figure S11, with a 100μm AIS, the *I*_*BP*_ slope flattens around *κ* ≈ 0.1, which again corresponds to a distance of roughly 20μm from the soma since the distance in μm to the crossover position is *κ* × *ℓ*_AIS_. This suggests that the threshold-lowering effect of Na_V_ separation for small *κ* (Figure 2) results from the increased proximal density of Na_V_1.6 when the crossover is brought near to the soma —and is not due to the proximal density of Na_V_1.2, consistent with Figure S12.

**Figure S11:**
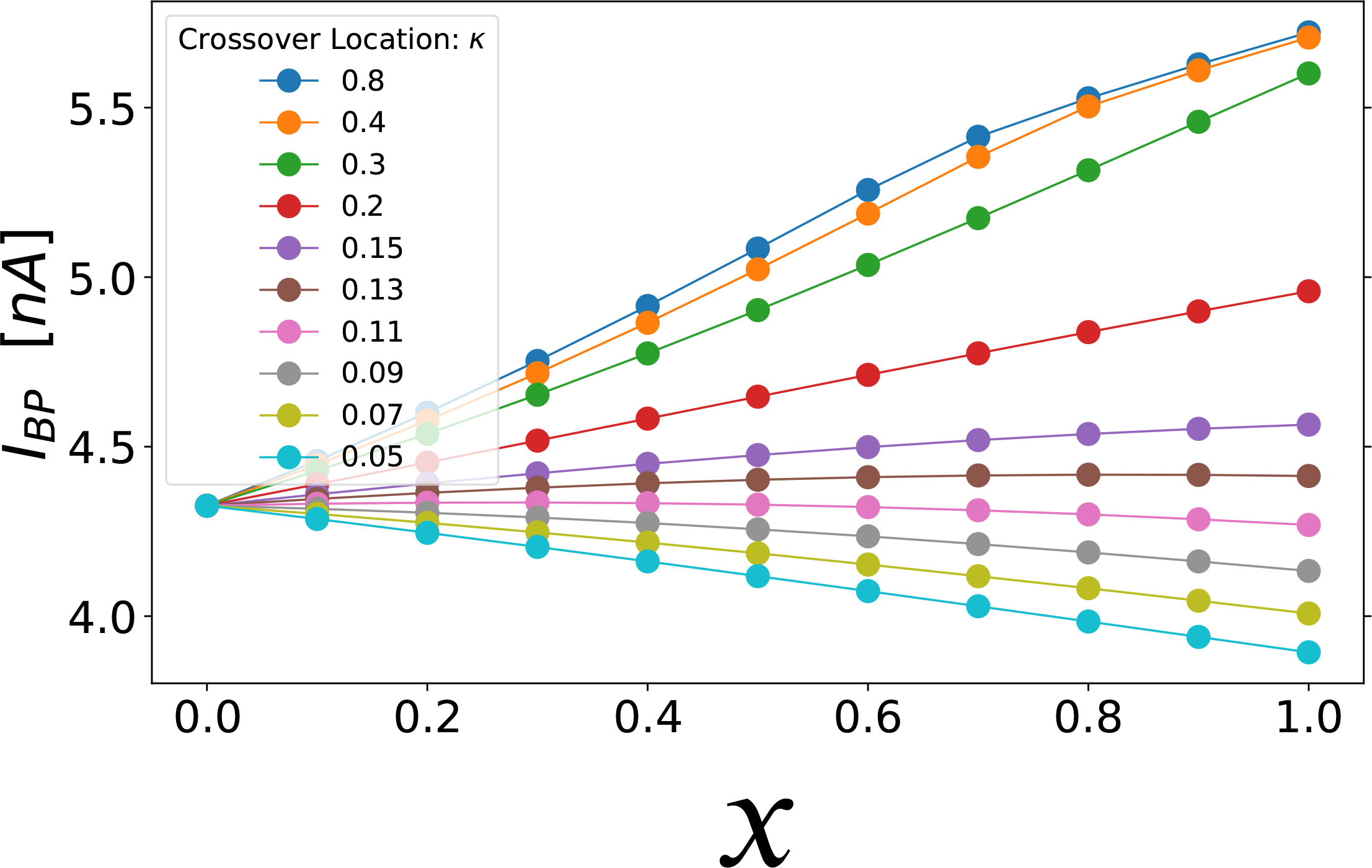
Somatic Stimulation with AIS lengthened to 100μm (compare with Figure 2 in the main text where the AIS length was 25μm): Combined effect of varying crossover location (*κ*) and Na_V_ separation (*x*) in the axon initial segment. The distance in μm to the crossover position is *κ* × *ℓ*_AIS_. The lines have been drawn to guide the eye.

Figure S12 demonstrates that increasing *κ* raises the backpropagation threshold when current is injected somatodendritically (orthodromic stimulation), even when the slopes in Figure 2 are negative. Increasing *κ* means moving the Na_V_ crossover location away from the soma, which increases the proportion of Na_V_1.2 (versus Na_V_1.6) in the AIS; see Figure 1b.

**Figure S12:**
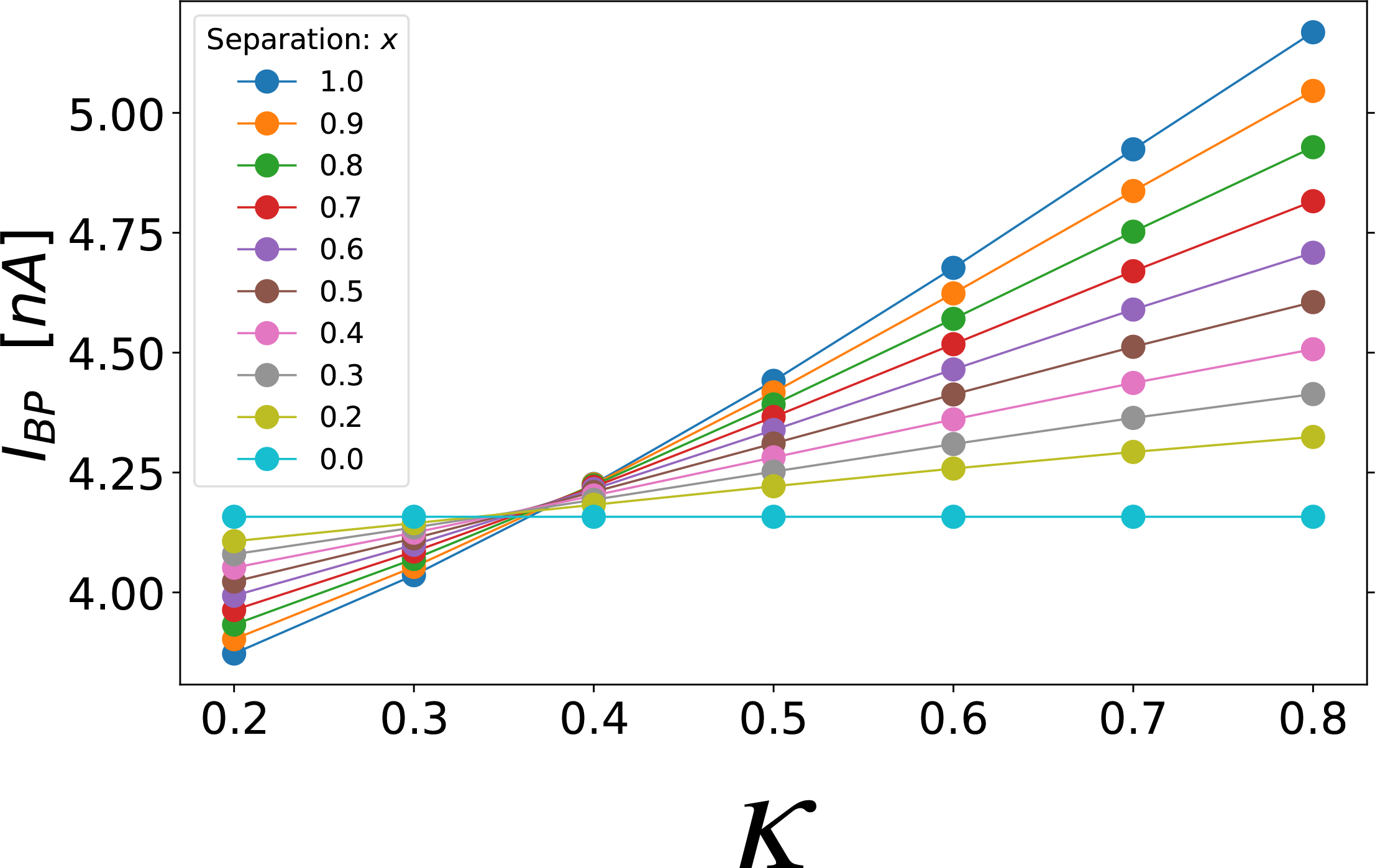
Somatic Stimulation with nominal AIS length (*ℓ*_AIS_ = 25.0μm). The combined effect of varying crossover location (*κ*) and Na_V_ separation (*x*) in the axon initial segment. Increasing *κ* raises the backpropagation threshold. Here the abscissa is the normalized crossover position *κ*, instead of Na_V_ separation (compare with Figure 2). In Figure 2, all curves converge at *x* = 0. Here, that intersection point is replaced by the *x* = 0 *line*. Notice that every *x* > 0 curve has a positive slope: the backpropagation threshold *I*_*BP*_ *increases* with *κ*. Increasing *κ* when *x* > 0 necessarily increases the ratio of Na_V_1.2 conductance to Na_V_1.6 conductance in the AIS (see Equation S1). It follows that concentrating Na_V_1.2 in the proximal AIS raises the backpropagation threshold, for somatic stimulation. The lines have been drawn to guide the eye.

**Figure S13:**
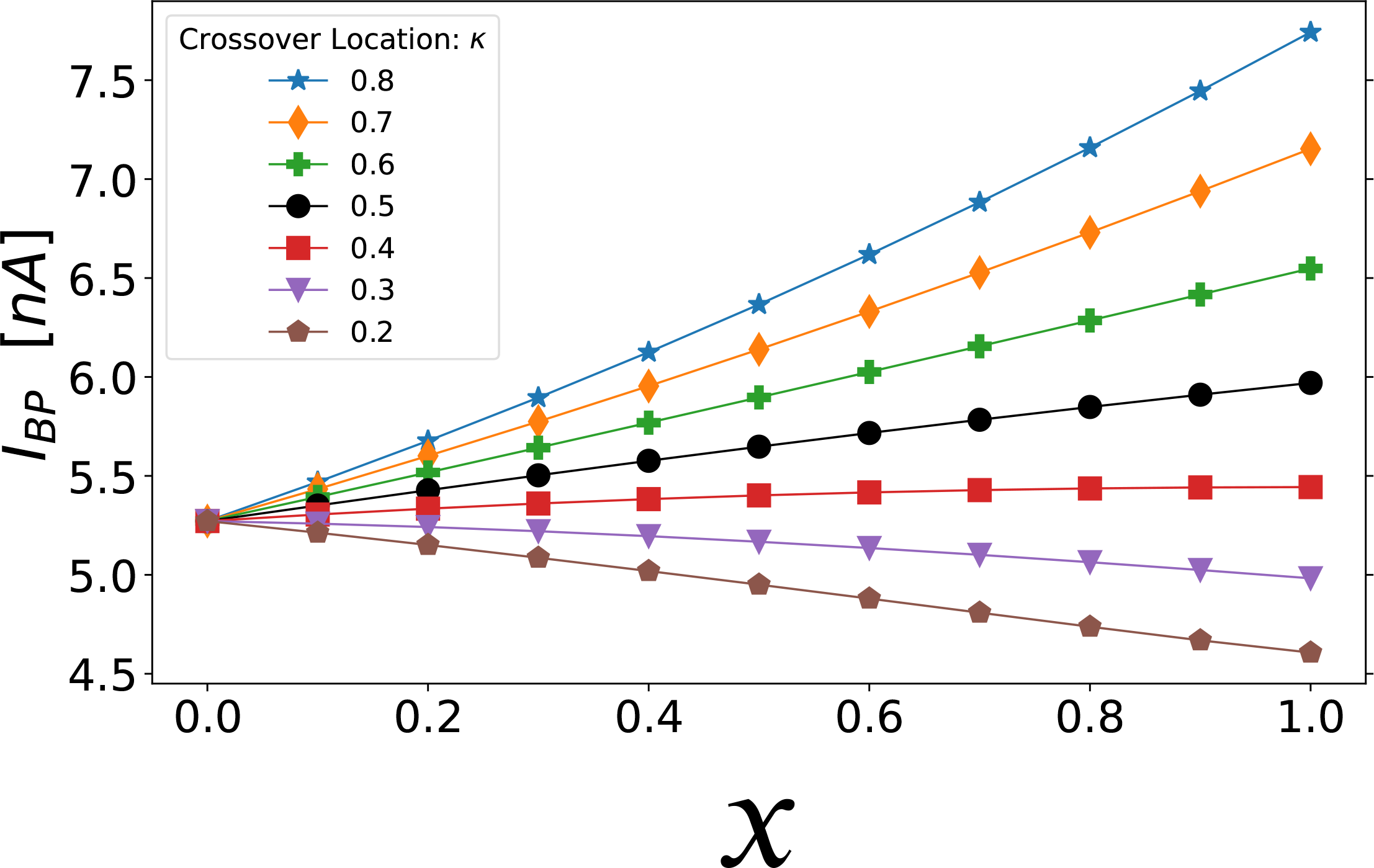
Dendritic Stimulation with nominal AIS length (*ℓ*_AIS_ = 25.0μm). Current injection at the main apical dendrite yields the same qualitative behaviour as for somatic stimulation. Compare with (Figure 2). Varying the separation parameter “*x*” from *x* = 0 to *x* = 1, the distribution of Na_V_ channels goes from flat (homogeneous) to separated, the latter approximating the distribution observed in developing pyramidal neurons (see Figure 1a). Note that curves for all values of *κ* converge to a single point at *x* = 0 since *κ* can have no effect when the two Na_V_ subtypes are uniformly distributed along the AIS. The lines have been drawn to guide the eye.

**Figure S14:**
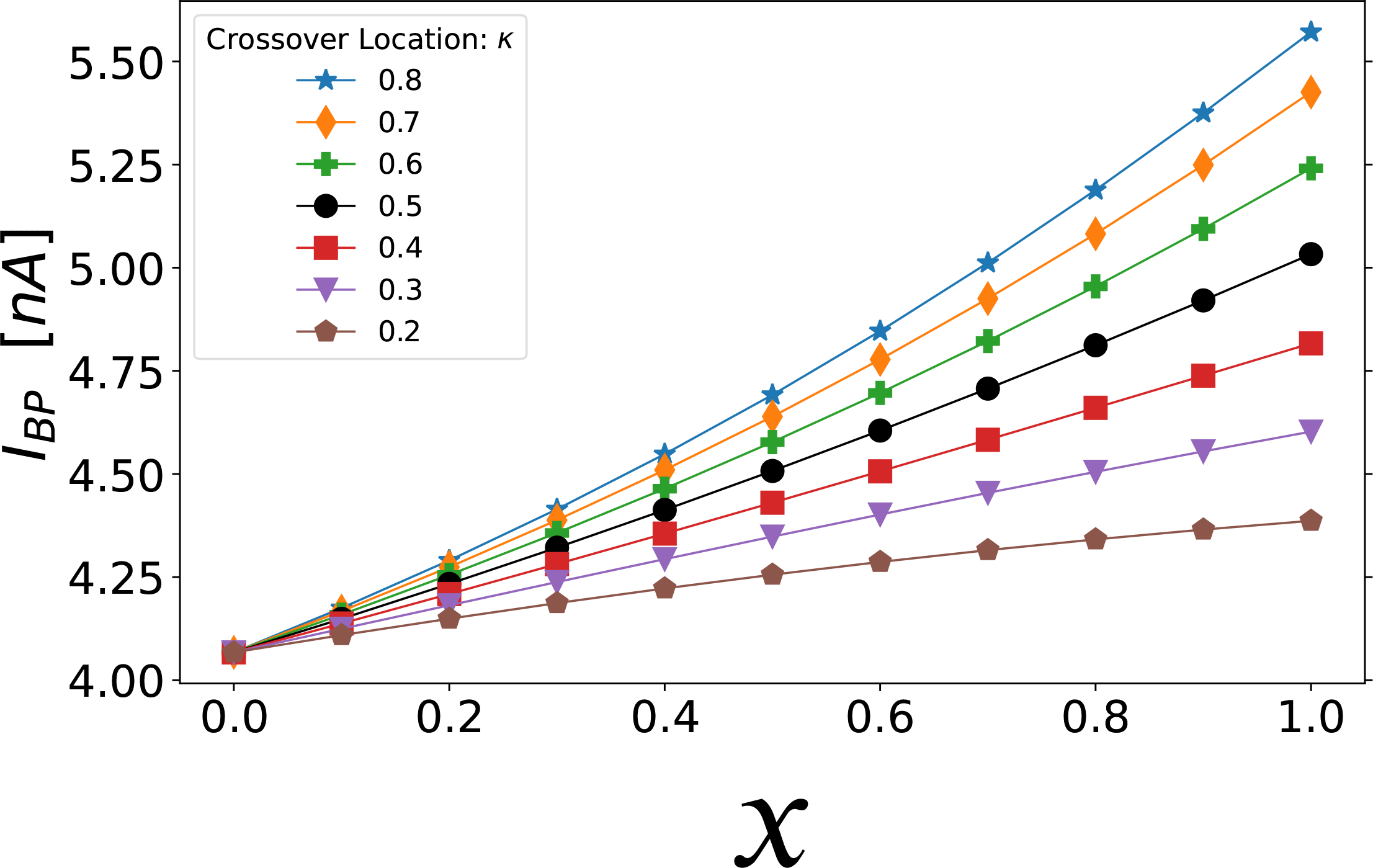
Somatic Stimulation: lengthening the hillock to 30μm removes the negative slopes observed in Figure 2 and Figure S11. In all other plots, we have used *L*_hillock_ = 10μm. AIS length is *ℓ*_AIS_ = 25.0μm as in all other plots unless indicated otherwise. The lines have been drawn to guide the eye.

### S1.5 AIS - technical details

In our model, the density profiles of Na_V_1.2 and Na_V_1.6 are left- and right-handed sigmoidal functions (respectively) of normalized length *s* along the AIS. The proximal end of the AIS is located at *s* = 0, and the distal end is located at *s* = 1. The channel densities are expressed as maximal conductances 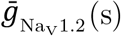 and 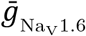 (s), where the total maximal Na_V_ conductance 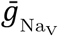 is constant along the AIS:

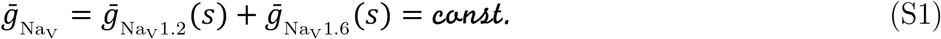

The density profiles are given by

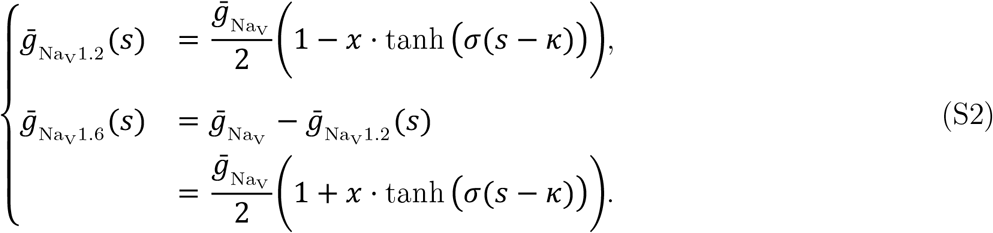

We chose the hyperbolic tangent function tanh(*s*), but other sigmoidal functions would do just as well. The parameter *x* controls the separation of the Na_V_ distribution, that is, how separated the two Na_V_ subtypes are. When *x* = 0, the distribution becomes flat —Na_V_1.2 and Na_V_1.6 are mixed uniformly along the AIS. When *x* = 1, the proximal end of the AIS contains only Na_V_1.2, and the distal end of the AIS contains only Na_V_1.6. The parameter *σ* is the reciprocal of the ‘transition width’ of the AIS Na_V_ distributions normalized by the AIS length. In all simulations shown here, *σ* = 10.0.

The total Na_V_ conductance of the AIS is proportional to AIS length *ℓ* since its diameter is constant

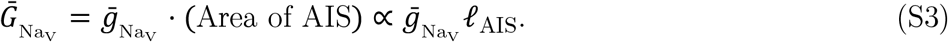

The contribution to this total conductance from Na_V_1.2 is

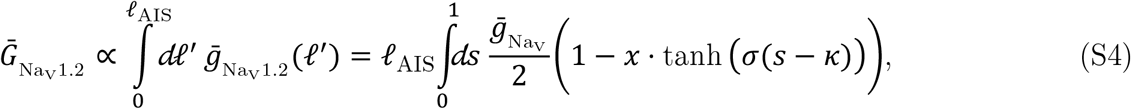

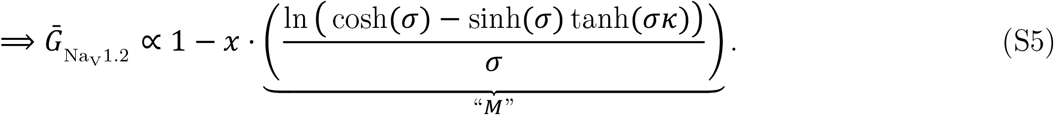

The root of the term labeled “*M*” in Equation S5 is 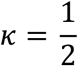, and the slope of *M* is negative: *M*(*κ* < 0.5) > 0, *M*(*κ* > 0.5) < 0. Since *M*(*κ* = 0.5) = 0, the derivative of 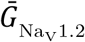 (and 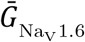) with respect to *x* is zero when the crossover is located in the middle of the AIS, which is the standard configuration for varying Na_V_ separation. We call *κ* = 0.5 standard because, in this configuration, the effects on the backpropagation threshold due to varying *x* can not be due to changes in the total conductance of Na_V_1.6 or Na_V_1.2 in the AIS. In other words, the results of sweeping *x* from 0 to 1 with *κ* fixed at 0.5 in Figure 4, Figure 5, and Figure 2 are purely due to mixing and separating Na_V_1.6 from Na_V_1.2 in the AIS.

Since by definition 0 *⩽ κ ⩽* 1 and *σ* > 0, the partial derivative of *M* with respect to *κ* is negative. It follows from Equation S5 that 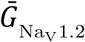 increases as the crossover position is moved distally. Likewise 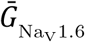 decreases with increasing *κ*.

### S1.6 Voltage-gated channels

The voltage-gated sodium and potassium conductances 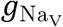 (*V, t*) and 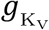 (*V, t*) in Equation 15 are modeled using HH-style kinetics [40] fitted to mammalian pyramidal cell data by [11], and then further adapted by [14] to include two Na_V_ variants (Equation S7). In the Hodgkin-Huxley model, the current density *I*_Z_ of ion species “Z” through voltage-gated channels of a given type is

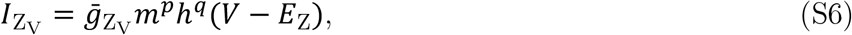

where *g* is the maximal conductance, *m*(*V, t*) is the probability for an *activation* gate to be open, *p* is the number of *activation* gates per channel, *h*(*V, t*) is the *availability* (probability that the channel is *not inactivated*), and *q* is the number of inactivation gates per channel. Na_V_ channels are modeled as having three *activation* gates (*p* = 3) and a single inactivation gate (*q* = 1) so that 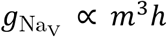 ∝ *m*^3^*h*. Likewise, K_V_ channels have a single *activation* gate and no inactivation (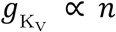). The gating variables *m, h*, and *n* evolve according to Equations 6, 7, and 8. Since the cell features two Na_V_ subtypes (Na_V_1.2 and Na_V_1.6), the model computes two sets of sodium *activation* and *availability* variables. The current density *I*_*Na*_ through the Na_V_ channels is then

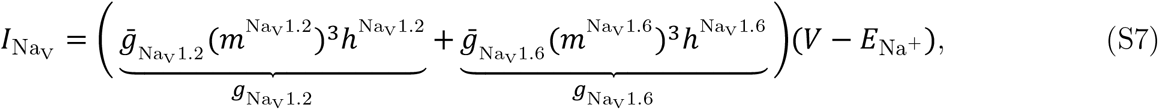

where 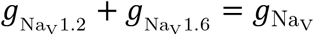 in Equation 15.

Figure S15 plots the voltage-dependent kinetics of 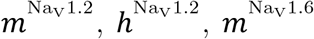 and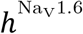. The steady-state *activation* and *availability* functions and their voltage-sensitive time constants were implemented using the parameters provided in [14] and in model code published by [24]. We further modified the channel models to allow shift-clamping; see Section 2.5 and Section 4.2.

**Figure S15:**
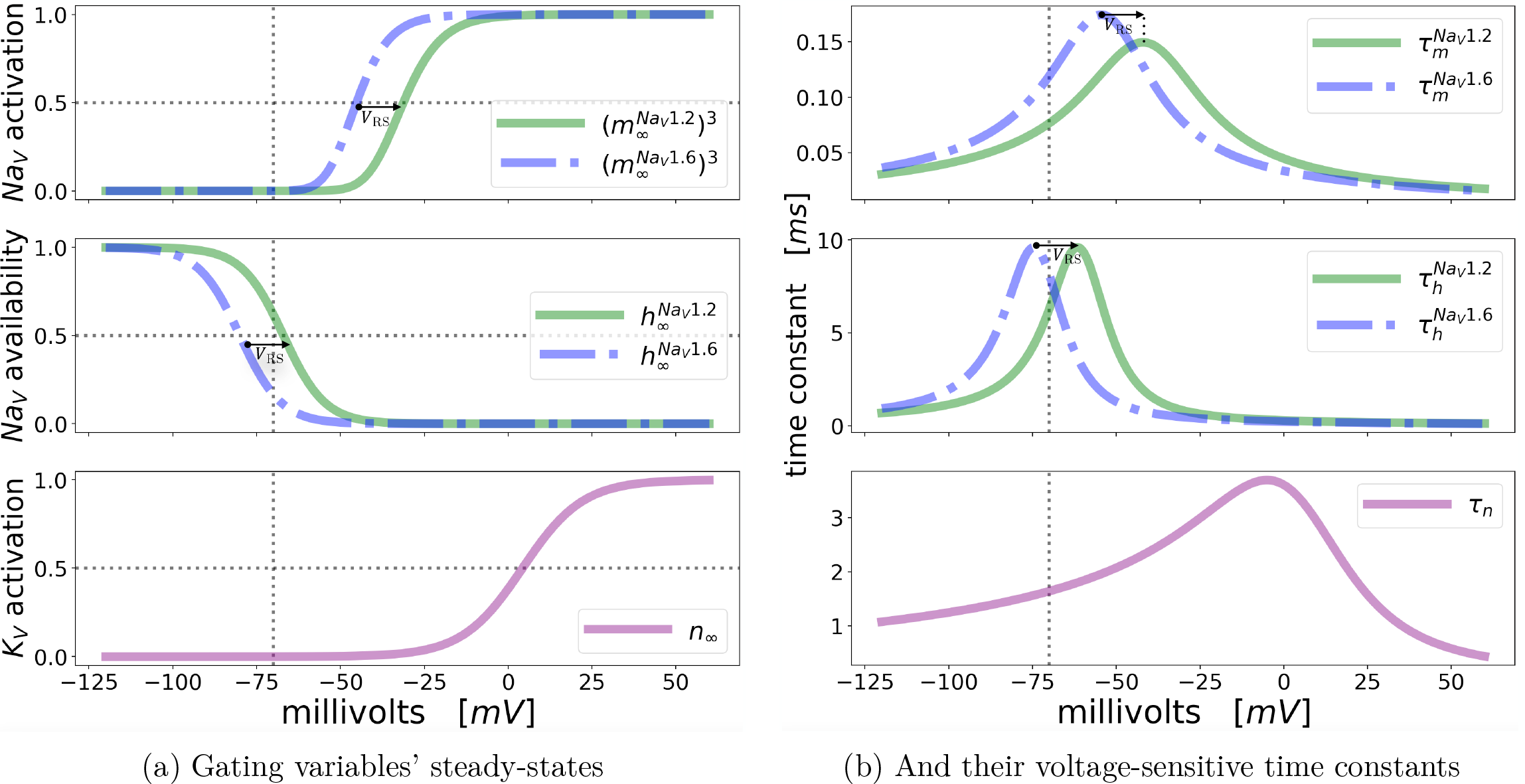
Gating properties of the voltage-gated sodium and potassium channels that are implemented in this model. Dotted vertical lines indicate the resting potential *V*_rest_. **(a)** Steady-state *activation* and *availability* curves for Na_V_1.2 and Na_V_1.6, and steady-state K_V_ *activation*. **(b)** Voltage-sensitive time constants of Na_V_1.2, Na_V_1.6, and K_V_. Na_V_1.2 is *right-shift*ed by an amount *V*_RS_ relative to Na_V_1.6 (defined in Section S1.6.1). In this model, *V*_RS_ = 13.0mV; also called the ‘nominal *right-shift*’ of Na_V_1.2. *V*_RS_ is **indicated by arrows** (small 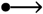) in the plots of Na_V_ steady-states **(a)** and time constants **(b)** above.

*V*_RS_ is indicated by arrows (small 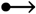) in the plots of Na_V_ steady-states (Figure S15a) and time constants (Figure S15b).

The *right-shift* of Na_V_1.2 (defined mathematically in Section S1.6.1) is easiest to observe in the top two plots of Figure S15a. *V*_RS_ is indicated by arrows (small 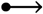). Steady-state *activation*.(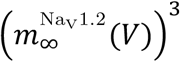) and *availability* 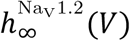 (*V*) curves of Na_V_ 1.2 are plotted as solid green lines, and dashed blue lines are the corresponding curves for Na_V_1.6. The voltage separating the two Na_V_ subtypes’ *activation* curves is approximately the nominal *right-shift, V*_RS_; however, it varies with position, since the gating variables of Na_V_ *activation* (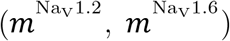) have different slopes [14]. However, the *availability* curves are easier to compare: in this model, 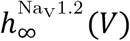 (*V*) is shifted exactly *V*_RS_ = 13.0mV to the right of 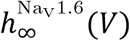 (*V*). The *right-shift* of Na_V_1.2 is also visible in the voltage-sensitive time constants (Figure S15b).

#### S1.6.1 Defining *V*_RS_: the *right-shift* of Na_V_1.2

*V*_RS_ is a parameter in this model representing the experimentally measured depolarizing shift in the voltage dependence of Na_V_1.2 *activation* and inactivation kinetics, relative to Na_V_1.6 kinetics. Because *V*_RS_ is empirical, it is fixed in this paper.

We define the Na_V_1.6 half-*activation* voltage 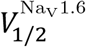 as the voltage at which a single Na_V_1.6 *activation* gate (randomly selected from an ensemble of gates held at 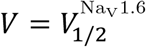) has a 50% chance of being in the open state:

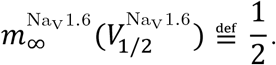

Although the kinetics of Na_V_1.2 differ from Na_V_1.6 kinetics in ways *other than right-shift*, we can now use 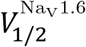 to define *V*_RS_ as the voltage that satisfies

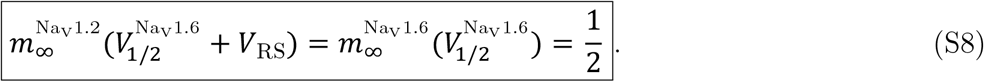

Note that 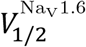 and *V*_RS_ are unique since *m*_∞_ is monotonically increasing. Equation S8 also contains the half-*activation* voltage for Na_V_1.2,

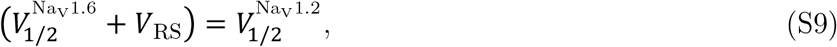

which is depolarized or “*right-shift*ed” by an amount *V*_RS_ relative to Na_V_1.6 (see Figure S15).

To simulate alterations to Na_V_1.2 *right-shift* in our shift-clamping method, we use another parameter called Δ*V*_RS_, which is not based on experiment. Although the Na_V_1.2 *right-shift* is not a high precision measurement (*V*_RS_ ∼ 10−15mV), the *parameter V*_RS_ is kept fixed in our model for conceptual purposes: we find it helpful to distinguish empirical parameters (like *V*_RS_) from exploratory parameters that intentionally deviate from experiment (like Δ*V*_RS_). In fact, we use Δ*V*_RS_ to selectively change the model’s Na_V_ kinetics to explain the effects of Na_V_ distribution on excitability in terms of the lengthwise distribution of gating properties (see Section 2.5 and Section 4.2).

#### S1.6.2 Notation: *V*_RS_, Δ*V*_RS_

In our notation, *V*_RS_ is *not written explicitly* in the argument of Na_V_1.2 gating variables or their time constants. Instead, we write

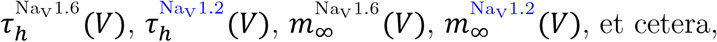

and let the superscript 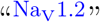 indicate that 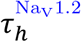 is *right-shift*ed relative to 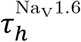, from the fact that Na_V_1.2 channels are *right-shift*ed in this model.

However, the parameter Δ*V*_RS_ *is* written explicitly in the argument when we model the effects of modifying the *right-shift* (Figure 6: shift-clamping). For example, when applying Δ*V*_RS_ ≠ 0 to the *selected* gating properties 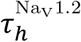 and 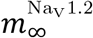, we would write

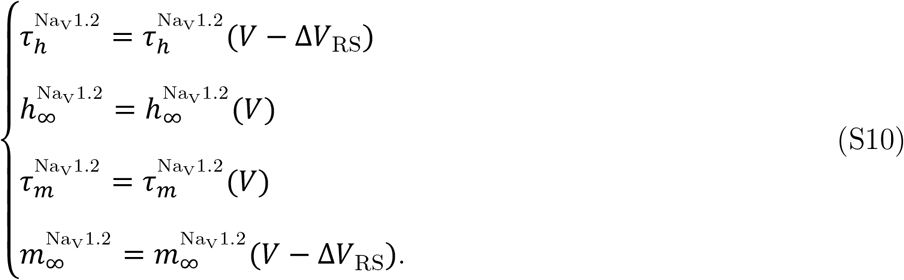

Using this notation, positive values of Δ*V*_RS_ will shift kinetic curves (gating properties) to the right in Figure S15 (depolarizing shift), and negative Δ*V*_RS_ produces a hyperpolarizing shift.

#### S1.6.3 Space plots of Na_V_ kinetics along the AIS — steady-state

The distribution of Na_V_1.2 and Na_V_1.6 in the AIS creates a lengthwise distribution of gating properties. Proximal Na_V_1.2 increases local steady-state *availability*, owing to these channels’ *right-shift* (*V*_RS_). In Figure S16a we visualize this effect using *ℌ*(*s*): the net *availability* of Na_V_s (*ℌ*) as a function of position (*s*), computed by weighting 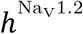 and 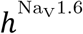 according to their respective local Na_V_ channel densities:

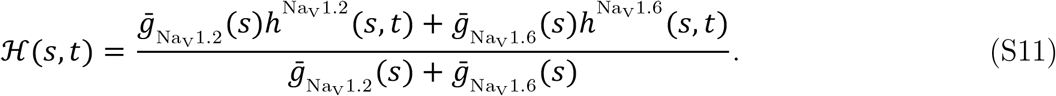

An effective local Na_V_ time constant of *availability* (𝒯) can be computed from 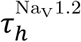and 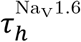 as

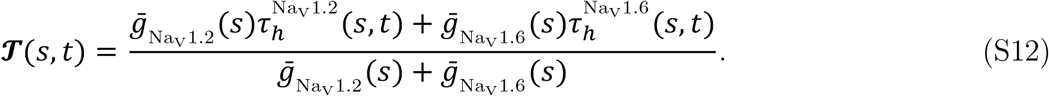

Above, we have abbreviated

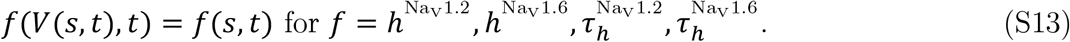

In Figure S16b we have selectively disabled the *right-shift* of 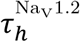 by setting Δ*V*_RS_ = −*V*_RS_ = −13.0mV in Equation 2, which leaves the *right-shift* of steady-state *availability* unchanged.

**Figure S16:**
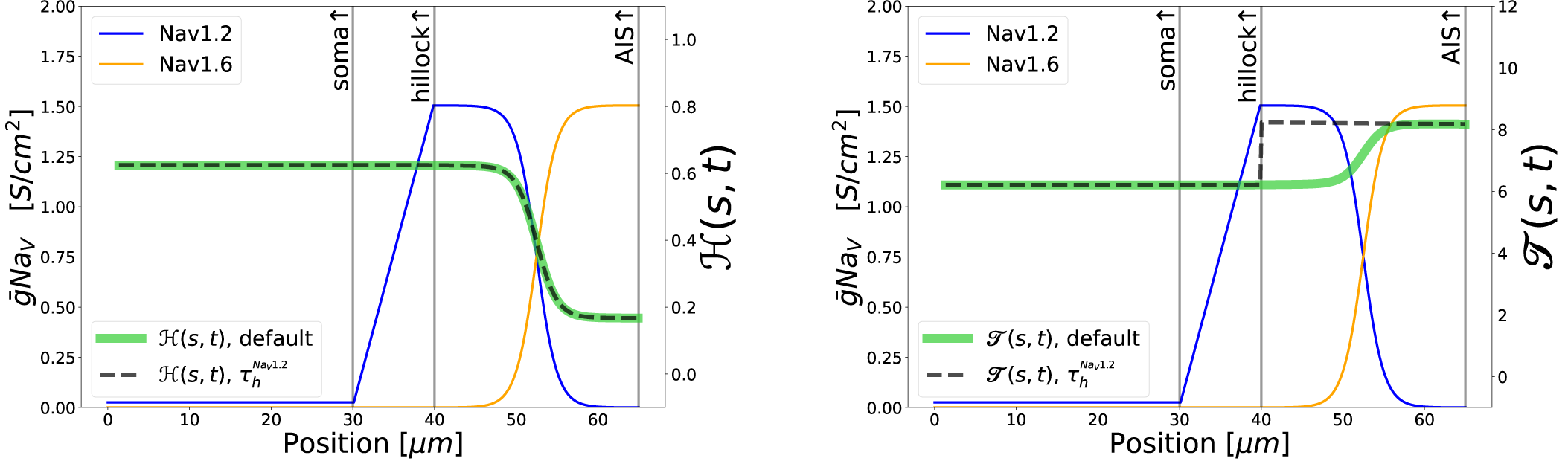
The effect of Na_V_1.2 *right-shift* on local Na_V_ *availability* along the cell is visible in space plots of steady-state total *availability ℌ*(*s*) (a) and total time constant *𝒯*(*s*) (b) at *V* = *V*_rest_. See Equation S11 and Equation S12, respectively.

### S1.7 Diffusion coefficients

Diffusion coefficients for Na^+^, K^+^, and Cl^-^ in water at 25.0 ° are provided by [46] :

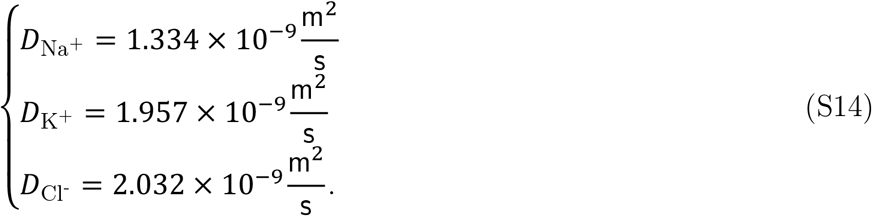

Our simulations are done at a warmer temperature, so these coefficients need to be adjusted. We make the adjustment using the Stokes-Einstein equation

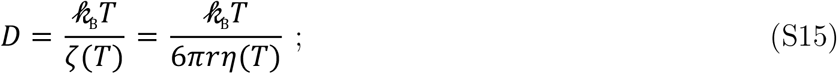

where 𝓀_B_ is Boltzmann’s constant, *T* is the temperature (Kelvins K), and ζ is called the drag coefficient. ζ is given by the ion’s radius *r* and the viscosity *η*(*T*) of the medium (liquid water), which depends on temperature. Hence the ratio of diffusion coefficients for ion species Z at *T*_2_ and *T*_1_ is

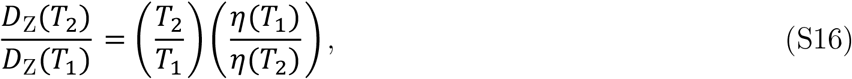

with *T*_2_ and *T*_1_ converted to K. The reference values provided by [46] are measured at 298.15K (25.0°), and the simulation temperature is 310.15K (37.0°). Assuming the viscosity of water, we have *η*(*T*_1_) ≅ 0.89 mPa s and *η*(*T*_2_) ≅ 0.691 mPa s. Substituting these *η*’s into Equation S16 gives for each ionic species Z

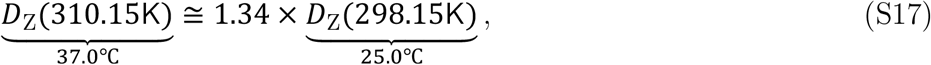

which yields the temperature-adjusted diffusion coefficients

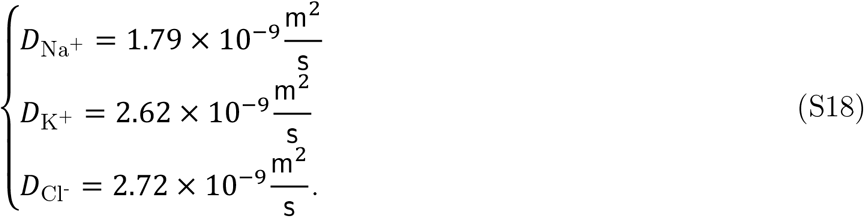

### S1.8 Tables of parameters

In the following tables, we compare our model to previous models on which it is based ([11], [24], [14]). At different locations in the cell, we list key parameter values from these papers alongside our own. We also include certain measurements (membrane potential, ion concentrations, Nernst potentials) taken from our model at each location once the cell has equilibrated.

The following symbols are used in the tables: (Units are given in the tables.)

**“**×**”** This symbol appears when a parameter is not featured in the model corresponding to a given column.

*V*_**rest**_ Resting potential. The transmembrane voltage of the model neuron at steady-state with no injected current.

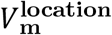 Transmembrane voltage at ‘location’. In our model, the steady-state transmembrane voltage is actively maintained everywhere by Na^+^/K^+^-pumps and longitudinal diffusion, which control the explicit intracellular and extracellular ion concentrations. As such, the tabulated values are *recorded* from the model, not parameters per se. In the other models listed, *V*_m_ is identical to *V*_rest_.

*R*_**axial**_ Axial resistance. The resistance to axial current flow across a compartment is *R*_axial_ multiplied by the compartment length, divided by the compartment’s cross-sectional area.

*D*_**ion**_ Diffusion coefficient of the specified ion in water at 37°.

*T*_**ref**_ Reference temperature of experimentally developed channel properties used in the models. Past modelling, on which this paper is based, used temperature factors (described below) to adjust channel densities and speed up channel kinetics to warmer temperatures.

*T* Simulation temperature.

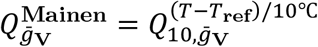 To adjust for a warmer simulation temperature of *T* = 37° [24] scales up the maximal voltage-gated conductances, which were originally developed at *T* = 23° ([11]), by a factor which we denote 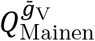. The authors warn in their model code that this scaling is valid only at 37° and state that their program is not designed to model other temperatures. This internal temperature scaling can accidentally obscure parameter settings when one attempts to borrow parameters separately from the past code or from later papers that reuse these channel models. For this reason, we have removed the temperature scaling from our own model code by setting our reference temperature equal to the simulation temperature, which sets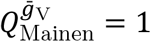.

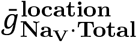 Total combined maximal voltage-gated conductance density of Na_V_1.2 and Na_V_1.6 at the specified ‘location’ in the cell (see Equation S1). The location can be an entire Section if the membrane properties are uniform (in NEURON, Sections consist of multiple compartments). For example, this model has uniform somatodendritic channel densities, so ‘soma’ is sufficient to specify those parameters.

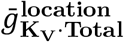 Maximal voltage-gated K^+^ conductance density at the specified ‘location’.

*C*_**m**_ Membrane capacitance per unit area. Applies everywhere except at internodes.

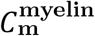 Membrane capacitance of myelinated internodes.

*L*_**Section**_ Length of the specified ‘Section’ (soma, hillock, AIS, etc.)

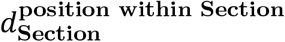 Diameter at a given normalized position (0 *⩽ s ⩽* 1) within the specified Section (soma, hillock, AIS, etc.). The position is not specified when the diameter is uniform.

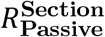 Membrane resistivity in ‘Section’.

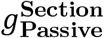 Passive generic transmembrane leak conductance density in ‘Section’.

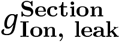 Specific leak conductance density of ‘Ion’ in ‘Section’.

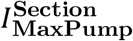 Maximum Na^+^/K^+^-pump current density in ‘Section’.

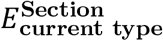 Nernst Potential (reversal potential) of transmembrane current ‘current type’ (e.g. ‘leak’) in ‘Section’ (e.g. ‘soma’).

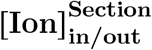 Intracellular (‘in’) or Extracellular (‘out’) concentration of ‘Ion’ at ‘Section’, with the neuron at steady-state. Because these concentrations are maintained by Na^+^/K^+^-pumps and longitudinal diffusion in our model, the concentrations tabulated here are measurements *recorded* from the simulation at steady-state, rather than being fixed parameters.

**ranvier0, myelin0:** In our model, there are 15 nodes of Ranvier and 16 myelinated internodes. The Section named ‘ranvier0’ is the first node of Ranvier. And ‘myelin0’ is the first myelinated internode of the axon, located between the distal end of the AIS (or the distal end of the bare axon when that is included) and the proximal end of ranvier0.

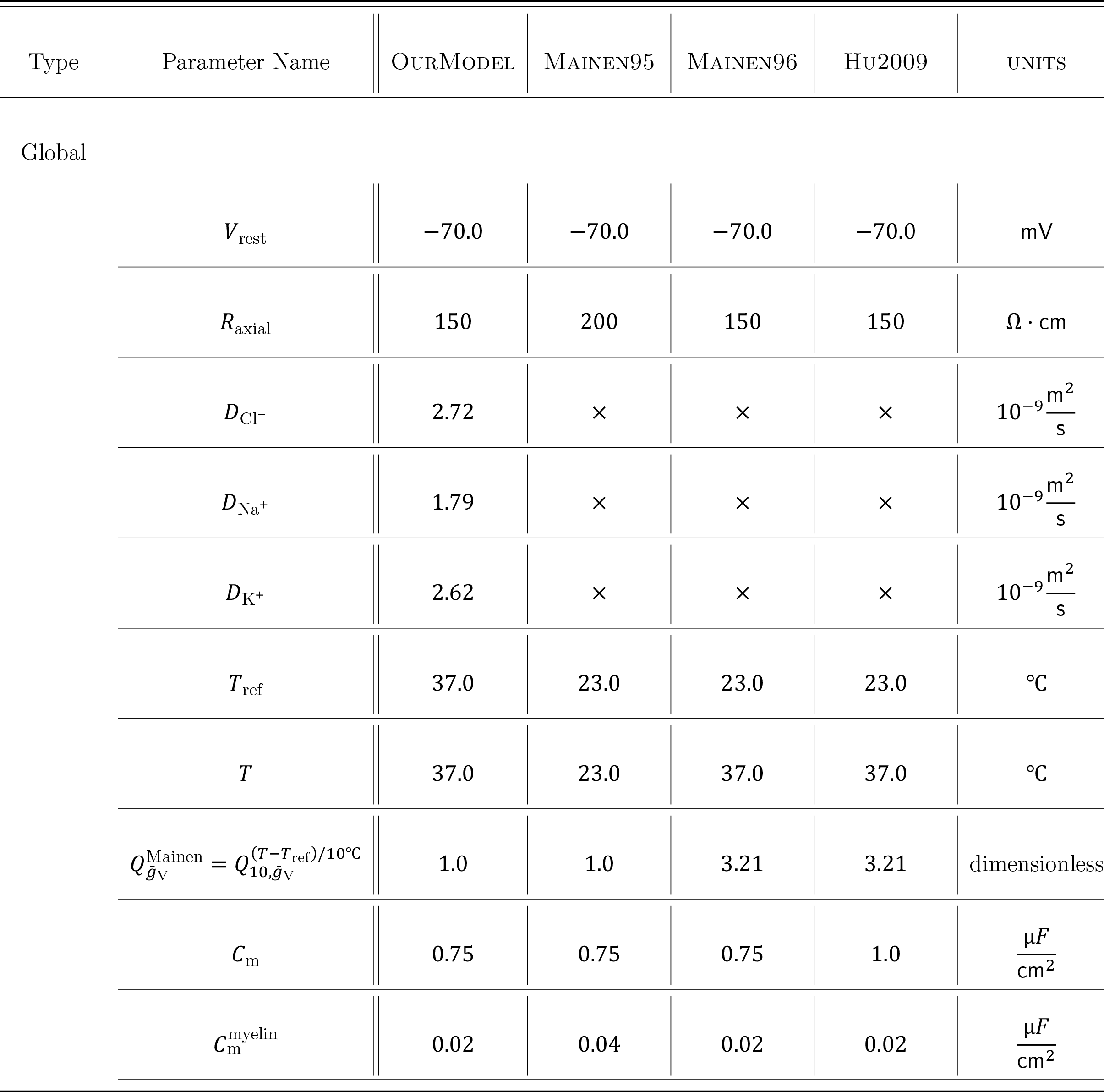
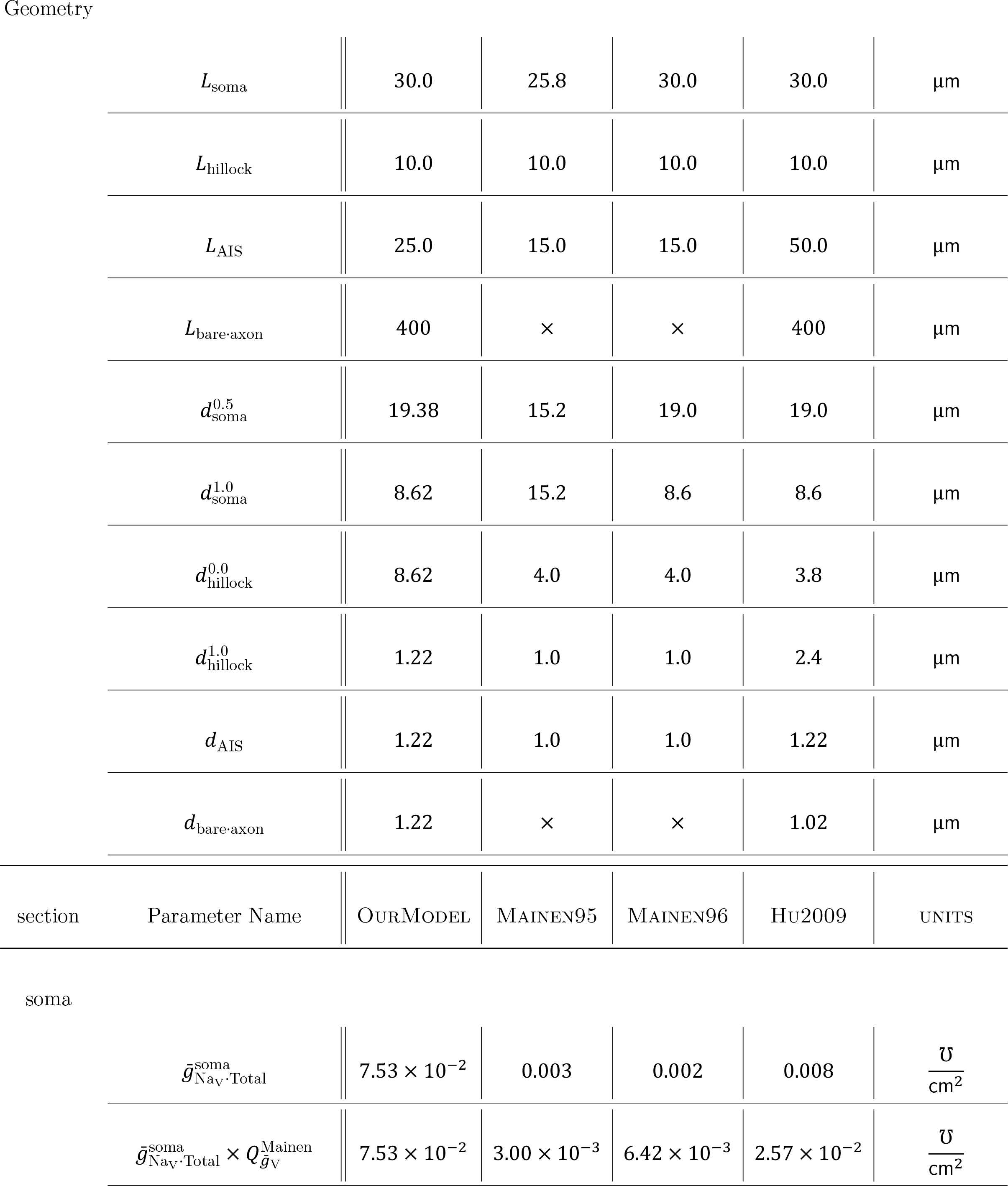
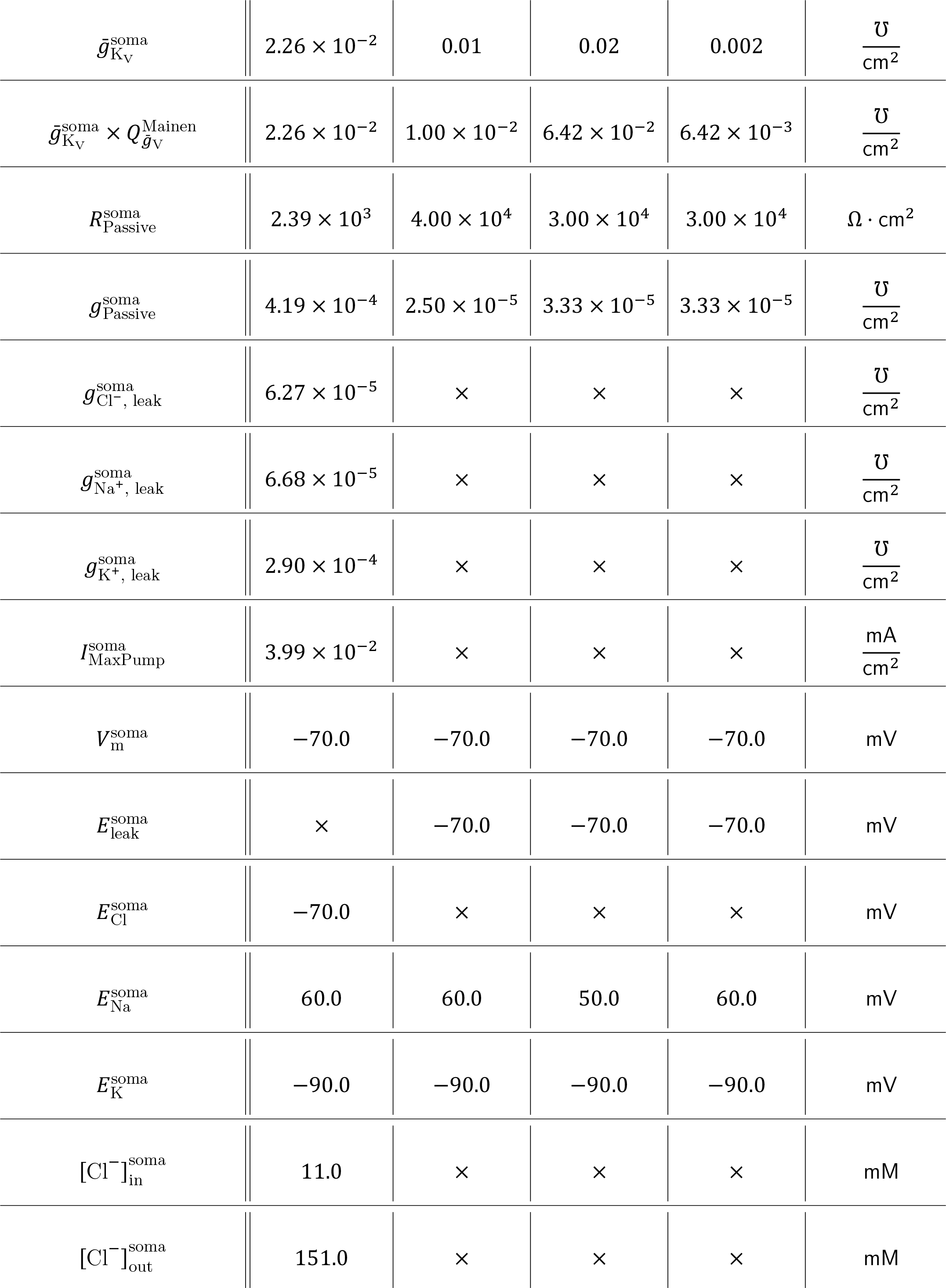
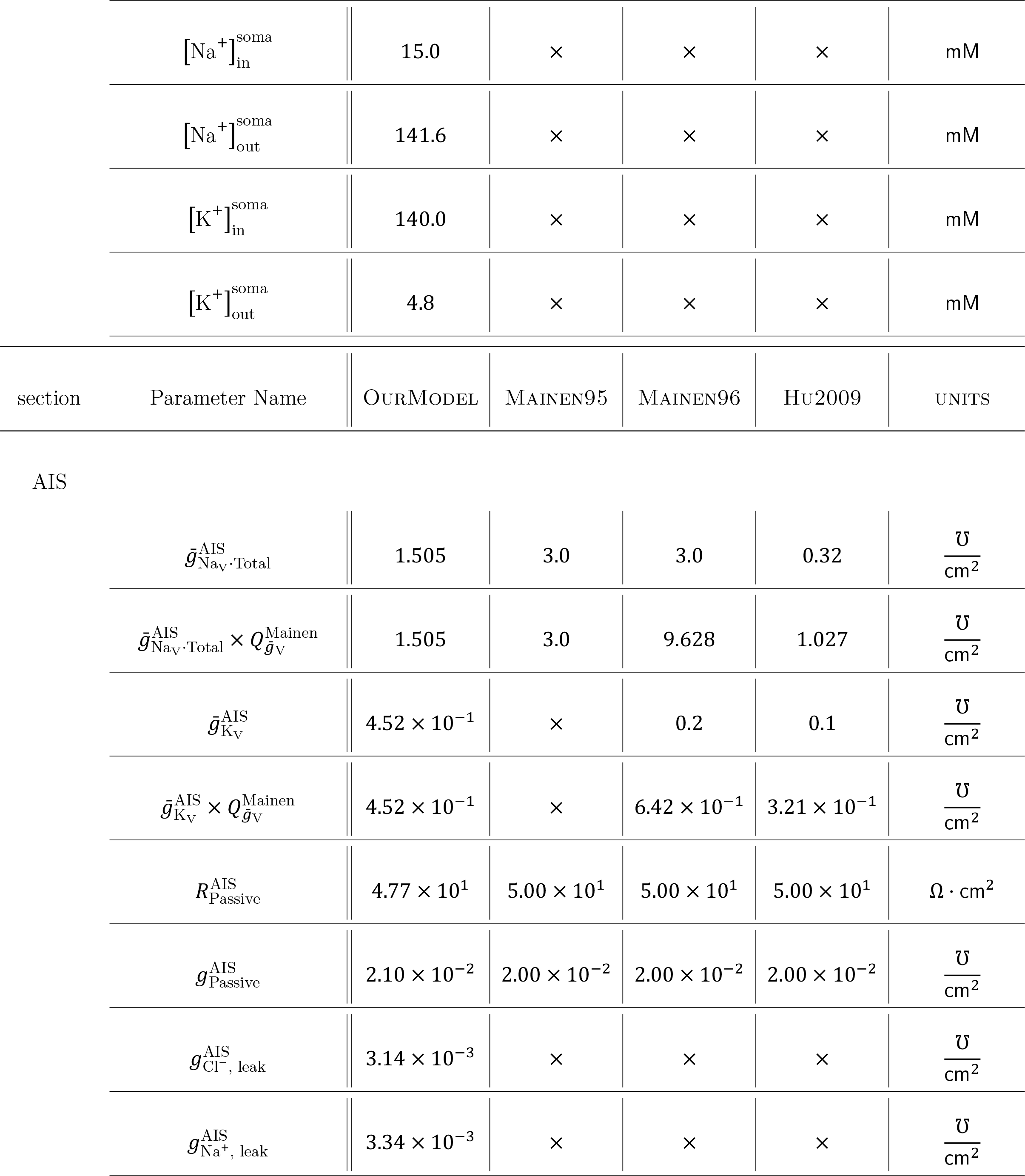
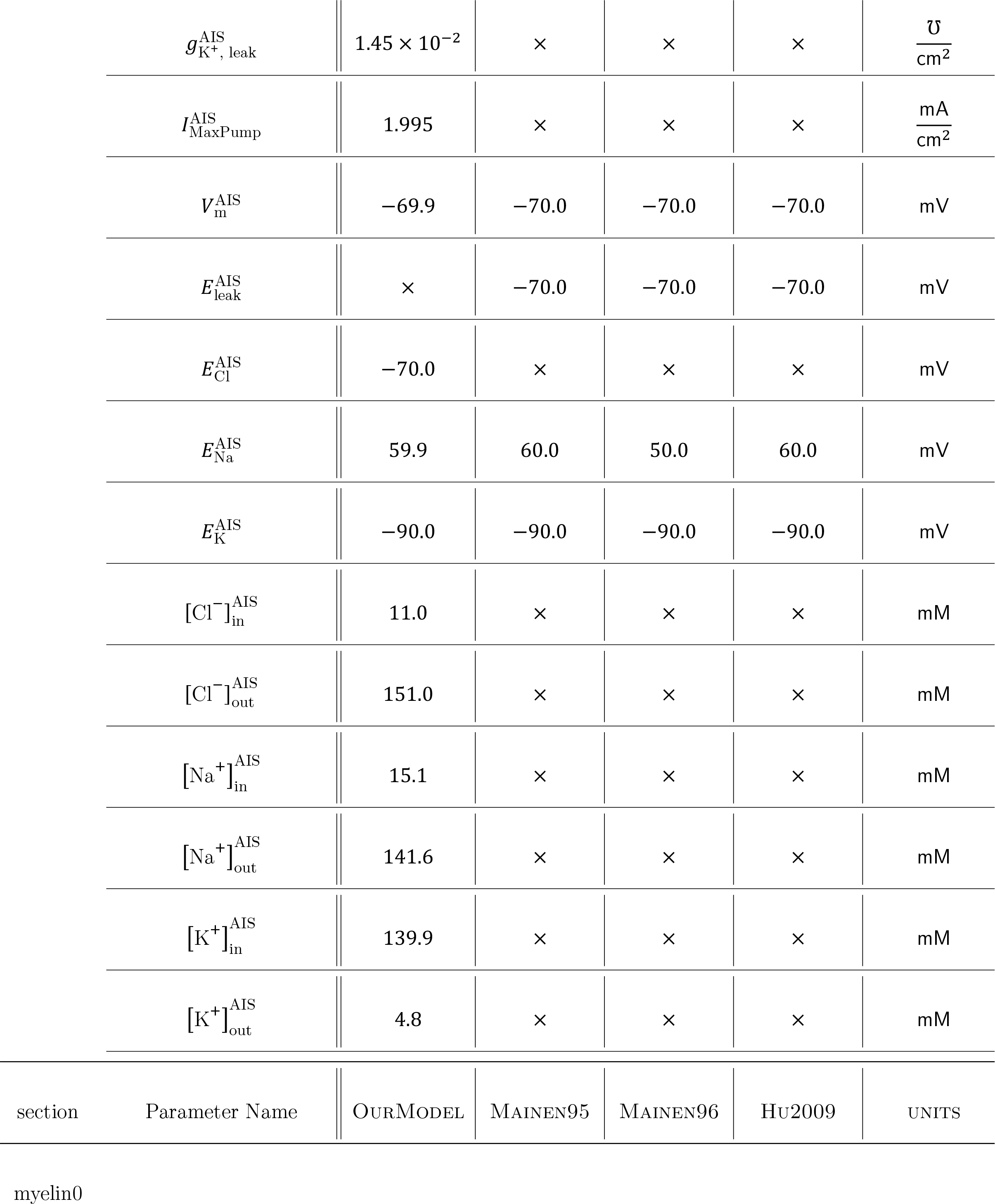
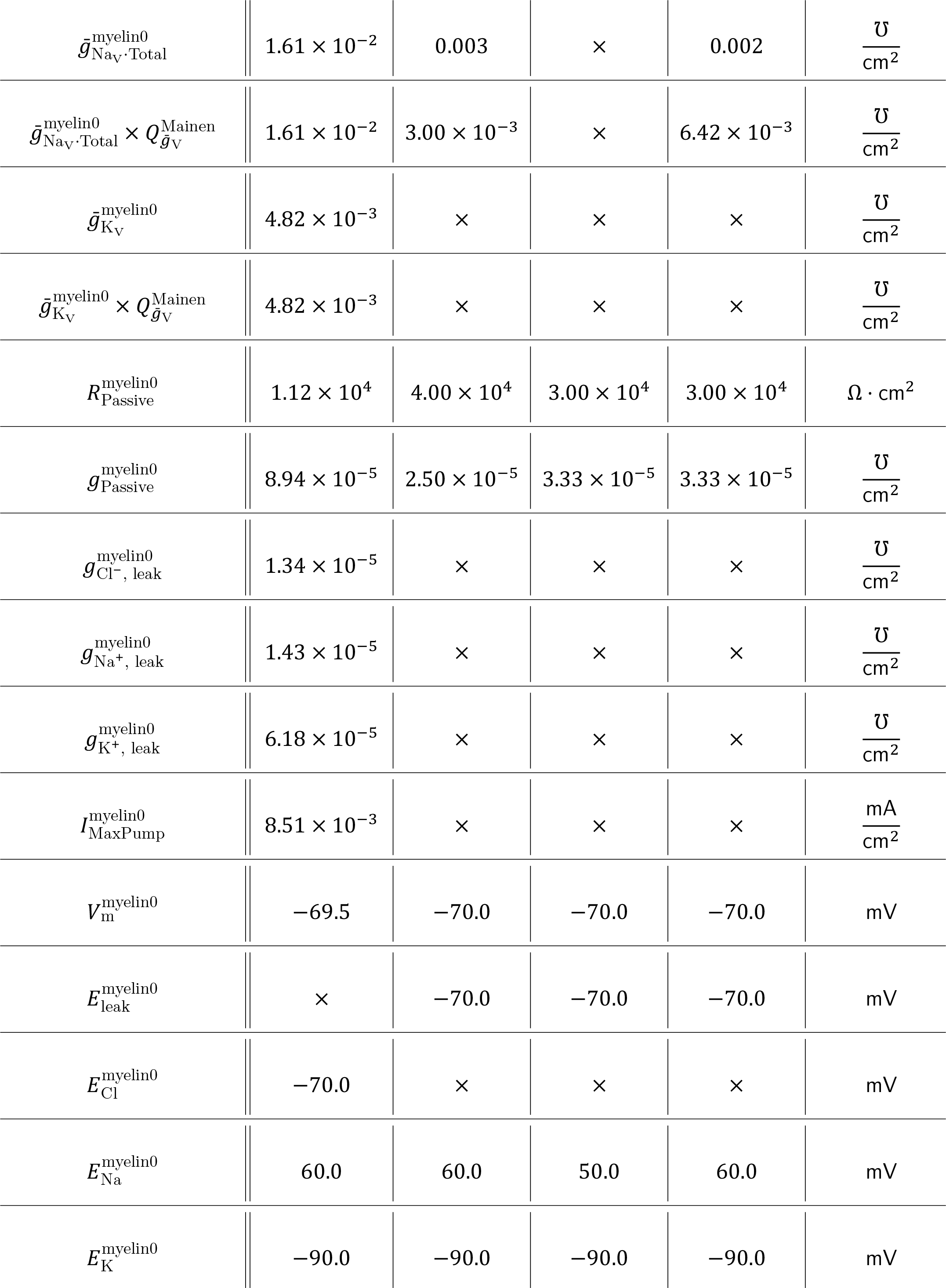
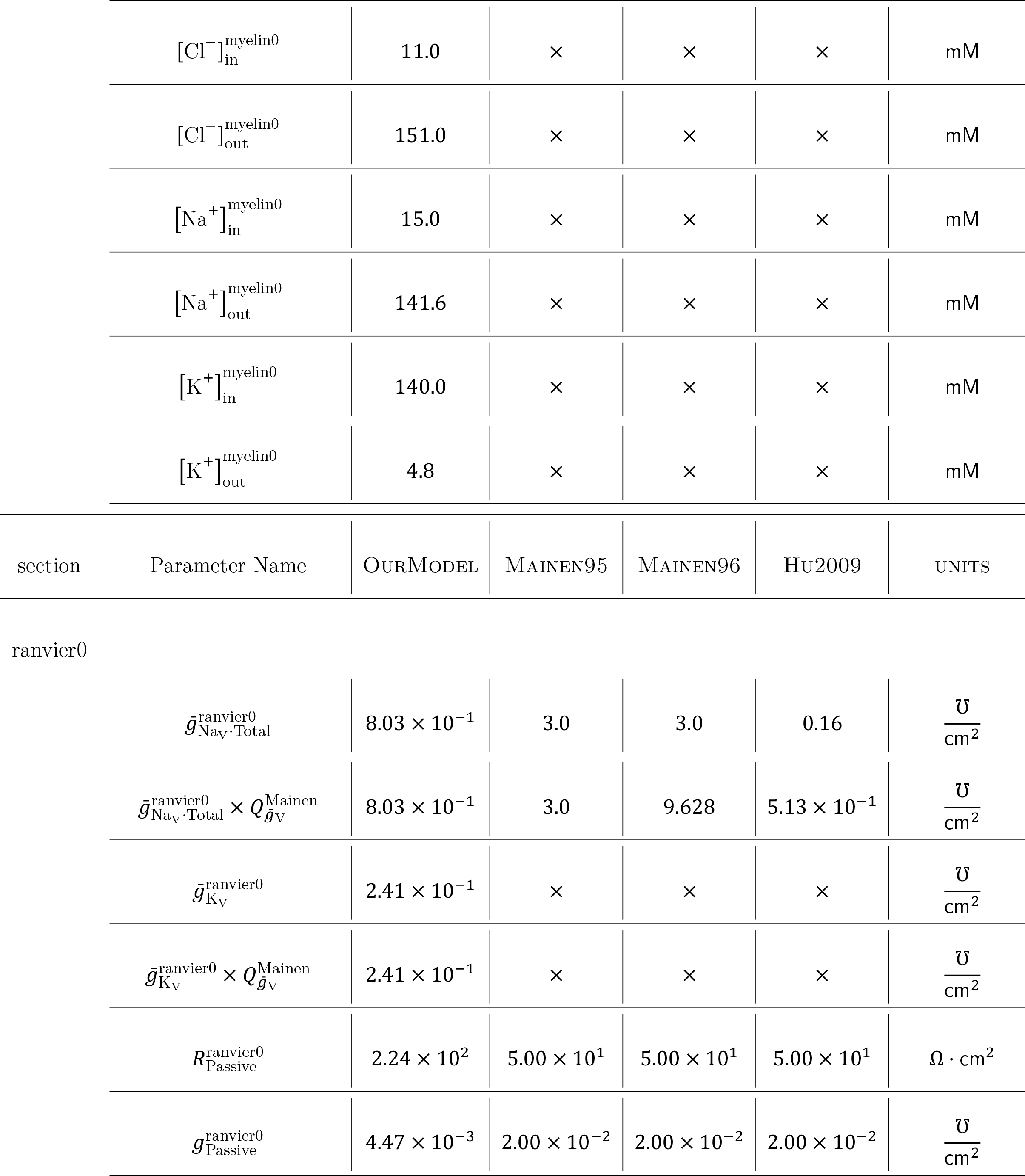
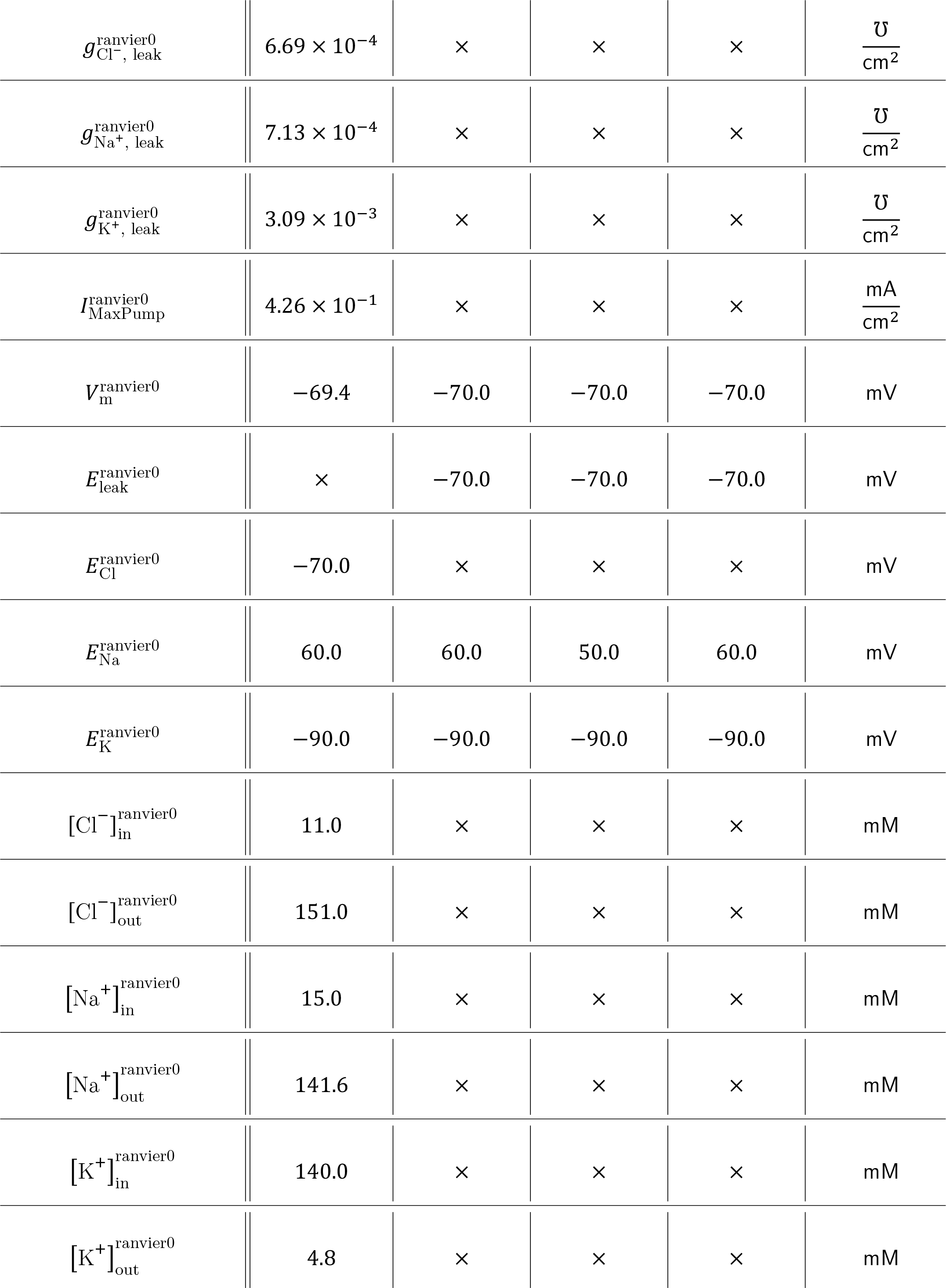
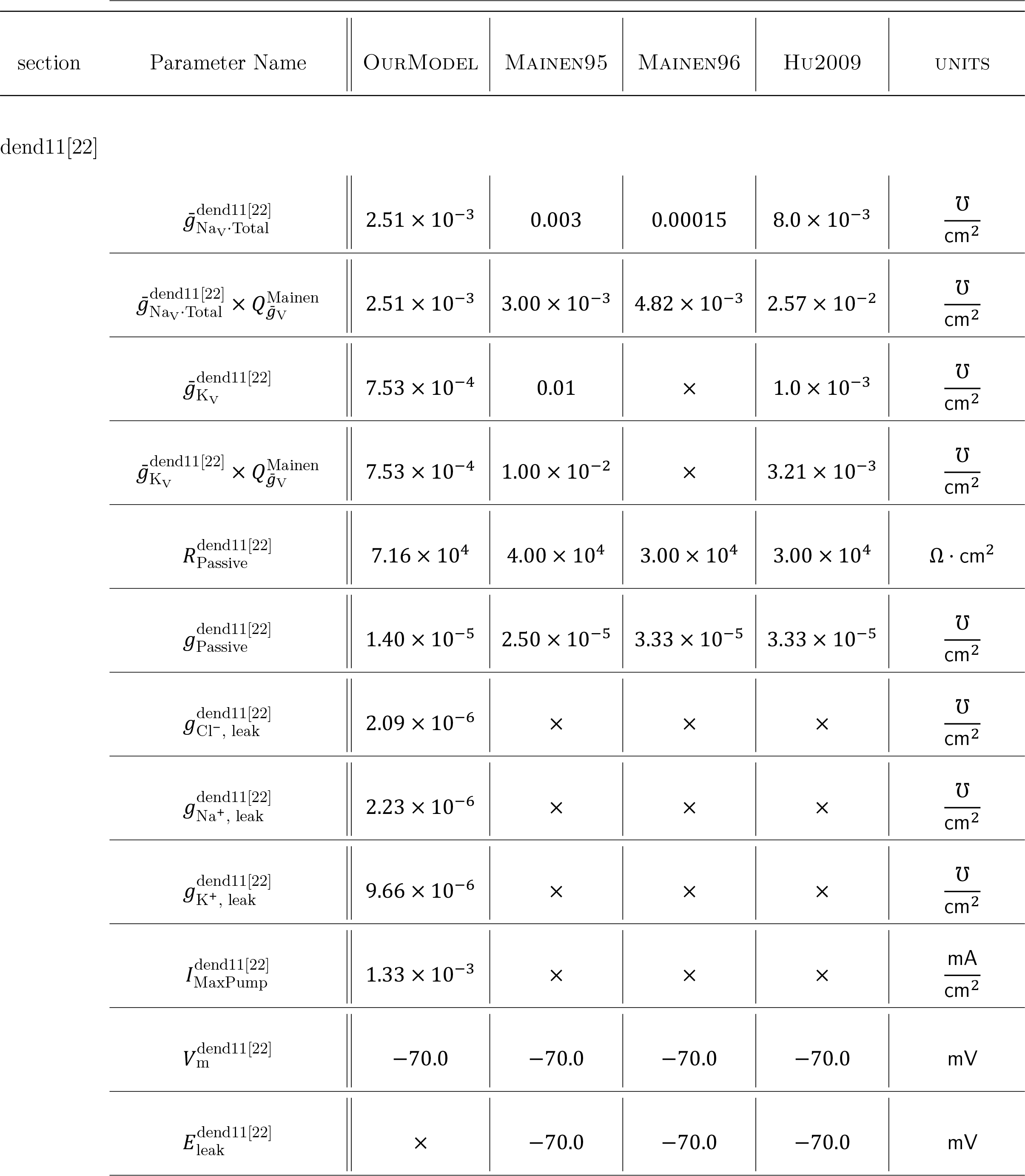
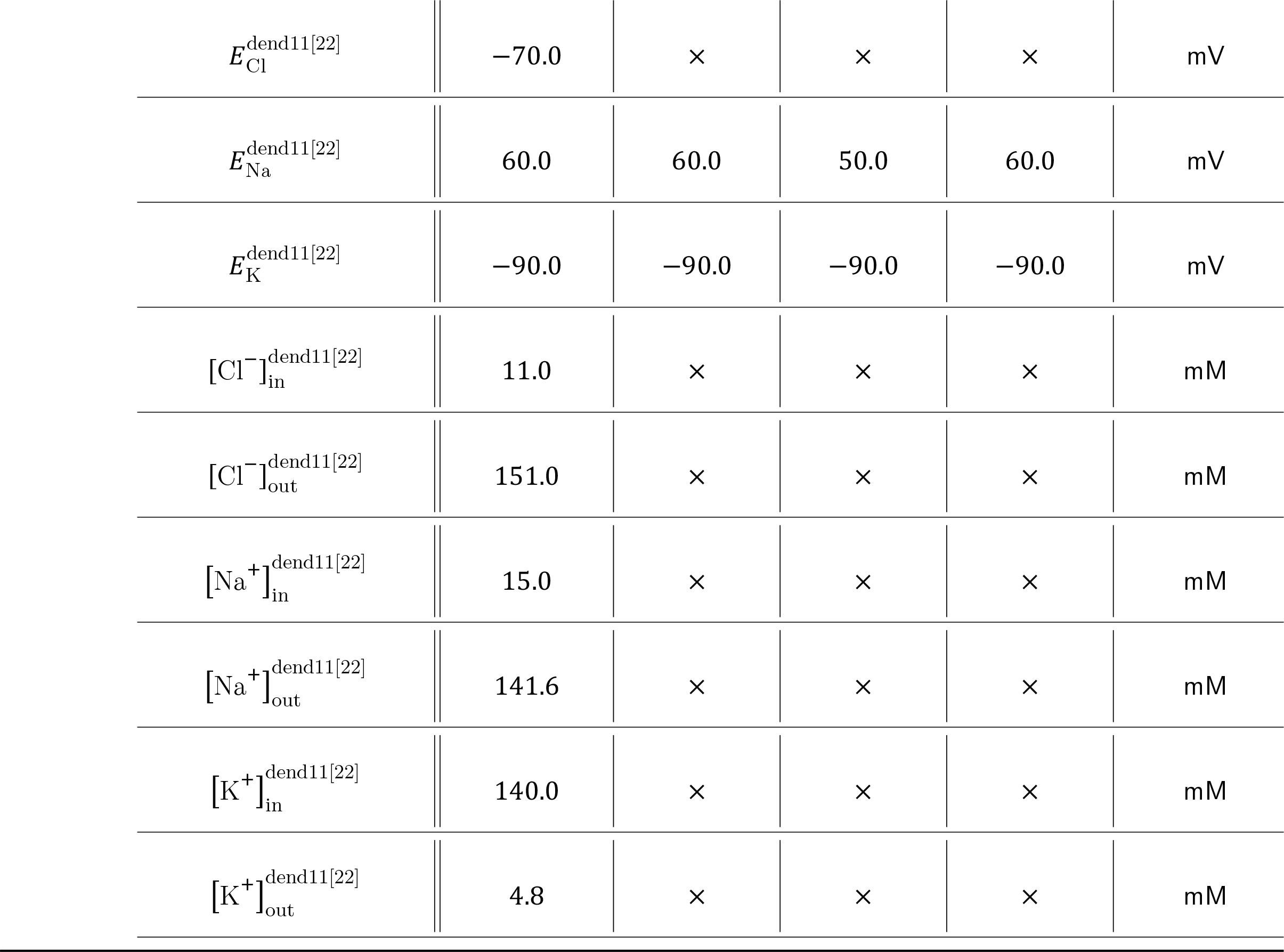

